# Convergent architecture of the transcriptome in human cancer

**DOI:** 10.1101/028415

**Authors:** Lihua Zou

## Abstract

Despite large-scale efforts to systematically map the cancer genome, little is known about how the interplay of genetic and epigenetic alternations shapes the architecture of the tumor’s transcriptome. With the goal of constructing a system-level view of the deregulated pathways in cancer cells, we systematically investigated the functional organization of the transcriptomes of 10 tumor types using data sets generated by The Cancer Genome Atlas project (TCGA). Our analysis indicates that the human cancer transcriptome is organized into well-conserved modules of co-expressed genes. In particular, our analysis identified a set of conserved gene modules with distinct cancer hallmark themes involving cell cycle regulation, angiogenesis, innate and adaptive immune response, differentiation, metabolism and regulation of protein phosphorylation. We applied a network inference approach to nominate candidate drivers of these conserved gene modules. The predicted drivers have consistent cancer-relevant functions related to the specific hallmarks and are enriched with cancer consensus genes and significantly mutated genes. We showed genetic alternations of *TP53* and other cell cycle drivers have major downstream transcriptional impact on cell cycle regulation. Collectively, our analysis provided global views of convergent transcriptome architecture of human cancer. The result of our analysis can serve as a foundation to link diverse genomic alternations to common transcriptomic features in human cancer.

## INTRODUCTION

The evolutionary history of a cancer cell has documented the accumulation of random ‘passenger’ mutations along with a few rate-limiting ‘driver’ events which confer fitness benefit to the cell. The cumulative effect of acquired driver events eventually set cancerous cells into uncontrolled state of proliferation, immortality, angiogenesis, cell death, invasion and metastasis. However, how the genetic and epigenetic effects propagate through intricate gene and biochemical circuits to cause deregulated cellular states within a cancer cell is still largely unknown. A promising approach is to decompose the oncogenic state of tumor cells directly in terms of the activity of sets of conserved biological features that has been described as the hallmarks of cancers^1^.

The comprehensive multi-dimensional genomic data provided by TCGA datasets across many tumor types offers an unprecedented opportunity to deepen our understanding of cancer biology and the discovery of new therapeutic and diagnostic targets. Gene expression analyses using unsupervised clustering approaches have been widely used in cancer research. Although informative, these methods analyzed gene level information and are prone to intrinsic noise caused by various biological and experimental factors. During the last decade, studies^2-5^ showed that gene modules consisting of co-regulated or co-expressed genes can be considered as higher-order building blocks of the global transcriptomic network and have been successfully applied to detect gene-disease association by improving the statistical power beyond the level of individual genes. Systematic data-driven approaches^6-10^ have also been developed to identify co-expressed gene modules in order to elucidate patterns of transcriptome organization across tissues and species.

We adopt the data-driven strategy to investigate the co-expression networks across 10 TCGA human cancer data sets including glioblastoma multiforme (GBM), ovarian carcinoma (OV), colon and rectal cancer (COADREAD), lung squamous cell carcinoma (LUSC), breast invasive carcinoma (BRCA), head and neck squamous cell carcinoma (HNSC), kidney renal clear cell carcinoma (KIRC), bladder urothelial carcinoma (BLCA), uterine corpus endometrioid carcinoma (UCEC) (Supplementary Table 1). Our analysis pipeline has four components: 1) construction of the co-expression network for each cancer data set and decomposition of the network into Co-expressed Gene Modules (CGMs) (Figure 1a); 2) identification of ‘conserved’ (or overlapping) CGMs across multiple tumor types (Figure 1b); 3) extraction of the core genes of conserved CGMs and classification of CGMs by cancer hallmark themes (Figure 1c); and 4) nomination of putative drivers of CGMs based on analysis of the human protein interaction network (Figure 1d).

**Figure 1.**
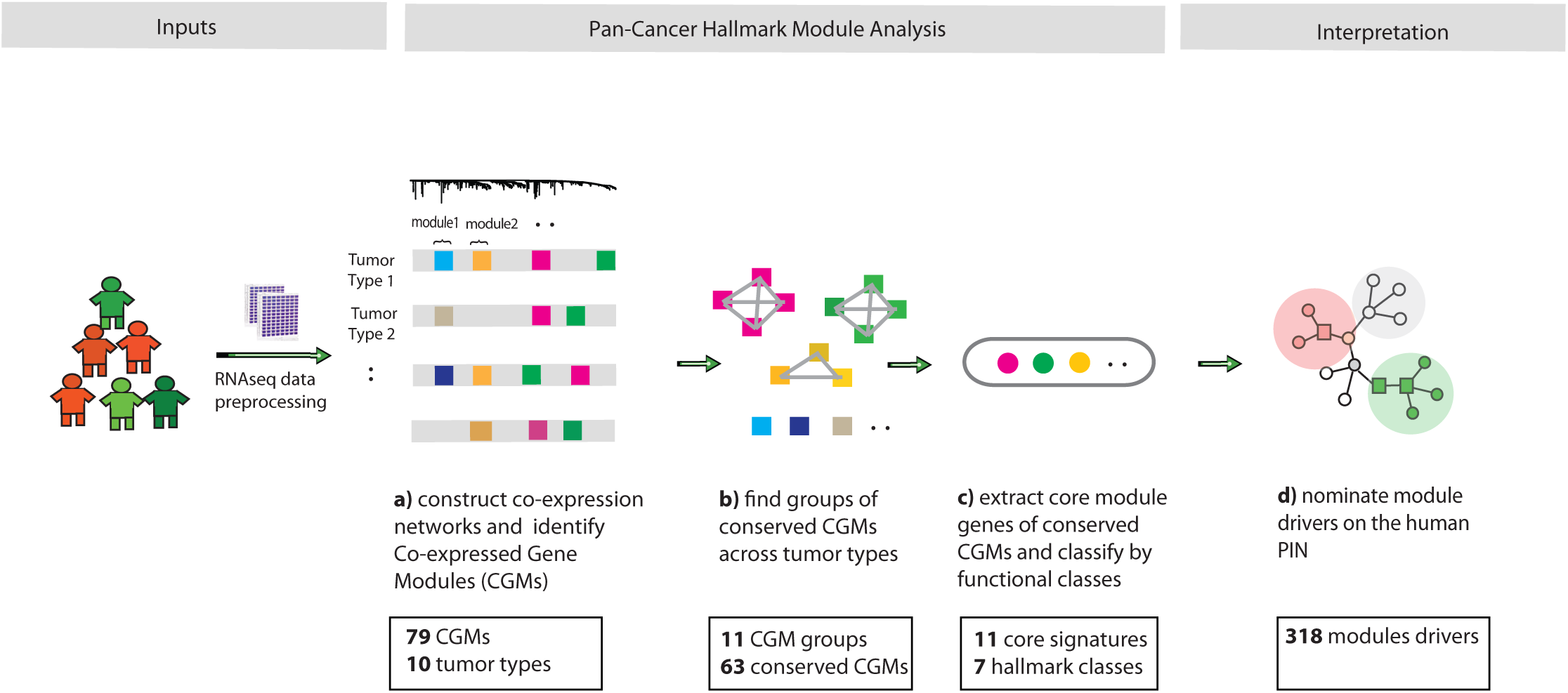
Overview of Pan-Cancer hallmark module analysis. **a**, For RNAseq data of each cancer data set, we construct the co-expression network based on the pair-wise Pearson correlation of gene expression profile. In total, 79 CGMs were identified by dynamic tree cut algorithm^12^. **b**, We enumerate all possible pairs of CGMs from any two different cancer data sets to find 11 groups of conserved CGMs across tumor types. **c**, We extract core module genes for each of the conserved CGM groups. We also classify the conserved CGMs into 7 functional categories by their enriched cancer hallmarks. **d**, We map the core module genes onto a human protein interaction network and apply NetSig to nominate module drivers (Supplementary Methods).

Overall, we identified a set of conserved cancer CGMs across multiple tissue lineages. These conserved cancer CGMs are enriched with distinct gene ontology and cancer hallmark themes. Predicted module drivers have consistent cellular functions with their corresponding hallmarks and are enriched with cancer consensus genes and significantly mutated genes.

## RESULTS

### Co-expression network analysis reveals conservation of gene modules

We followed a previously described method^9^ and constructed gene co-expression networks on the basis of Pearson correlation between the RNA-seq expression profiles for the 10 TCGA cancer data sets (Supplementary Methods). For each cancer data set, co-expressed gene modules (CGMs) were identified in two steps: First, we performed unsupervised hierarchical clustering of the gene expression profiles using the topological overlap^8,11^ as a pair-wise distance measure; and then applied the dynamic tree cut method^12^ to decompose the dendrogram at different height and merge the clusters to form stable CGMs iteratively (Supplementary Methods). We defined 12 CGMs in GBM, 5 in OV, 10 in COADREAD, 8 in LUSC, 7 in BRCA, 9 in HNSC, 11 in BLCA, 8 in KIRC, and 9 in UCEC. In total, 79 cancer CGMs were identified by this analysis.

Next we tested whether these CGMs can be validated using data from other experimental platforms and in independent data sets. We first compared the TCGA GBM expression data (syn1446212) measured by RNA-seq and by Affymetrix microarrays. We applied the same procedures to both data sets and were able to detect the majority of the CGMs defined using Affymetrix microarray when using RNA-seq data, suggesting that CGMs are reproducible across technical platforms (Supplementary Figure 1). We performed a similar comparison using 2 independently published GBM expression data sets (GSE4271, GSE13041) and reassuringly, as above, the majority of CGMs detected in the TCGA data were also detected in the external data sets demonstrating the robustness of the CGMs (Supplementary Figure 1).

To determine whether CGMs from different cancer data sets were composed of overlapping genes, we enumerated all pair-wise CGMs (n=2,756) from any two different cancer data sets and identified 509 pairs of CGMs with significant mutual overlap (Bonferroni corrected hypergeometric test, p < 0.05, Supplementary Materials). The list of overlapping pairs of CGMs is provided in Supplementary Table 2. Surprisingly, 63 out of 79 (82%) CGMs showed significant overlap with CGMs detected in at least two different tumor types (Figures 2 and 3). Taken together, our analysis suggests that the human cancer transcriptome is organized into modules of co-expressed genes that are conserved across tumor types.

**Figure 2.**
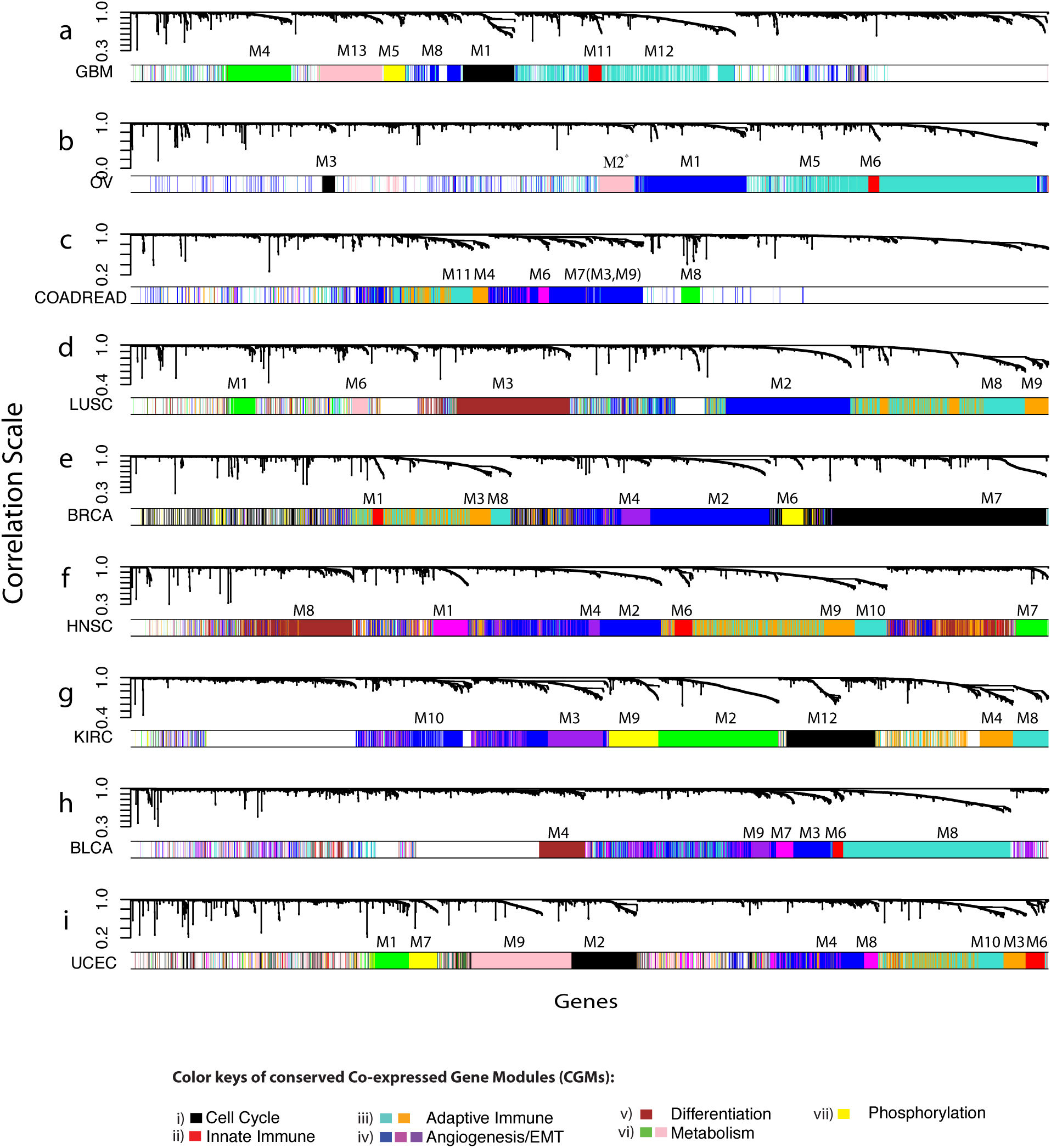
Detection of conserved co-expressed gene modules. **a-i**, Each dendrogram corresponds to the co-expression network of one of the 10 cancer data set (COAD and READ are combined into one cancer data set) produced by average linkage hierarchical clustering of genes on the basis of topological overlap^11^. The vertical scale of the dendrogram is the measure of distance i.e. 1 − ||*d*||, where *d* is Pearson correlation of a pair of expression profile. CGMs in each cancer data set are assigned unique color to distinguish different modules and numbered uniquely (e.g. M1, M2.) within the set. The M2 module of OV (indicated by ^*^) is at the borderline of modularity cutoff (Supplementary Materials) and is manually assigned to the pink group. We merged the M7 module of COADREAD with two close neighbors M3, M9 as indicated within the brackets. Conserved CGMs from different cancer data sets with significant gene overlap were assigned the same color as indicated by the horizontal bar beneath each dendrogram. For clarity, only CGMs conserved in at least 3 different tumor types are shown. The remaining non-conserved or poorly connected genes are colored white. The color key at the bottom shows the function categories of the conserved CGMs.

**Figure 3.**
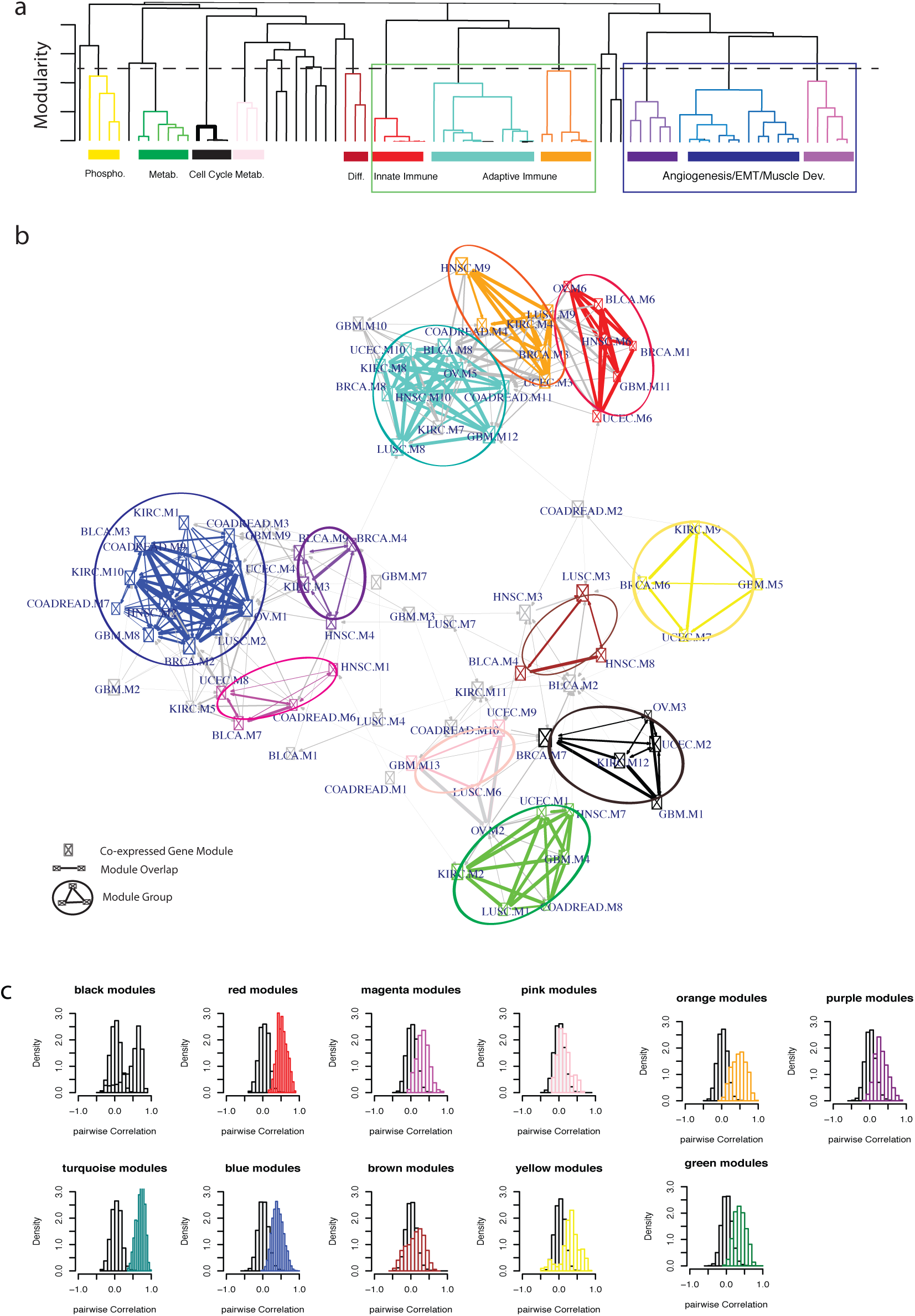
Hierarchical organization of conserved co-expressed gene modules. **a**, The leafs of the hierarchical tree represent a CGM. Groups of highly conserved CGMs are shown as distinct subtrees of the hierarchical tree corresponding to the module-module network shown below. The dotted line represents modularity threshold. **b**, Each node in the module-module network represents a CGM. Each edge represents the gene overlap between two CGMs of different tumor types. The color circles indicate the conserved CGM groups. The node color matches the leaf color of the tree. **c**, Each colored histogram shows the Pearson correlation of all possible gene pairs from one of the conserved module group. The overlapping grey histogram to the left shows the Pearson correlation of all genes pairs from a random gene set which has the same size of the conserved module group.

### Hierarchical organization of co-expressed gene modules

To better understand their global organization, we sought to define the relationships between these CGMs through the construction of a module-module network. The resulting network has 79 nodes, each represents one CGM, and 509 edges between significantly overlapping CGMs. Many module nodes appear to be closely connected on the resulting network (Figure 3b). To visualize the multi-level relationship among the CGMs, we transformed the module-module network into a hierarchical tree (Figure 3a) based on the random walk distance^13^. We partitioned the hierarchical tree into subtrees or module groups based on network modularity^14^ (modularity threshold=0.37, Supplementary Materials), a graph structure measure reflecting the enrichment of edges among nodes within a group (or cluster) relative to the nodes between the groups.

As shown in Figure 3b, our analysis revealed 11 groups of CGMs comprised of the 63 CGMs with high degree of mutual overlap (Figures 2 and 3). The 11 CGM groups have a median of 5 CGMs representing at least 3 different tumor types in each group. This implies that these CGM groups capture common or ‘conserved’ cellular processes across cancer types. The 11 CGM groups are shown as distinct subtrees on the transformed hierarchical tree of CGMs (Figure 3a). Notably, the red, turquoise, and orange CGMs fall within one branch delineated by the green square on Figure 3a which mostly involves immune response. The blue, purple, and magenta modules fall within another branch (enclosed by the blue square on Figure 3a) which primarily involves angiogenesis, epithelial-mesenchymal transformation (EMT) and muscle development. The hierarchical tree of identified gene modules suggests convergent functional organization of cellular programs of tumor independent of their tissue lineage.

To test whether genes belonging to each of the CGM groups are indeed co-expressed across their tumor types, we calculated the Pearson correlation for all gene pairs between their expression levels in the relevant tumor types. We confirmed the gene-gene correlation within the overlapping CGMs is significantly higher than expected from a random gene set of the same size (Kolmogorov-Smirnov test *p* < 2.6 × 10^−16^, Figure 3c).

### Conserved expression modules are enriched with cancer hallmarks

To assess possible biological functions of the CGMs, we calculated gene ontology enrichment for individual CGM (Figure 4, Supplementary Materials). We also extract core module genes (Supplementary Materials) for each of the 11 CGM group and calculate its functional enrichment using DAVID tools (http://david.niaid.nih.gov/david/). We classified the CGMs into 7 functional categories based on their specific enrichment of gene ontology^15^ and cancer hallmarks (Supplementary Table 3): (i) cell cycle (FDR< 2.2 × 10^−51^, black modules of Figure 2-4); (ii) innate immune response (FDR< 7 × 10^−14^, red modules of Figure 2-4); (iii) adaptive immune response (FDR< 5.6 × 10^−109^, turquoise and orange modules of Figure 2-4); (iv) differentiation (FDR< 9 × 10^−10^, brown modules of Figure 2-4); (v) angiogenesis/EMT (FDR< 2 × 10^−7^, blue, purple, and magenta modules of Figure 2-4); (vi) metabolism (FDR<0.1 green and pink modules of Figure 2-4); (vii) phosphorylation (FDR<0.3, yellow modules). We referred to these classified CGMs as “hallmark gene modules” and highlighted their core module genes below. We provide the full list of core module genes in Supplementary Table 4.

**Figure 4:**
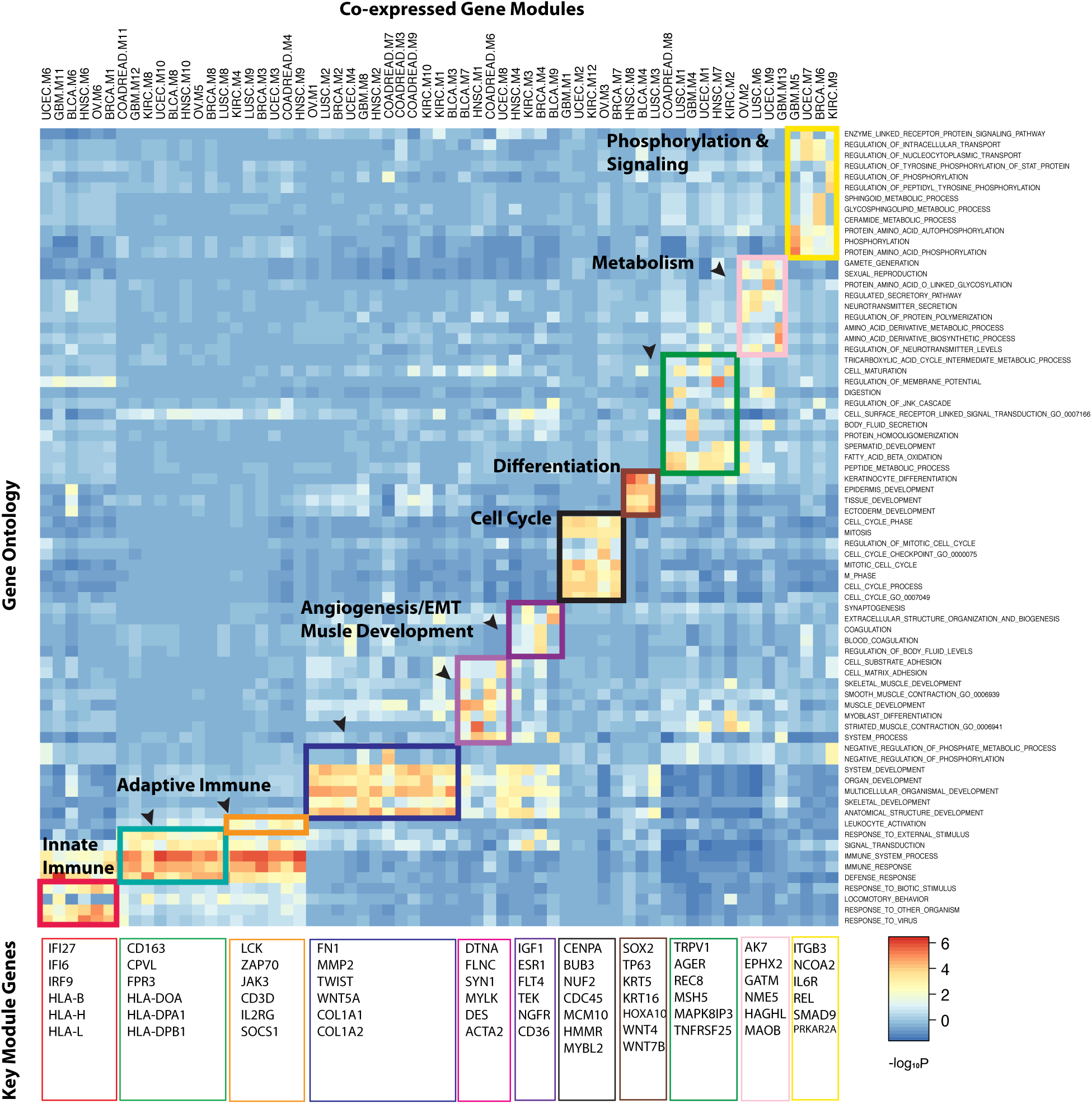
Gene ontology of co-expressed gene modules. The columns of the matrix are the conserved CGMs. Examples of core genes (indicated within the colored square) for each of the conserved CGMs are shown below the matrix. The color rectangulars along the diagonal indicate the conserved CGM groups (columns) and their most significantly associated gene ontology terms (rows). The color intensity of the matrix cell represents the level of association by one-sided hypergeometric test. Columns are ordered so the modules of the same group are together. The rows are grouped so the most significantly enriched gene ontology terms for a module group are together.

The cell cycle modules (class i, black modules of Figure 2-4) are enriched with genes involving chromosome instability (i.e. *CENPA, BUB3, NUF2*^16-18^) and DNA replication (i.e. *CDC45, MCM10*^19^). Other notable genes in this module include transcription factors *MYBL2, FOXM1, E2F2* and mitotic regulators such as *CCNB2, CDK1, AURKB, CDC25A and CDC25C.* The innate immune modules (class ii, red modules of Figure 2-4) include multiple INF-signaling and virus-response pathway genes including *IFI27, IFI6, IRF9* and several MHC class I genes including *HLA-B, HLA-H, HLA-L.* Mutations and somatic copy number alternations (SCNAs)(doi:10.7303/syn300013) in several *HLA* gene family members affect about 10% of total TCGA cancer patients (Supplementary Figure 2). The adaptive immune modules (class iii, turquoise and orange modules of Figure 2-4) are present in all 10 TCGA cancer data sets for this analysis. The core adaptive immune genes include multiple inflammatory response genes *CD163, CPVL, FPR3* and members of MHC class II genes including *HLA-DOA, HLA-DPA1, HLA-DPB1.*

The angiogenesis/EMT modules (class iv, blue modules of Figure 2-4) are present across all 10 TCGA data sets in this study. Core genes of the angiogenesis/EMT modules include *FN1, MMP2, TWIST1, and WNT5A* that are involved with cell adhesion, embryonic development, angiogenesis and TGF-beta mediated signaling^1^. Somatic mutations and SCNAs in members of angiogenesis and TGF-beta pathway are shown to affect 37% of human cancers (Supplementary Figure 2). The differentiation modules (class v, brown modules of Figure 2-4) appear to be specific to squamous carcinoma including LUSC, HNSC and BLCA. The core genes of the differentiation module contains multiple tissue differentiation genes, for example, *SOX2, TP63, KRT5, KRT16, HOXA10* and *WNT* signaling pathway genes *WNT4* and WNT7B^20,21^. *SOX2* is reported previously as lineage-survival proto-oncogene with essential roles in the differentiation and development of normal squamous cell^22^. *SOX2* is located on 3q26, a frequent amplified region in HNSC and LUSC^23^. The mutual exclusivity between *SOX2* and *TP63* mutations (Supplementary Figure 2) suggests a convergent mechanism driving the differentiation transcriptional programs among a subset of squamous carcinoma.

### Functions of imputed module drivers reflect hallmark themes

We next sought to nominate candidate drivers of the conserved CGMs. Previous studies suggest that disease genes in close proximity on the protein interaction network (PIN) will likely be functionally related^24,25^. We compiled a joint human PIN dataset from various resources including HPRD^26^, NCI-PID^27^, Cancer Cell Map from (http://cancer.cellmap.org/cellmap/), and REACTOME^28^ (Supplementary Material). For each of the CGM groups, we applied a network-based driver inference algorithm (*NetSig*, Supplementary Materials) to prioritize putative module drivers based on their proximity to the core module genes on the PIN. Figure 5 shows examples of protein subnetworks comprised of the direct protein interactions between the top candidate module drivers (red node, Figure 5) and the most connected core module genes (Supplementary Materials).

**Figure 5:**
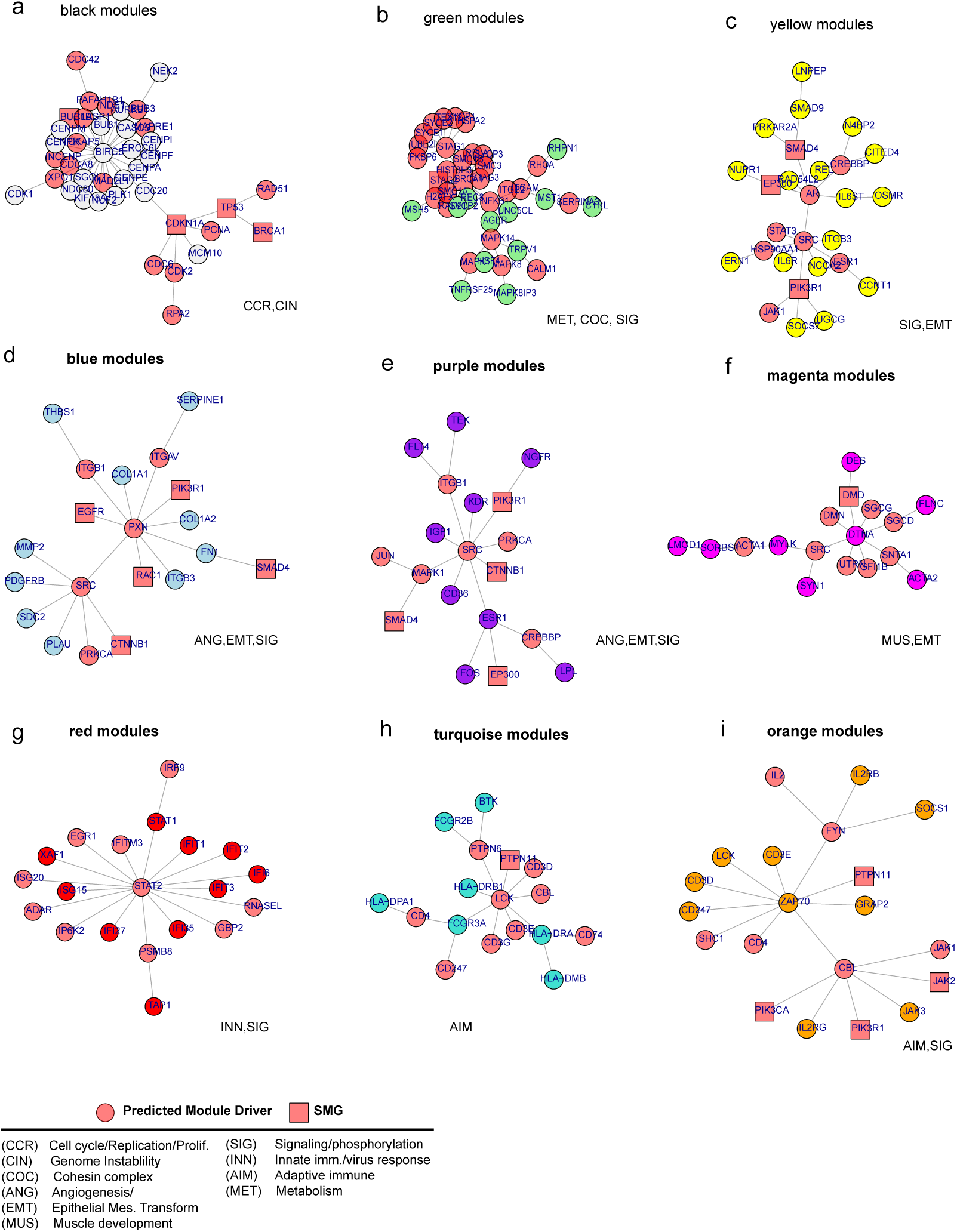
Examples of protein subnetworks connecting conserved module genes. **a-i**, The red round shaped nodes in each protein subnetwork indicate top candidate drivers of one of the conserved CGM group. The red square shaped nodes represent the predicted drivers that are also SMGs according to MutSig analysis. The edges connecting two nodes represent the human protein-protein interactions. The predominate cellular functions of the predicted module drivers are indicated next to each protein subnetwork. The functional classes of the predicted module drivers are shown at the bottom.

Our analysis identified a total of 318 candidate module drivers (doi:10.7303/syn1960770). Many of the top candidate drivers identified by *NetSig* have consistent oncogenic roles relevant to the functional classes of their CGMs. For example, top candidate drivers (red nodes, Figure 5a) of the cell cycle modules (black modules, Figure 5) include *TP53, BRCA1, CDKN1A, CDK2, CDC6, PCNA and BUB1A*, all of which have well-known roles in the regulation of cell cycle and genomic stability^1^. Notably, non-synonymous mutations and foal SCNAs of *TP53* and other cell cycle drivers are present in 55% of the human cancers (Figure 6d). This highlights the role of DNA repair and genomic instability as the key enabling hallmarks of human cancer. Tumor patients carrying genetic alternations of *TP53* and other cell cycle drivers tend to have increased number of SCNA events (especially deletions) (Figure 6b), which is in agreement with the observation made independently by Zack et al. *Submitted.* To assess the downstream transcriptional impact of the genetic alternations to the cell cycle drivers, we defined the pathway activity of a gene set in terms of the coefficient of the first principle component of the expression matrix of the pre-defined gene set^29^. Then we calculated the pathway activity for the cell cycle modules and the knowledge-based KEGG cell cycle pathway^30^. The activity profile of the cell cycle modules and the KEGG cell cycle pathway (Figure 6e) are concordant with each other and both highly up-regulated among tumor patients with genetic changes of cell cycle drivers (student t-test, *p* < 2.2 × 10^−16^). In addition, the cell cycle module genes are expressed at much higher level than adjacent non-tumor tissues across 6 TCGA tumor types with such comparison available (Figure 6f).

**Figure 6:**
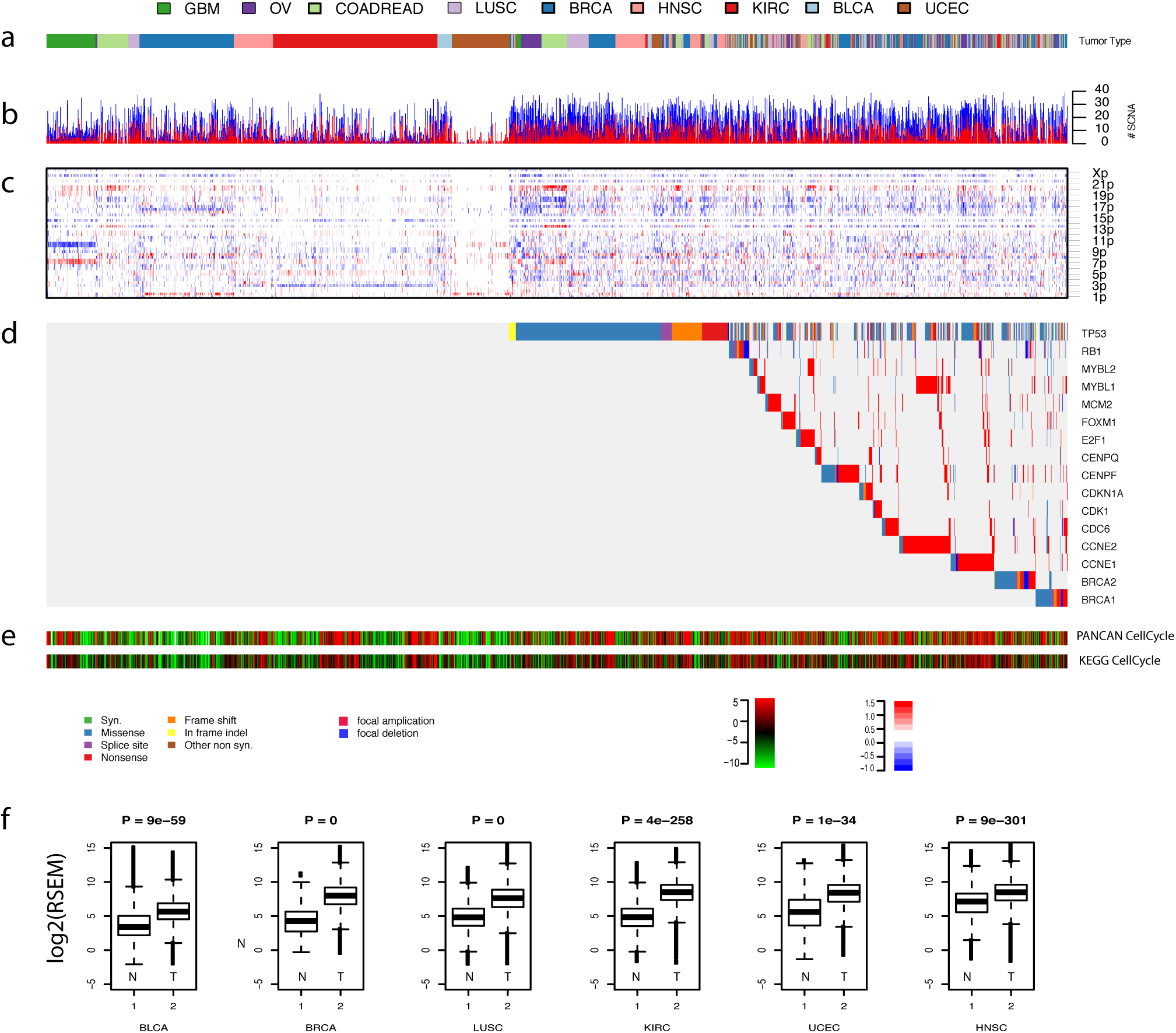
Transcriptional impact of genetic alternations of cell cycle drivers. **a**, The color keys for different tumor types used in this study. **b**, The stacked barplot shows sample-by-sample counts of broad amplification (red) and deletion events (blue) across tumor types (horizontal axis). **c**, The heatmap shows the SCNAs in each cancer sample (horizontal axis) plotted by chromosomal locations (vertical axis). **d**, The heatmap shows the somatic mutations of key cell cycle drivers sample-by-sample across tumor types. **e**, The pathway activity profile ^29^ of the core cell cycle modules and the KEGG cell cycle pathway. The color key of somatic mutations and SCNA, the scale of pathway activity and copy number value based on GISTIC2^43^ are shown below the activity profile from left to right. **f**, The boxplots show the comparison between the expression level (vertical axis) of the core cell cycle genes in adjacent normal tissue versus tumor tissue in 6 different TCGA data sets.

The protein subnetworks of the angiogenesis/EMT modules (blue and purple modules, Figure 5) share several common candidate drivers including *CTNNB1, SMAD4, SRC*, and *ITGB1*, which have known regulatory and signaling roles in TCF-beta signaling, angiogenesis and EMT^1,31,32^. Other notable candidate drivers in the subnetworks include *RAC1, EP300, EGFR* which have been shown to be frequently mutated in human cancer (doi:10.7303/syn1715784.2). The top predicted drivers of the closely related module group (magenta modules, Figure 5f and Figure 3) include several muscle development genes including *DMD, DMN, SGCD* and *SRC* among others.

The top predicted drivers of the innate immune modules (red modules, Figure 5g) include *STAT2* and *RNASEL* among others. *STAT2* is reported to mediate the innate immunity and host defense mechanism via the Type I Interferon receptor ^33^. *RNASEL* contributes in many ways to the interferon anti-virus system^34^. The top candidate drivers of the adaptive immune modules (turquoise and orange module, Figure 5) include *LCK, CD4, CBL, PTPN11* (also known as *SHP2*) with important roles in diverse immune response through the antigen recognition by the T cell receptor (TCR)^35^. The top predicted drivers of the phosphorylation modules (yellow modules, Figure 5c) are linked to members of several major signaling pathways including *AR, ESR1, PIK3R1, HSP90AA1, JAK1, STAT3, SMAD4*^1^.

The top predicted drivers of the metabolism modules (green modules, Figure 5b) include multiple members of the cohesin complex *SMC1A, SMC1B, SMC3, RAD21, STAG1 and STAG2.* Cohesin complex are frequently mutated in acute myeloid leukemia (AML)^36^ and defects in cohesin complex may underlie chromosomal instability of human cancers^37,38^. Other notable candidates of the subnetwork include *H2AFX, HIST3H3, RHOA, MAPK8* and *MAPK14. H2AFX* is frequently mutated in the Pan-Cancer data by MutSig algorithm^39^ (doi:10.7303/syn1715784.2).

We next asked whether the module drivers are enriched with cancer genes targeted by genomic alternations. We compared the predicted module drivers with the mutation drivers identified by the MutSig algorithm in the Pan-Cancer data (doi:10.7303/syn1715784.2). There are 29 out of 318 predicted modules drivers (Supplementary Table 3) overlapping with mutation drivers (hypergeometric test *p* < 8 × 10^−26^). There are 41 out of 318 module drivers (Supplementary Table 5) overlapping with the Cancer Consensus Genes (CCG)^40^^38^ (hypergeometric test *p* < 1 × 10^−20^). Overall, the result suggests that the predicted module drivers likely capture aspects of the biology and shed new light on the functional context of the cancer drivers discovered via mutation analysis.

## DISCUSSION

To our knowledge, this is the first extensive investigation of human cancer transcriptome to date. Our study defined the global transcriptome architecture of human cancer. We identified 11 groups of CGMs conserved across multiple tumor types. These conserved CGMs display distinct gene ontologies and cancer hallmark themes. The highly conserved CGMs across tumor types suggest the existence of selective force shaping the cancer transcriptome. Such conserved CGMs can be viewed as ‘building blocks’ of the cancer transcriptiome and be used as surrogate for the common cellular functions of human cancer. We noticed these conserved CGMs are significantly correlated with a large collection of published gene signatures (data not shown) although they are derived directly from the data and not dependent on any prior knowledge. As an example, we showed that the activity profile of the cell cycle modules is highly concordant with the knowledge-based KEGG cell cycle pathway (Figure 6e).

As a starting point to understand the common genetic and regulatory mechanism of these conserved CGMs, we integrated cancer transcriptome and interactome to nominated drivers for the conserved CGMs. We showed the predicted module drivers recapitulate known tumor biology and have consistent cancer-relevant functions with the hallmark themes of the conserved CGMs. Our analysis support that the cell cycle regulation and chromatin maintenance are the key system hallmarks of human cancer. Genetic alternations of *TP53* and other cell cycle drivers have major downstream transcriptional impact on the cell cycle regulation from our Pan-Cancer analysis. We anticipate future studies to elucidate mechanism leading to the deregulation of the hallmark gene modules would have direct implications for the design of new drug targets and contribute to novel pathway-focused therapeutic paradigm.

## METHODS SUMMARY

The 10 TCGA Pan-Cancer data sets used for our study are available from Sage Bionetworks (doi:10.7303/syn300013). The RNAseq molecular profiling was based on IlluminaHiSeq_RNAseqV2. The RNAseq data preprocessing was carried out using the SeqWare Pipeline project, version 0.7.0. The normalized level 3 RNAseq expression was based on RSEM^41^. Significantly recurrent SCNAs were identified using GISTIC 2.0^42,43^ on relative total copy-number data after correction for sample purity. Significantly mutated genes were identified using MutSig^39^. Co-expression network analysis was based on WGCNA package^44^.

## ACKNOWLEDGEMENTS

The authors would like to acknowledge TCGA and Sage Bionetworks for the collaborative atmosphere to generate and make the PanCan12 data freeze public available and the support of all institutions involved. We would like to thank Gad Getz, Lynda Chin, Chip Stewart for the commenting on the preliminary draft of the manuscript. We would like to thank our Broad Institute colleagues Gordon Saksena, Mike Lawrence, Andrew Cherniack and members of the Broad GDAC team for their contribution to the generation of the PanCan12 data freeze during construction of the study.

## Author Contributions

LZ conceived the study and performed the analysis and wrote the paper.

---

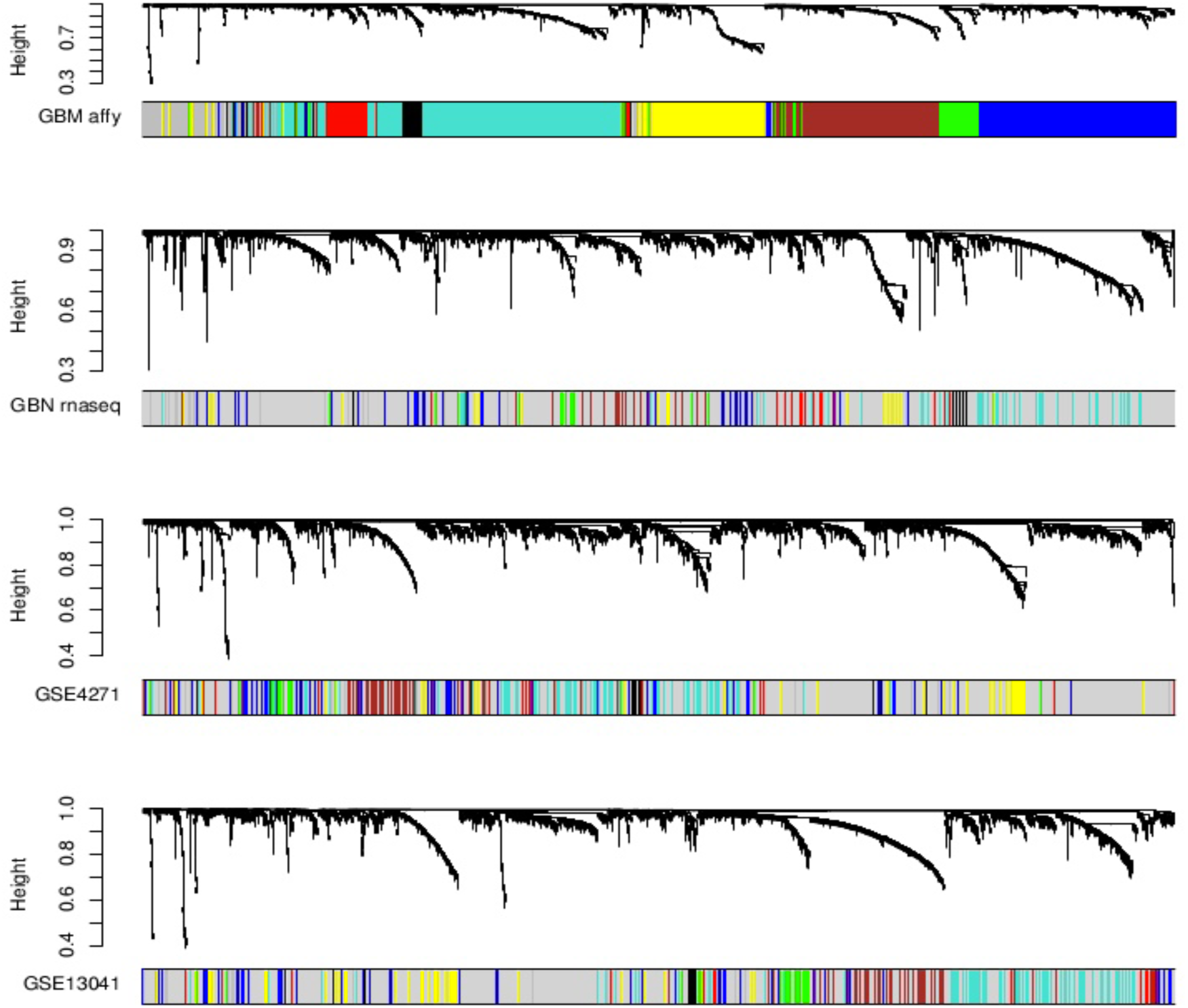

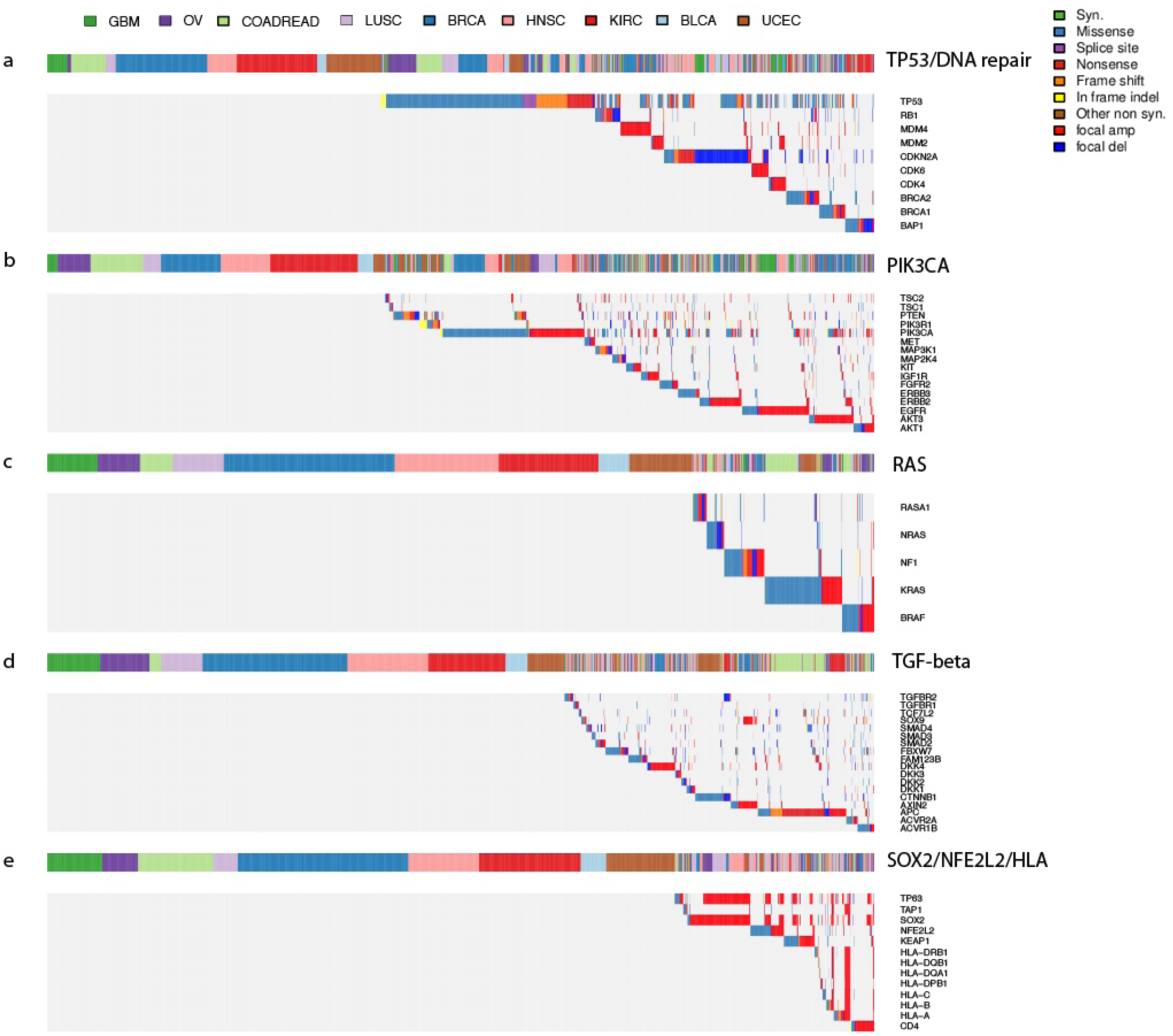

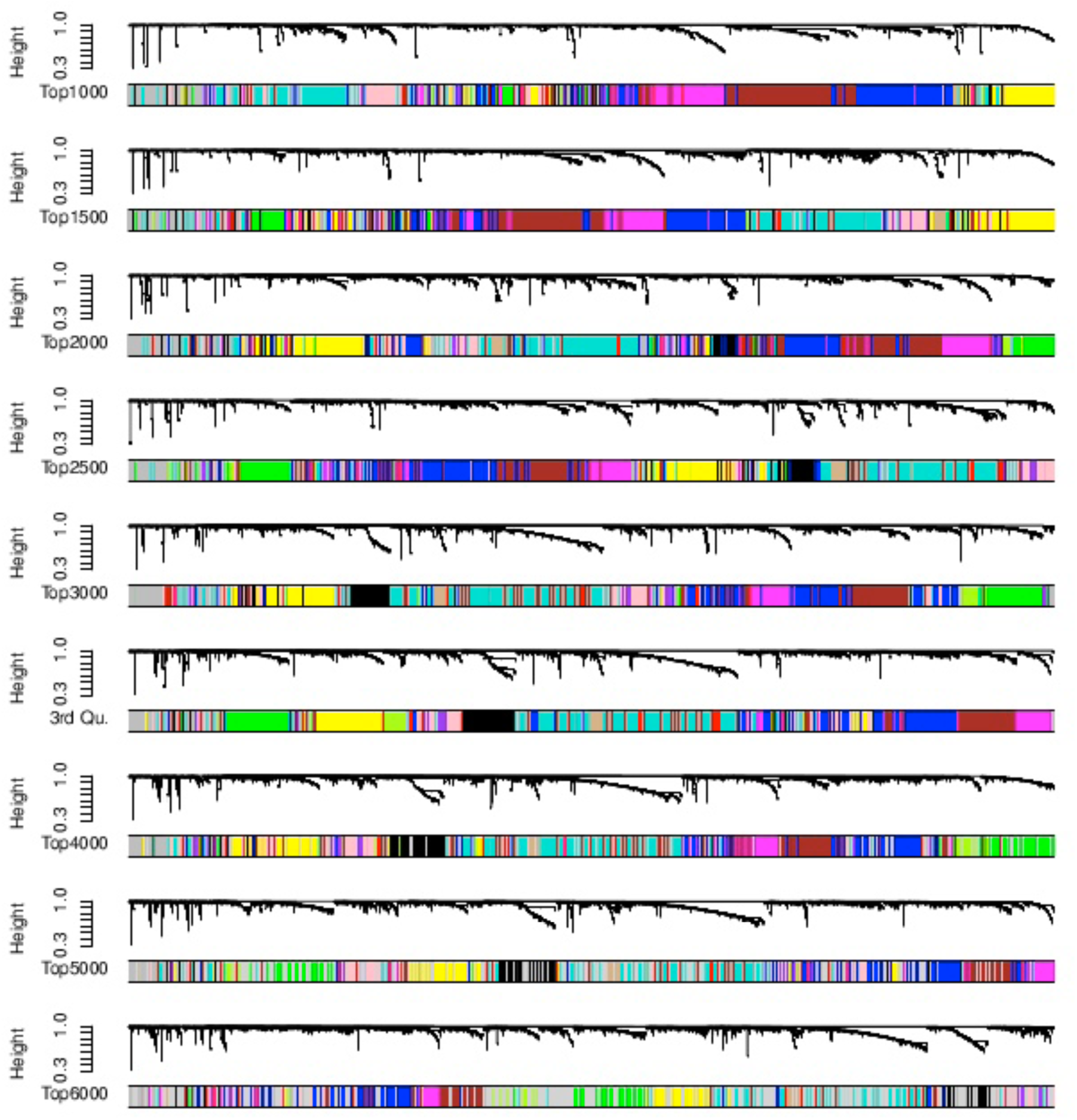

**Zou\_Suppl\_Table\_1**
| **Cancer** | **SynapseID** | **Number of Samples** | **Number of Exome** | **Number of CNV** | **Number of MIR** | **Number of RPPA** | **Number of Co-expr. Modules** | **Gene Size of Co-expr. Network** |
| --- | --- | --- | --- | --- | --- | --- | --- | --- |
| GBM | syn1461183 | 154 | 563 | 112 | 491 | 214 | 12 | 3526 |
| OV | syn1461171 | 262 | 559 | 582 | 568 | 332 | 5 | 3487 |
| COADREAD | syn1461155<br>and<br>syn1461175 | 264 | 575 | 350 | 551 | 399 | 10 | 2712 |
| LUSC | syn1461168 | 220 | 343 | 226 | 332 | 195 | 8 | 3499 |
| BRCA | syn1461151 | 841 | 866 | 575 | 868 | 408 | 7 | 3268 |
| HNSC | syn1461156 | 303 | 306 | 310 | 309 | 212 | 9 | 3394 |
| KIRC | syn1461159 | 469 | 493 | 283 | 480 | 453 | 11 | 3363 |
| BLCA | syn1461149 | 96 | 135 | 138 | 124 | 54 | 8 | 3380 |
| UCEC | syn1461180 | 333 | 492 | 383 | 485 | 200 | 9 | 3251 |

Zou\_Suppl\_Table\_2
| Module.1 | Module.2 | Adjusted.P-value |
| --- | --- | --- |
| GBM.black | OV.green | 3.60E-38 |
| GBM.black | BRCA.turquoise | 2.87E-31 |
| GBM.black | KIRC.yellow | 3.91E-54 |
| GBM.black | UCEC.blue | 1.81E-25 |
| GBM.blue | LUSC.blue | 0.001955469 |
| GBM.blue | BRCA.blue | 0.00028959 |
| GBM.blue | HNSC.blue | 8.68E-07 |
| GBM.blue | KIRC.red | 0.02992095 |
| GBM.blue | UCEC.red | 0.002824684 |
| GBM.brown | COADREAD.turquoise | 0.02830111 |
| GBM.brown | HNSC.brown | 0.00147624 |
| GBM.brown | KIRC.brown | 0.004390626 |
| GBM.brown | BLCA.yellow | 2.06E-05 |
| GBM.green | OV.brown | 1.45E-11 |
| GBM.green | COADREAD.purple | 6.96E-46 |
| GBM.green | LUSC.black | 1.09E-63 |
| GBM.green | BRCA.turquoise | 0.002820383 |
| GBM.green | HNSC.pink | 4.29E-50 |
| GBM.green | KIRC.blue | 1.23E-88 |
| GBM.green | UCEC.black | 1.13E-61 |
| GBM.greenyellow | BRCA.red | 2.40E-18 |
| GBM.greenyellow | KIRC.purple | 1.05E-29 |
| GBM.greenyellow | UCEC.pink | 9.97E-33 |
| GBM.magenta | LUSC.green | 0.0136676 |
| GBM.magenta | BRCA.green | 0.006473287 |
| GBM.magenta | KIRC.brown | 0.01165685 |
| GBM.pink | OV.blue | 3.41E-34 |
| GBM.pink | COADREAD.brown | 1.94E-08 |
| GBM.pink | LUSC.blue | 3.51E-34 |
| GBM.pink | BRCA.blue | 9.83E-31 |
| GBM.pink | HNSC.blue | 2.54E-22 |
| GBM.pink | KIRC.red | 3.92E-25 |
| GBM.pink | BLCA.brown | 3.03E-14 |
| GBM.pink | UCEC.green | 2.36E-22 |
| GBM.purple | OV.blue | 2.68E-07 |
| GBM.purple | COADREAD.brown | 8.20E-06 |
| GBM.purple | LUSC.blue | 8.82E-05 |
| GBM.purple | BRCA.green | 0.000226317 |
| GBM.purple | HNSC.blue | 0.000124524 |
| GBM.purple | KIRC.brown | 4.09E-12 |
| GBM.purple | BLCA.yellow | 0.02121653 |
| GBM.purple | UCEC.green | 5.95E-08 |
| GBM.red | OV.turquoise | 8.97E-13 |
| GBM.red | COADREAD.green | 9.66E-06 |
| GBM.red | LUSC.turquoise | 5.50E-11 |
| GBM.red | BRCA.yellow | 5.99E-14 |
| GBM.red | HNSC.yellow | 3.92E-06 |
| GBM.red | KIRC.pink | 1.10E-20 |
| GBM.red | BLCA.turquoise | 7.33E-12 |
| GBM.red | UCEC.yellow | 3.84E-13 |
| GBM.tan | OV.yellow | 2.51E-40 |
| GBM.tan | COADREAD.yellow | 9.95E-14 |
| GBM.tan | LUSC.yellow | 3.41E-31 |
| GBM.tan | BRCA.black | 5.00E-45 |
| GBM.tan | HNSC.magenta | 6.00E-49 |
| GBM.tan | KIRC.green | 2.85E-16 |
| GBM.tan | BLCA.pink | 4.51E-36 |
| GBM.tan | UCEC.magenta | 1.68E-60 |
| GBM.turquoise | OV.turquoise | 9.89E-63 |
| GBM.turquoise | COADREAD.yellow | 2.50E-22 |
| GBM.turquoise | LUSC.turquoise | 4.56E-37 |
| GBM.turquoise | BRCA.yellow | 3.25E-35 |
| GBM.turquoise | HNSC.yellow | 1.93E-51 |
| GBM.turquoise | KIRC.pink | 3.10E-26 |
| GBM.turquoise | BLCA.turquoise | 1.72E-39 |
| GBM.turquoise | UCEC.yellow | 9.09E-38 |
| GBM.yellow | OV.brown | 5.97E-48 |
| GBM.yellow | COADREAD.magenta | 0.00554752 |
| GBM.yellow | LUSC.pink | 7.30E-31 |
| GBM.yellow | BRCA.turquoise | 2.05E-05 |
| GBM.yellow | KIRC.turquoise | 6.07E-06 |
| GBM.yellow | UCEC.turquoise | 5.97E-23 |
| OV.blue | GBM.pink | 1.39E-29 |
| OV.blue | COADREAD.brown | 5.15E-24 |
| OV.blue | LUSC.blue | 8.16E-140 |
| OV.blue | BRCA.blue | 1.55E-109 |
| OV.blue | HNSC.blue | 4.75E-77 |
| OV.blue | KIRC.red | 4.62E-43 |
| OV.blue | BLCA.brown | 5.69E-73 |
| OV.blue | UCEC.green | 2.74E-90 |
| OV.brown | GBM.yellow | 2.77E-47 |
| OV.brown | COADREAD.purple | 5.77E-14 |
| OV.brown | LUSC.pink | 3.09E-62 |
| OV.brown | BRCA.turquoise | 3.08E-13 |
| OV.brown | HNSC.pink | 3.98E-19 |
| OV.brown | KIRC.blue | 4.73E-15 |
| OV.brown | UCEC.turquoise | 3.93E-49 |
| OV.green | GBM.black | 1.52E-34 |
| OV.green | BRCA.turquoise | 1.31E-14 |
| OV.green | KIRC.yellow | 3.13E-19 |
| OV.green | BLCA.blue | 0.0086865 |
| OV.green | UCEC.blue | 1.14E-24 |
| OV.turquoise | GBM.turquoise | 2.60E-71 |
| OV.turquoise | COADREAD.yellow | 1.46E-37 |
| OV.turquoise | LUSC.turquoise | 3.91E-49 |
| OV.turquoise | BRCA.yellow | 2.19E-71 |
| OV.turquoise | HNSC.yellow | 1.05E-55 |
| OV.turquoise | KIRC.pink | 1.65E-56 |
| OV.turquoise | BLCA.turquoise | 3.06E-75 |
| OV.turquoise | UCEC.yellow | 2.07E-71 |
| OV.yellow | GBM.tan | 2.51E-45 |
| OV.yellow | COADREAD.yellow | 5.45E-10 |
| OV.yellow | LUSC.yellow | 3.97E-19 |
| OV.yellow | BRCA.black | 1.53E-57 |
| OV.yellow | HNSC.magenta | 2.96E-40 |
| OV.yellow | KIRC.green | 7.32E-08 |
| OV.yellow | BLCA.pink | 2.47E-40 |
| OV.yellow | UCEC.magenta | 2.47E-43 |
| COADREAD.black | HNSC.black | 0.007794982 |
| COADREAD.black | KIRC.blue | 0.03646471 |
| COADREAD.black | BLCA.green | 0.002211642 |
| COADREAD.blue | GBM.turquoise | 0.00457559 |
| COADREAD.blue | LUSC.brown | 0.01203316 |
| COADREAD.blue | BRCA.red | 0.006733661 |
| COADREAD.blue | HNSC.brown | 0.001154678 |
| COADREAD.blue | KIRC.purple | 0.001713155 |
| COADREAD.blue | UCEC.magenta | 0.000212338 |
| COADREAD.brown | GBM.pink | 2.23E-09 |
| COADREAD.brown | OV.blue | 9.77E-31 |
| COADREAD.brown | LUSC.blue | 2.15E-30 |
| COADREAD.brown | BRCA.blue | 9.80E-20 |
| COADREAD.brown | HNSC.blue | 6.42E-26 |
| COADREAD.brown | KIRC.red | 1.69E-08 |
| COADREAD.brown | BLCA.brown | 1.77E-23 |
| COADREAD.brown | UCEC.green | 9.80E-12 |
| COADREAD.green | GBM.turquoise | 5.53E-09 |
| COADREAD.green | OV.turquoise | 3.01E-32 |
| COADREAD.green | LUSC.yellow | 2.40E-30 |
| COADREAD.green | BRCA.brown | 1.70E-40 |
| COADREAD.green | HNSC.turquoise | 3.86E-18 |
| COADREAD.green | KIRC.green | 3.98E-26 |
| COADREAD.green | BLCA.turquoise | 7.34E-30 |
| COADREAD.green | UCEC.brown | 2.15E-22 |
| COADREAD.magenta | OV.blue | 1.20E-11 |
| COADREAD.magenta | LUSC.blue | 1.61E-06 |
| COADREAD.magenta | BRCA.blue | 3.34E-08 |
| COADREAD.magenta | HNSC.green | 3.29E-11 |
| COADREAD.magenta | KIRC.greenyellow | 2.01E-17 |
| COADREAD.magenta | BLCA.red | 1.89E-33 |
| COADREAD.magenta | UCEC.red | 8.68E-26 |
| COADREAD.pink | GBM.pink | 1.12E-06 |
| COADREAD.pink | OV.blue | 3.07E-12 |
| COADREAD.pink | LUSC.blue | 2.95E-19 |
| COADREAD.pink | BRCA.blue | 4.96E-17 |
| COADREAD.pink | HNSC.blue | 3.96E-17 |
| COADREAD.pink | KIRC.red | 8.40E-14 |
| COADREAD.pink | BLCA.brown | 3.28E-12 |
| COADREAD.pink | UCEC.green | 1.84E-10 |
| COADREAD.purple | GBM.green | 1.20E-36 |
| COADREAD.purple | OV.brown | 1.16E-08 |
| COADREAD.purple | LUSC.black | 1.08E-30 |
| COADREAD.purple | HNSC.pink | 5.64E-35 |
| COADREAD.purple | KIRC.blue | 9.05E-40 |
| COADREAD.purple | UCEC.black | 2.63E-38 |
| COADREAD.red | GBM.pink | 1.57E-06 |
| COADREAD.red | OV.blue | 1.89E-13 |
| COADREAD.red | LUSC.blue | 1.36E-13 |
| COADREAD.red | BRCA.blue | 1.17E-21 |
| COADREAD.red | HNSC.blue | 5.52E-20 |
| COADREAD.red | KIRC.red | 1.31E-07 |
| COADREAD.red | BLCA.brown | 8.38E-12 |
| COADREAD.red | UCEC.green | 3.40E-11 |
| COADREAD.turquoise | BRCA.turquoise | 7.55E-08 |
| COADREAD.turquoise | KIRC.turquoise | 1.76E-06 |
| COADREAD.turquoise | BLCA.green | 0.002901911 |
| COADREAD.turquoise | UCEC.turquoise | 0.01069324 |
| COADREAD.yellow | GBM.turquoise | 1.67E-20 |
| COADREAD.yellow | OV.turquoise | 1.12E-26 |
| COADREAD.yellow | LUSC.turquoise | 1.91E-19 |
| COADREAD.yellow | BRCA.yellow | 3.15E-31 |
| COADREAD.yellow | HNSC.yellow | 1.10E-24 |
| COADREAD.yellow | KIRC.pink | 6.02E-25 |
| COADREAD.yellow | BLCA.turquoise | 4.03E-31 |
| COADREAD.yellow | UCEC.yellow | 1.08E-33 |
| LUSC.black | GBM.green | 3.12E-52 |
| LUSC.black | OV.brown | 1.19E-08 |
| LUSC.black | COADREAD.purple | 1.80E-28 |
| LUSC.black | HNSC.pink | 1.25E-42 |
| LUSC.black | KIRC.blue | 3.82E-59 |
| LUSC.black | UCEC.black | 8.91E-50 |
| LUSC.blue | GBM.pink | 2.08E-29 |
| LUSC.blue | OV.blue | 1.21E-146 |
| LUSC.blue | COADREAD.brown | 2.03E-28 |
| LUSC.blue | BRCA.blue | 4.53E-148 |
| LUSC.blue | HNSC.blue | 3.68E-159 |
| LUSC.blue | KIRC.red | 4.83E-60 |
| LUSC.blue | BLCA.brown | 2.52E-95 |
| LUSC.blue | UCEC.green | 1.76E-69 |
| LUSC.brown | HNSC.red | 1.79E-18 |
| LUSC.brown | BLCA.green | 2.14E-28 |
| LUSC.green | KIRC.turquoise | 0.004866037 |
| LUSC.green | BLCA.black | 2.00E-09 |
| LUSC.pink | GBM.yellow | 2.10E-31 |
| LUSC.pink | OV.brown | 4.44E-56 |
| LUSC.pink | BRCA.turquoise | 2.90E-09 |
| LUSC.pink | HNSC.pink | 0.01970745 |
| LUSC.pink | KIRC.turquoise | 0.0251461 |
| LUSC.pink | UCEC.turquoise | 1.01E-31 |
| LUSC.red | BRCA.green | 0.01869355 |
| LUSC.red | HNSC.green | 0.000642402 |
| LUSC.red | KIRC.turquoise | 0.001324934 |
| LUSC.red | BLCA.blue | 0.00032609 |
| LUSC.turquoise | GBM.turquoise | 7.65E-47 |
| LUSC.turquoise | OV.turquoise | 1.66E-60 |
| LUSC.turquoise | COADREAD.yellow | 7.80E-28 |
| LUSC.turquoise | BRCA.yellow | 1.29E-76 |
| LUSC.turquoise | HNSC.yellow | 5.00E-110 |
| LUSC.turquoise | KIRC.pink | 7.22E-75 |
| LUSC.turquoise | BLCA.turquoise | 3.52E-59 |
| LUSC.turquoise | UCEC.yellow | 7.86E-59 |
| LUSC.yellow | GBM.tan | 2.28E-34 |
| LUSC.yellow | OV.turquoise | 4.28E-21 |
| LUSC.yellow | COADREAD.green | 5.21E-36 |
| LUSC.yellow | BRCA.brown | 1.74E-62 |
| LUSC.yellow | HNSC.turquoise | 9.20E-62 |
| LUSC.yellow | KIRC.green | 1.47E-112 |
| LUSC.yellow | BLCA.turquoise | 2.11E-23 |
| LUSC.yellow | UCEC.brown | 2.24E-48 |
| BRCA.black | GBM.tan | 5.95E-50 |
| BRCA.black | OV.yellow | 2.29E-56 |
| BRCA.black | COADREAD.yellow | 1.98E-09 |
| BRCA.black | LUSC.yellow | 3.27E-23 |
| BRCA.black | HNSC.magenta | 9.05E-43 |
| BRCA.black | KIRC.green | 1.28E-11 |
| BRCA.black | BLCA.pink | 1.35E-41 |
| BRCA.black | UCEC.magenta | 5.76E-55 |
| BRCA.blue | GBM.pink | 8.15E-29 |
| BRCA.blue | OV.blue | 5.32E-118 |
| BRCA.blue | COADREAD.brown | 7.69E-21 |
| BRCA.blue | LUSC.blue | 1.21E-145 |
| BRCA.blue | HNSC.blue | 4.59E-143 |
| BRCA.blue | KIRC.red | 1.05E-58 |
| BRCA.blue | BLCA.brown | 6.51E-90 |
| BRCA.blue | UCEC.green | 3.17E-60 |
| BRCA.brown | GBM.turquoise | 1.80E-16 |
| BRCA.brown | OV.turquoise | 1.35E-32 |
| BRCA.brown | COADREAD.green | 2.80E-44 |
| BRCA.brown | LUSC.yellow | 4.60E-59 |
| BRCA.brown | HNSC.turquoise | 7.13E-54 |
| BRCA.brown | KIRC.green | 1.77E-67 |
| BRCA.brown | BLCA.turquoise | 5.43E-36 |
| BRCA.brown | UCEC.brown | 3.56E-58 |
| BRCA.green | GBM.purple | 0.001997649 |
| BRCA.green | OV.blue | 2.31E-12 |
| BRCA.green | COADREAD.brown | 6.27E-06 |
| BRCA.green | LUSC.turquoise | 0.003788179 |
| BRCA.green | HNSC.green | 3.95E-20 |
| BRCA.green | KIRC.brown | 2.89E-33 |
| BRCA.green | BLCA.yellow | 9.53E-21 |
| BRCA.green | UCEC.green | 2.13E-09 |
| BRCA.red | COADREAD.blue | 1.76E-05 |
| BRCA.red | LUSC.brown | 0.01183595 |
| BRCA.red | KIRC.purple | 2.86E-42 |
| BRCA.red | BLCA.blue | 0.00402368 |
| BRCA.red | UCEC.pink | 4.11E-32 |
| BRCA.turquoise | GBM.black | 4.85E-42 |
| BRCA.turquoise | OV.green | 3.30E-15 |
| BRCA.turquoise | COADREAD.turquoise | 2.97E-05 |
| BRCA.turquoise | HNSC.brown | 3.97E-06 |
| BRCA.turquoise | KIRC.yellow | 2.36E-35 |
| BRCA.turquoise | BLCA.blue | 7.58E-16 |
| BRCA.turquoise | UCEC.blue | 1.70E-27 |
| BRCA.yellow | GBM.turquoise | 8.66E-39 |
| BRCA.yellow | OV.turquoise | 2.09E-68 |
| BRCA.yellow | COADREAD.yellow | 1.91E-38 |
| BRCA.yellow | LUSC.turquoise | 1.01E-75 |
| BRCA.yellow | HNSC.yellow | 2.95E-80 |
| BRCA.yellow | KIRC.pink | 1.02E-82 |
| BRCA.yellow | BLCA.turquoise | 3.71E-67 |
| BRCA.yellow | UCEC.yellow | 4.56E-86 |
| HNSC.black | COADREAD.magenta | 1.17E-05 |
| HNSC.black | BRCA.green | 0.004042421 |
| HNSC.black | BLCA.red | 3.14E-11 |
| HNSC.black | UCEC.red | 1.02E-08 |
| HNSC.blue | GBM.pink | 7.18E-23 |
| HNSC.blue | OV.blue | 8.20E-83 |
| HNSC.blue | COADREAD.brown | 9.67E-23 |
| HNSC.blue | LUSC.blue | 8.04E-159 |
| HNSC.blue | BRCA.blue | 1.01E-138 |
| HNSC.blue | KIRC.red | 3.21E-49 |
| HNSC.blue | BLCA.brown | 5.41E-98 |
| HNSC.blue | UCEC.green | 5.20E-58 |
| HNSC.brown | GBM.brown | 0.003436834 |
| HNSC.brown | LUSC.brown | 8.58E-20 |
| HNSC.brown | BRCA.turquoise | 0.000143654 |
| HNSC.brown | BLCA.blue | 9.45E-07 |
| HNSC.brown | UCEC.turquoise | 4.01E-06 |
| HNSC.green | GBM.purple | 0.008049011 |
| HNSC.green | OV.blue | 7.14E-11 |
| HNSC.green | COADREAD.magenta | 8.67E-16 |
| HNSC.green | BRCA.green | 2.18E-22 |
| HNSC.green | KIRC.brown | 4.54E-09 |
| HNSC.green | BLCA.yellow | 1.03E-19 |
| HNSC.green | UCEC.green | 6.80E-09 |
| HNSC.magenta | GBM.tan | 3.15E-52 |
| HNSC.magenta | OV.yellow | 6.50E-41 |
| HNSC.magenta | COADREAD.yellow | 1.19E-09 |
| HNSC.magenta | LUSC.yellow | 7.11E-17 |
| HNSC.magenta | BRCA.black | 1.22E-43 |
| HNSC.magenta | KIRC.green | 1.20E-09 |
| HNSC.magenta | BLCA.pink | 4.48E-53 |
| HNSC.magenta | UCEC.magenta | 8.20E-59 |
| HNSC.pink | GBM.green | 2.17E-47 |
| HNSC.pink | OV.brown | 2.72E-14 |
| HNSC.pink | COADREAD.purple | 5.69E-35 |
| HNSC.pink | LUSC.black | 2.45E-43 |
| HNSC.pink | KIRC.blue | 3.49E-63 |
| HNSC.pink | BLCA.blue | 0.002532541 |
| HNSC.pink | UCEC.black | 1.84E-37 |
| HNSC.red | LUSC.brown | 4.40E-22 |
| HNSC.red | KIRC.turquoise | 9.54E-05 |
| HNSC.red | BLCA.green | 3.78E-45 |
| HNSC.turquoise | GBM.red | 4.98E-12 |
| HNSC.turquoise | OV.turquoise | 8.27E-21 |
| HNSC.turquoise | COADREAD.green | 9.31E-29 |
| HNSC.turquoise | LUSC.yellow | 3.40E-61 |
| HNSC.turquoise | BRCA.brown | 4.85E-59 |
| HNSC.turquoise | KIRC.green | 4.40E-49 |
| HNSC.turquoise | BLCA.turquoise | 9.44E-24 |
| HNSC.turquoise | UCEC.brown | 6.20E-48 |
| HNSC.yellow | GBM.turquoise | 7.52E-56 |
| HNSC.yellow | OV.turquoise | 1.84E-55 |
| HNSC.yellow | COADREAD.yellow | 9.35E-36 |
| HNSC.yellow | LUSC.turquoise | 2.69E-118 |
| HNSC.yellow | BRCA.yellow | 2.02E-88 |
| HNSC.yellow | KIRC.pink | 2.43E-60 |
| HNSC.yellow | BLCA.turquoise | 4.43E-80 |
| HNSC.yellow | UCEC.yellow | 5.41E-98 |
| KIRC.black | GBM.purple | 2.98E-08 |
| KIRC.black | OV.blue | 3.65E-18 |
| KIRC.black | COADREAD.brown | 8.80E-11 |
| KIRC.black | LUSC.blue | 2.13E-17 |
| KIRC.black | BRCA.blue | 4.00E-16 |
| KIRC.black | HNSC.blue | 6.43E-18 |
| KIRC.black | BLCA.yellow | 8.16E-15 |
| KIRC.black | UCEC.green | 1.68E-18 |
| KIRC.blue | GBM.green | 1.08E-87 |
| KIRC.blue | OV.brown | 9.70E-13 |
| KIRC.blue | COADREAD.purple | 1.31E-48 |
| KIRC.blue | LUSC.black | 1.29E-68 |
| KIRC.blue | HNSC.pink | 2.36E-70 |
| KIRC.blue | UCEC.black | 1.41E-68 |
| KIRC.brown | GBM.purple | 1.16E-16 |
| KIRC.brown | OV.blue | 9.71E-14 |
| KIRC.brown | COADREAD.red | 0.002307719 |
| KIRC.brown | LUSC.blue | 2.76E-05 |
| KIRC.brown | BRCA.green | 2.07E-43 |
| KIRC.brown | HNSC.green | 1.82E-09 |
| KIRC.brown | BLCA.yellow | 2.05E-17 |
| KIRC.brown | UCEC.green | 1.06E-15 |
| KIRC.green | GBM.turquoise | 7.48E-18 |
| KIRC.green | OV.turquoise | 1.58E-24 |
| KIRC.green | COADREAD.green | 8.60E-32 |
| KIRC.green | LUSC.yellow | 1.23E-96 |
| KIRC.green | BRCA.brown | 2.13E-71 |
| KIRC.green | HNSC.turquoise | 4.31E-40 |
| KIRC.green | BLCA.turquoise | 2.29E-38 |
| KIRC.green | UCEC.brown | 1.98E-52 |
| KIRC.greenyellow | GBM.pink | 0.000250163 |
| KIRC.greenyellow | OV.blue | 1.51E-08 |
| KIRC.greenyellow | COADREAD.magenta | 3.77E-19 |
| KIRC.greenyellow | LUSC.blue | 2.03E-06 |
| KIRC.greenyellow | BRCA.blue | 1.78E-10 |
| KIRC.greenyellow | HNSC.blue | 1.42E-07 |
| KIRC.greenyellow | BLCA.red | 2.90E-14 |
| KIRC.greenyellow | UCEC.red | 3.07E-22 |
| KIRC.magenta | GBM.turquoise | 7.35E-23 |
| KIRC.magenta | OV.turquoise | 2.81E-18 |
| KIRC.magenta | COADREAD.green | 2.34E-13 |
| KIRC.magenta | LUSC.turquoise | 1.29E-14 |
| KIRC.magenta | BRCA.yellow | 2.08E-13 |
| KIRC.magenta | HNSC.yellow | 2.05E-22 |
| KIRC.magenta | BLCA.turquoise | 4.39E-16 |
| KIRC.magenta | UCEC.yellow | 3.49E-31 |
| KIRC.pink | GBM.turquoise | 3.57E-24 |
| KIRC.pink | OV.turquoise | 4.24E-55 |
| KIRC.pink | COADREAD.yellow | 1.54E-32 |
| KIRC.pink | LUSC.turquoise | 6.76E-70 |
| KIRC.pink | BRCA.yellow | 8.40E-84 |
| KIRC.pink | HNSC.yellow | 3.28E-60 |
| KIRC.pink | BLCA.turquoise | 1.74E-44 |
| KIRC.pink | UCEC.yellow | 4.22E-57 |
| KIRC.purple | COADREAD.blue | 0.005537789 |
| KIRC.purple | BRCA.red | 2.87E-37 |
| KIRC.purple | UCEC.pink | 2.18E-37 |
| KIRC.red | GBM.pink | 4.34E-24 |
| KIRC.red | OV.blue | 6.66E-43 |
| KIRC.red | COADREAD.pink | 1.04E-13 |
| KIRC.red | LUSC.blue | 1.26E-52 |
| KIRC.red | BRCA.blue | 1.66E-54 |
| KIRC.red | HNSC.blue | 3.72E-45 |
| KIRC.red | BLCA.brown | 2.47E-33 |
| KIRC.red | UCEC.green | 4.60E-29 |
| KIRC.turquoise | GBM.yellow | 5.32E-05 |
| KIRC.turquoise | LUSC.red | 0.001475322 |
| KIRC.turquoise | BLCA.blue | 0.007362857 |
| KIRC.yellow | GBM.black | 6.05E-54 |
| KIRC.yellow | OV.green | 8.70E-20 |
| KIRC.yellow | BRCA.turquoise | 1.17E-22 |
| KIRC.yellow | BLCA.blue | 0.01085711 |
| KIRC.yellow | UCEC.blue | 6.11E-20 |
| BLCA.black | HNSC.green | 0.000186086 |
| BLCA.blue | LUSC.brown | 6.56E-16 |
| BLCA.blue | BRCA.turquoise | 8.33E-09 |
| BLCA.blue | HNSC.brown | 3.84E-08 |
| BLCA.blue | KIRC.turquoise | 0.000131954 |
| BLCA.brown | GBM.pink | 6.02E-16 |
| BLCA.brown | OV.blue | 3.58E-78 |
| BLCA.brown | COADREAD.brown | 7.84E-27 |
| BLCA.brown | LUSC.blue | 3.18E-97 |
| BLCA.brown | BRCA.blue | 1.09E-95 |
| BLCA.brown | HNSC.blue | 2.46E-96 |
| BLCA.brown | KIRC.red | 1.51E-37 |
| BLCA.brown | UCEC.green | 8.65E-48 |
| BLCA.green | LUSC.brown | 4.55E-29 |
| BLCA.green | HNSC.red | 4.56E-42 |
| BLCA.green | UCEC.turquoise | 0.007472986 |
| BLCA.pink | GBM.tan | 1.43E-41 |
| BLCA.pink | OV.yellow | 2.27E-40 |
| BLCA.pink | COADREAD.yellow | 1.54E-10 |
| BLCA.pink | LUSC.yellow | 2.85E-21 |
| BLCA.pink | BRCA.black | 5.02E-43 |
| BLCA.pink | HNSC.magenta | 2.29E-51 |
| BLCA.pink | KIRC.green | 1.92E-07 |
| BLCA.pink | UCEC.magenta | 6.37E-55 |
| BLCA.red | OV.blue | 3.72E-08 |
| BLCA.red | COADREAD.magenta | 8.58E-40 |
| BLCA.red | LUSC.blue | 3.27E-14 |
| BLCA.red | BRCA.blue | 2.91E-09 |
| BLCA.red | HNSC.black | 2.36E-09 |
| BLCA.red | KIRC.greenyellow | 1.22E-14 |
| BLCA.red | UCEC.red | 3.81E-30 |
| BLCA.turquoise | GBM.turquoise | 3.23E-52 |
| BLCA.turquoise | OV.turquoise | 7.02E-88 |
| BLCA.turquoise | COADREAD.yellow | 8.47E-45 |
| BLCA.turquoise | LUSC.turquoise | 7.53E-60 |
| BLCA.turquoise | BRCA.yellow | 4.51E-73 |
| BLCA.turquoise | HNSC.yellow | 4.86E-74 |
| BLCA.turquoise | KIRC.pink | 2.37E-43 |
| BLCA.turquoise | UCEC.yellow | 7.90E-77 |
| BLCA.yellow | GBM.brown | 0.000627877 |
| BLCA.yellow | OV.blue | 0.000259226 |
| BLCA.yellow | COADREAD.brown | 0.003223737 |
| BLCA.yellow | LUSC.blue | 9.16E-05 |
| BLCA.yellow | BRCA.green | 8.26E-22 |
| BLCA.yellow | HNSC.green | 3.52E-23 |
| BLCA.yellow | KIRC.black | 9.87E-16 |
| BLCA.yellow | UCEC.green | 5.59E-10 |
| UCEC.black | GBM.green | 1.04E-57 |
| UCEC.black | OV.brown | 2.24E-08 |
| UCEC.black | COADREAD.purple | 6.97E-38 |
| UCEC.black | LUSC.black | 1.79E-53 |
| UCEC.black | HNSC.pink | 5.16E-41 |
| UCEC.black | KIRC.blue | 1.97E-65 |
| UCEC.black | BLCA.blue | 0.01510793 |
| UCEC.blue | GBM.black | 4.20E-27 |
| UCEC.blue | OV.green | 3.91E-24 |
| UCEC.blue | BRCA.turquoise | 8.61E-24 |
| UCEC.blue | KIRC.yellow | 1.37E-24 |
| UCEC.blue | BLCA.blue | 0.000279847 |
| UCEC.brown | GBM.turquoise | 1.72E-16 |
| UCEC.brown | OV.turquoise | 5.03E-53 |
| UCEC.brown | COADREAD.green | 1.49E-28 |
| UCEC.brown | LUSC.yellow | 2.79E-47 |
| UCEC.brown | BRCA.brown | 2.73E-61 |
| UCEC.brown | HNSC.turquoise | 1.87E-44 |
| UCEC.brown | KIRC.green | 1.52E-54 |
| UCEC.brown | BLCA.turquoise | 2.25E-43 |
| UCEC.green | GBM.pink | 1.08E-19 |
| UCEC.green | OV.blue | 6.98E-95 |
| UCEC.green | COADREAD.brown | 2.08E-12 |
| UCEC.green | LUSC.blue | 5.76E-69 |
| UCEC.green | BRCA.blue | 1.35E-59 |
| UCEC.green | HNSC.blue | 3.22E-55 |
| UCEC.green | KIRC.red | 5.11E-24 |
| UCEC.green | BLCA.brown | 3.21E-45 |
| UCEC.magenta | GBM.tan | 1.52E-69 |
| UCEC.magenta | OV.yellow | 4.75E-43 |
| UCEC.magenta | COADREAD.yellow | 7.11E-11 |
| UCEC.magenta | LUSC.yellow | 1.81E-29 |
| UCEC.magenta | BRCA.black | 6.93E-57 |
| UCEC.magenta | HNSC.magenta | 1.85E-58 |
| UCEC.magenta | KIRC.green | 1.85E-13 |
| UCEC.magenta | BLCA.pink | 3.92E-57 |
| UCEC.pink | BRCA.red | 9.08E-34 |
| UCEC.pink | KIRC.purple | 1.31E-38 |
| UCEC.pink | BLCA.blue | 0.007672225 |
| UCEC.red | GBM.pink | 0.000169972 |
| UCEC.red | OV.blue | 1.74E-24 |
| UCEC.red | COADREAD.magenta | 7.97E-28 |
| UCEC.red | LUSC.blue | 1.26E-32 |
| UCEC.red | BRCA.blue | 1.08E-29 |
| UCEC.red | HNSC.blue | 6.67E-22 |
| UCEC.red | KIRC.greenyellow | 5.87E-21 |
| UCEC.red | BLCA.red | 1.13E-25 |
| UCEC.turquoise | GBM.yellow | 6.82E-22 |
| UCEC.turquoise | OV.brown | 1.71E-40 |
| UCEC.turquoise | LUSC.pink | 3.89E-30 |
| UCEC.turquoise | BRCA.turquoise | 4.03E-12 |
| UCEC.turquoise | HNSC.brown | 1.79E-05 |
| UCEC.turquoise | KIRC.turquoise | 1.34E-07 |
| UCEC.yellow | GBM.turquoise | 5.63E-49 |
| UCEC.yellow | OV.turquoise | 1.21E-84 |
| UCEC.yellow | COADREAD.yellow | 8.05E-39 |
| UCEC.yellow | LUSC.turquoise | 1.85E-59 |
| UCEC.yellow | BRCA.yellow | 4.22E-88 |
| UCEC.yellow | HNSC.yellow | 1.93E-88 |
| UCEC.yellow | KIRC.pink | 4.17E-45 |
| UCEC.yellow | BLCA.turquoise | 3.73E-74 |

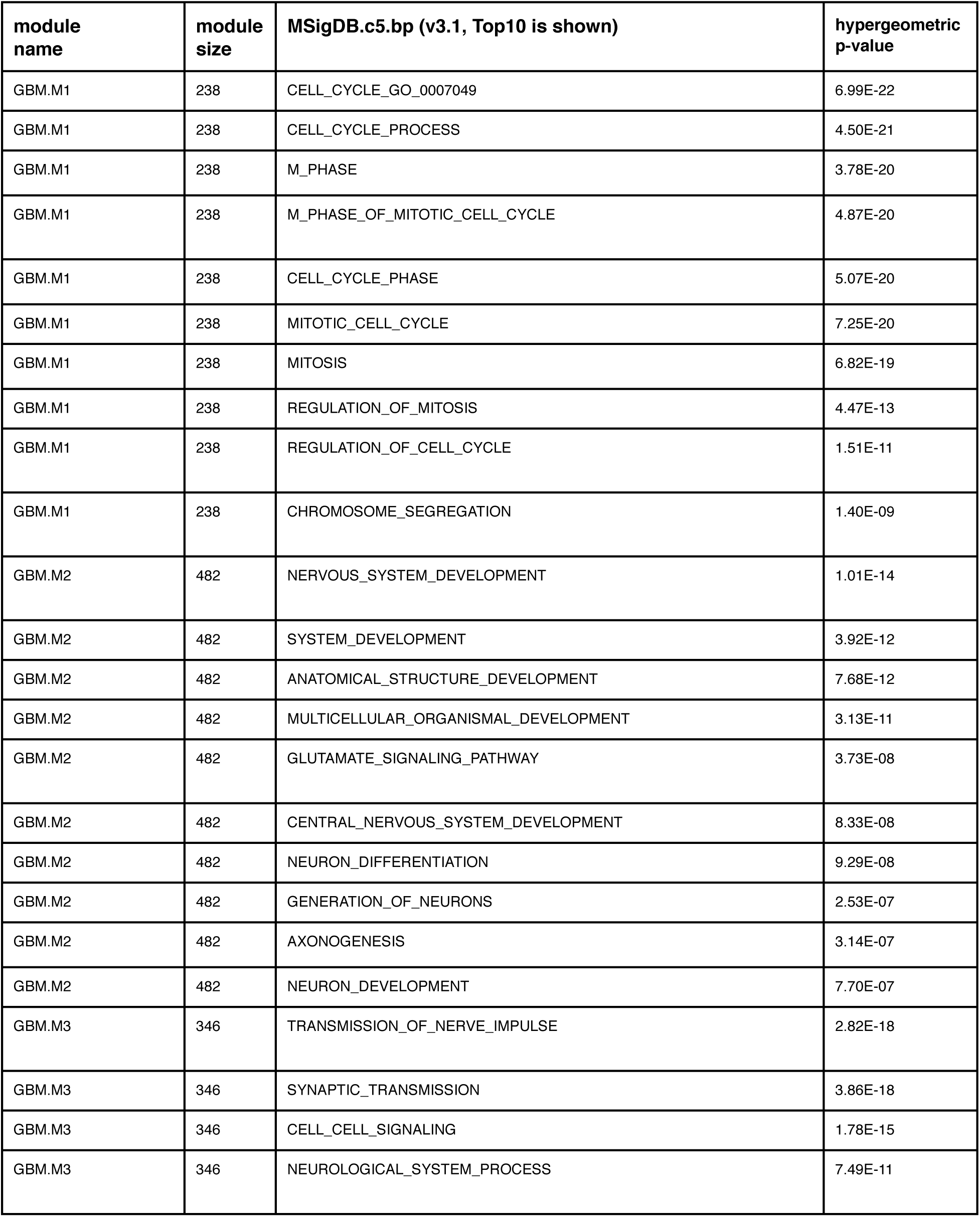
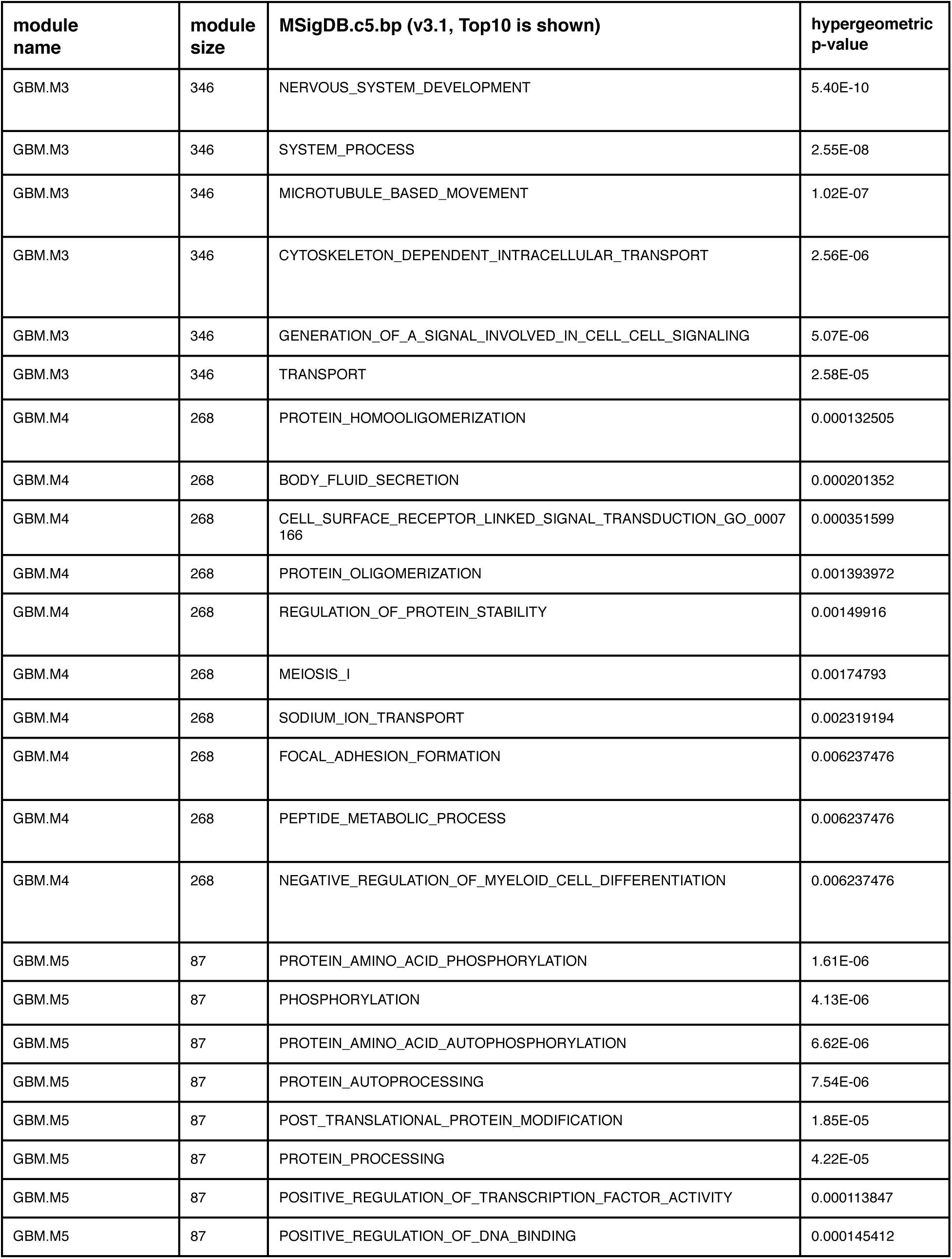
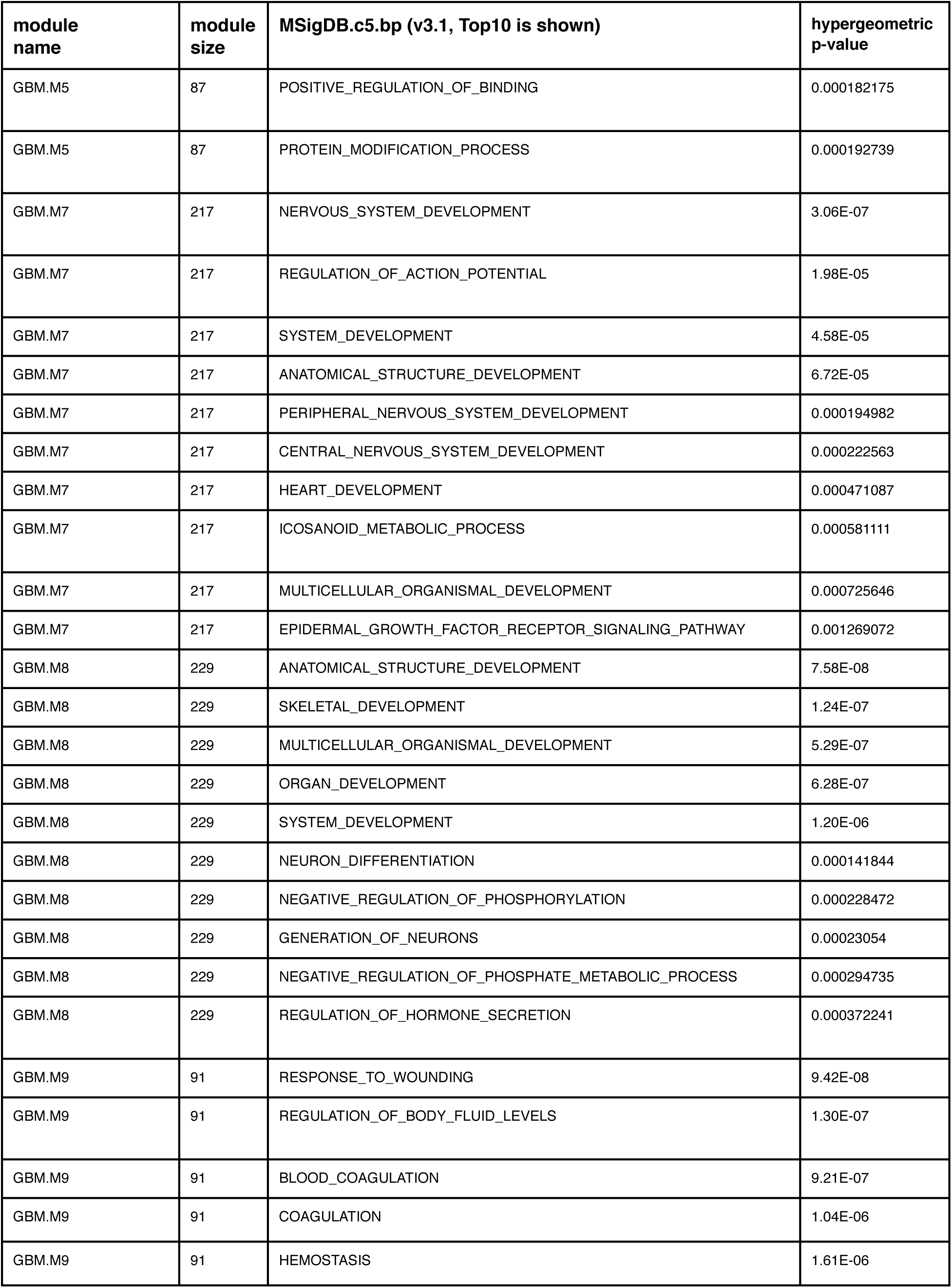
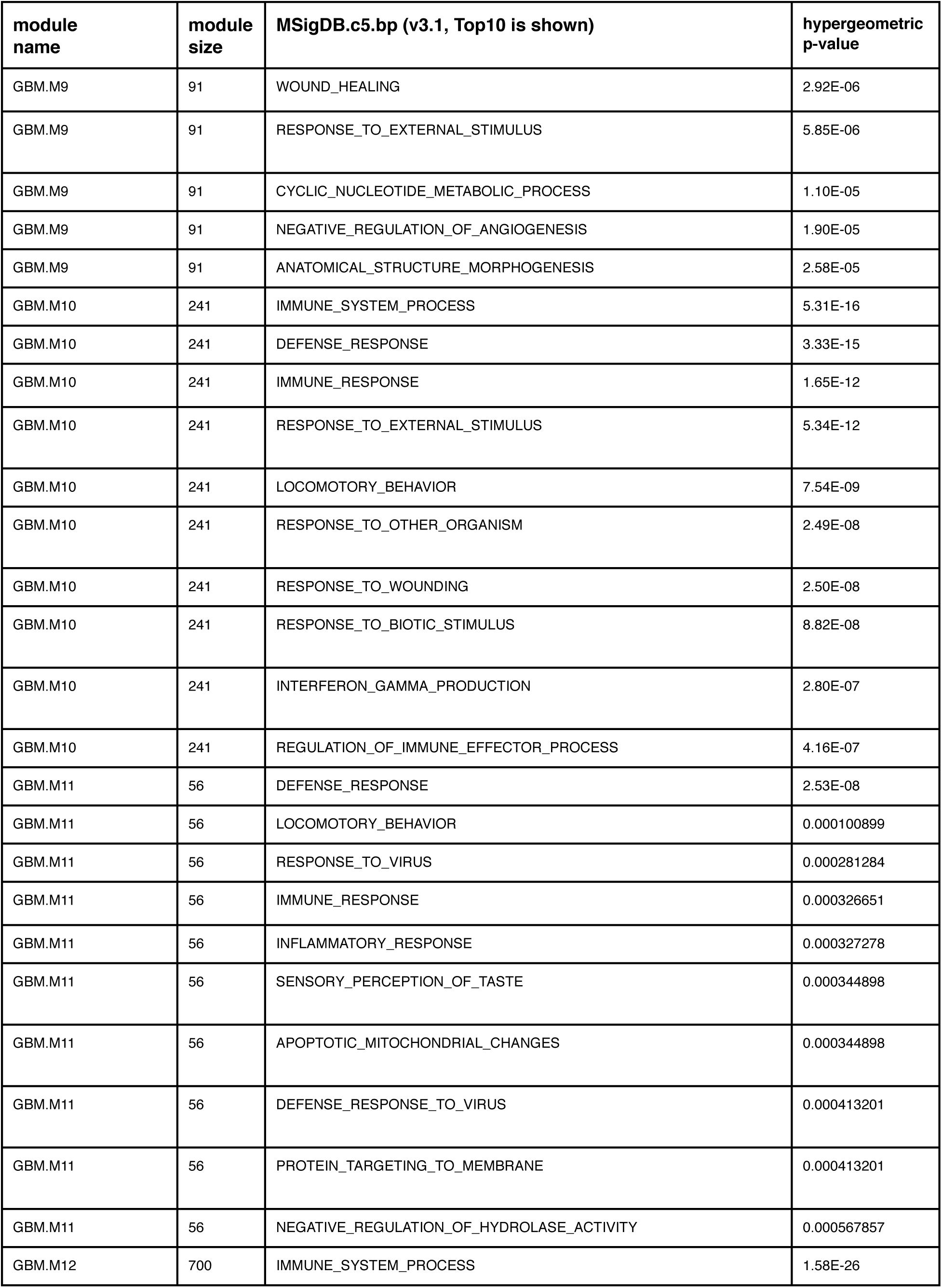
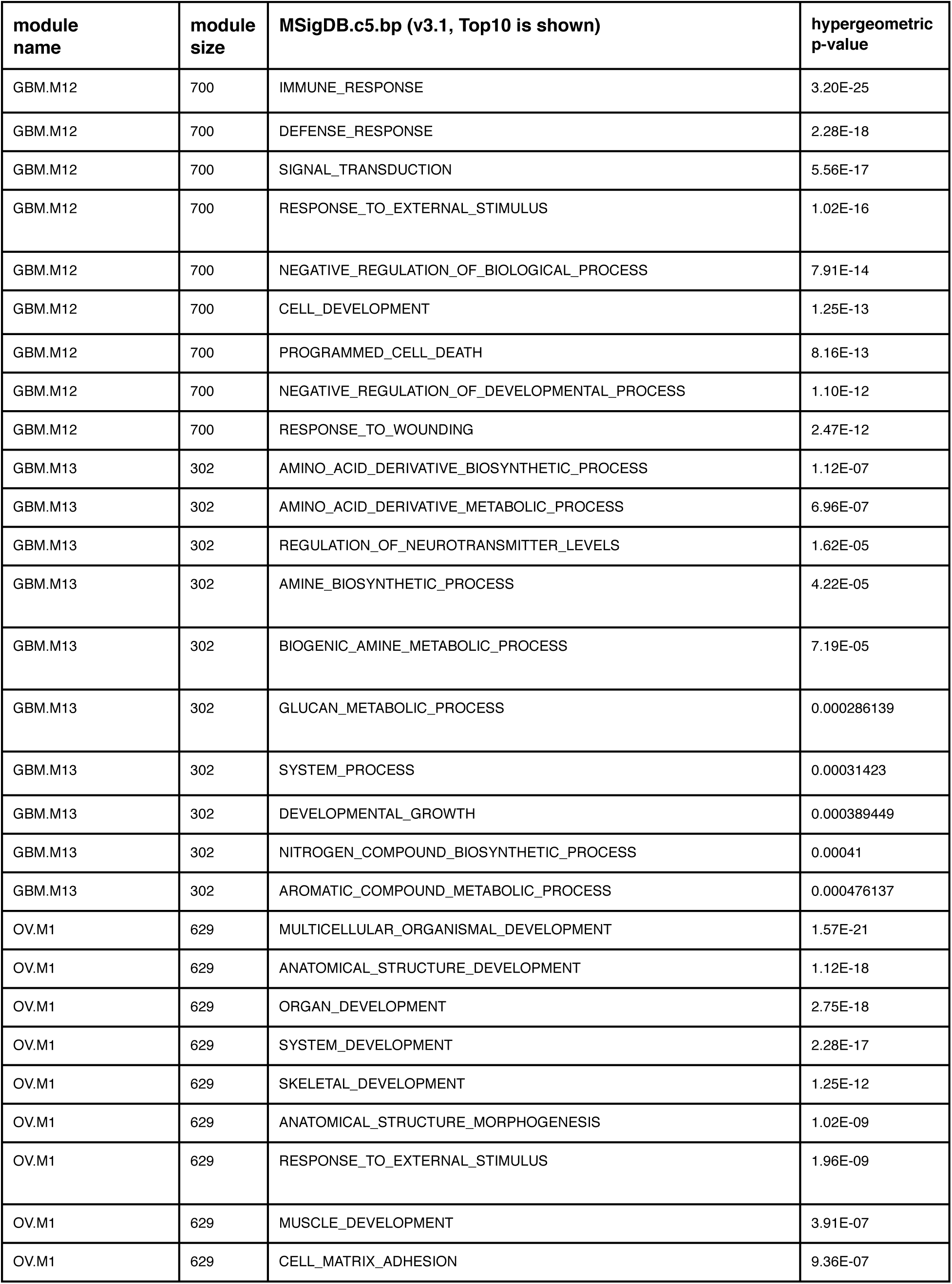
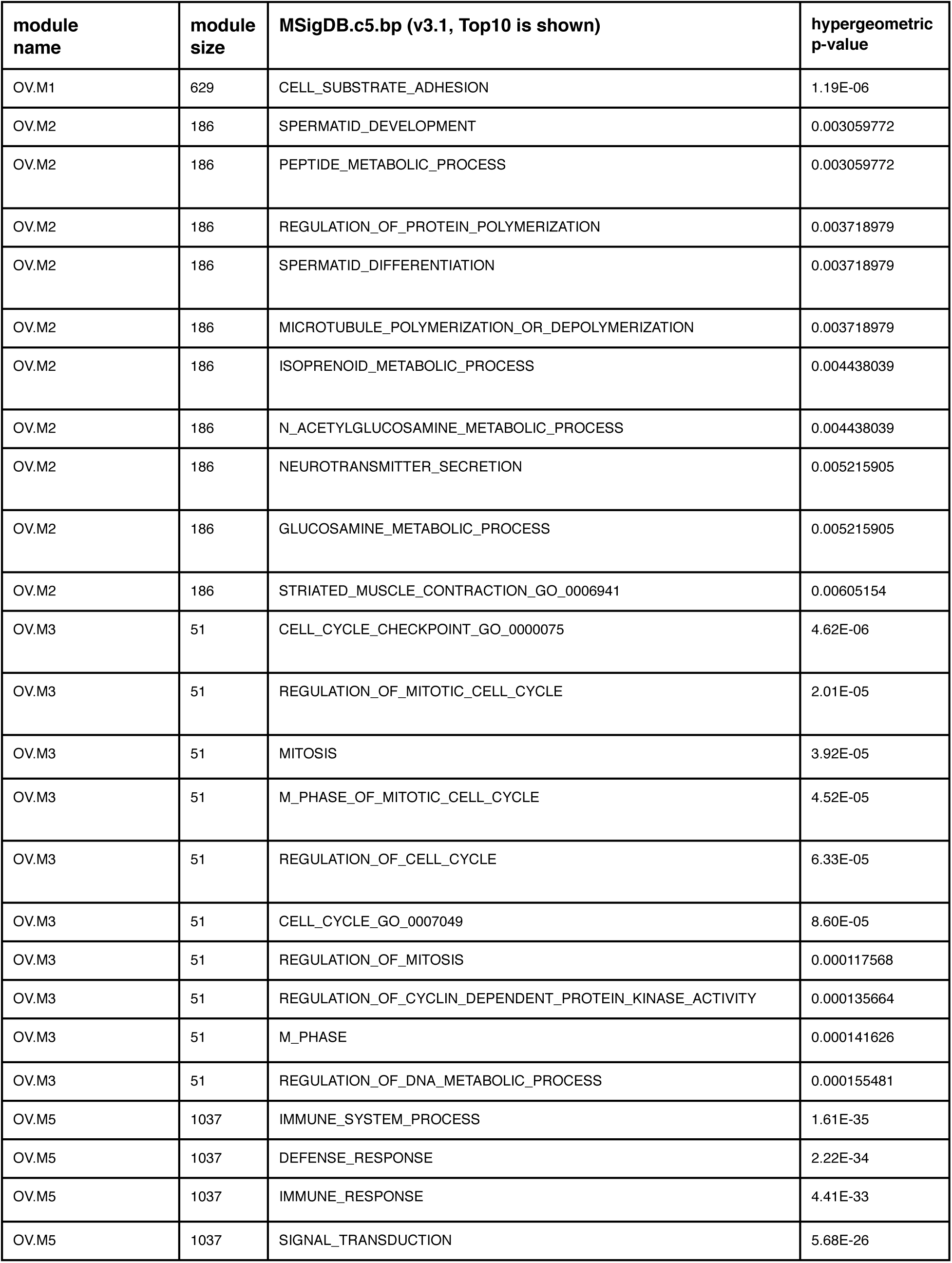
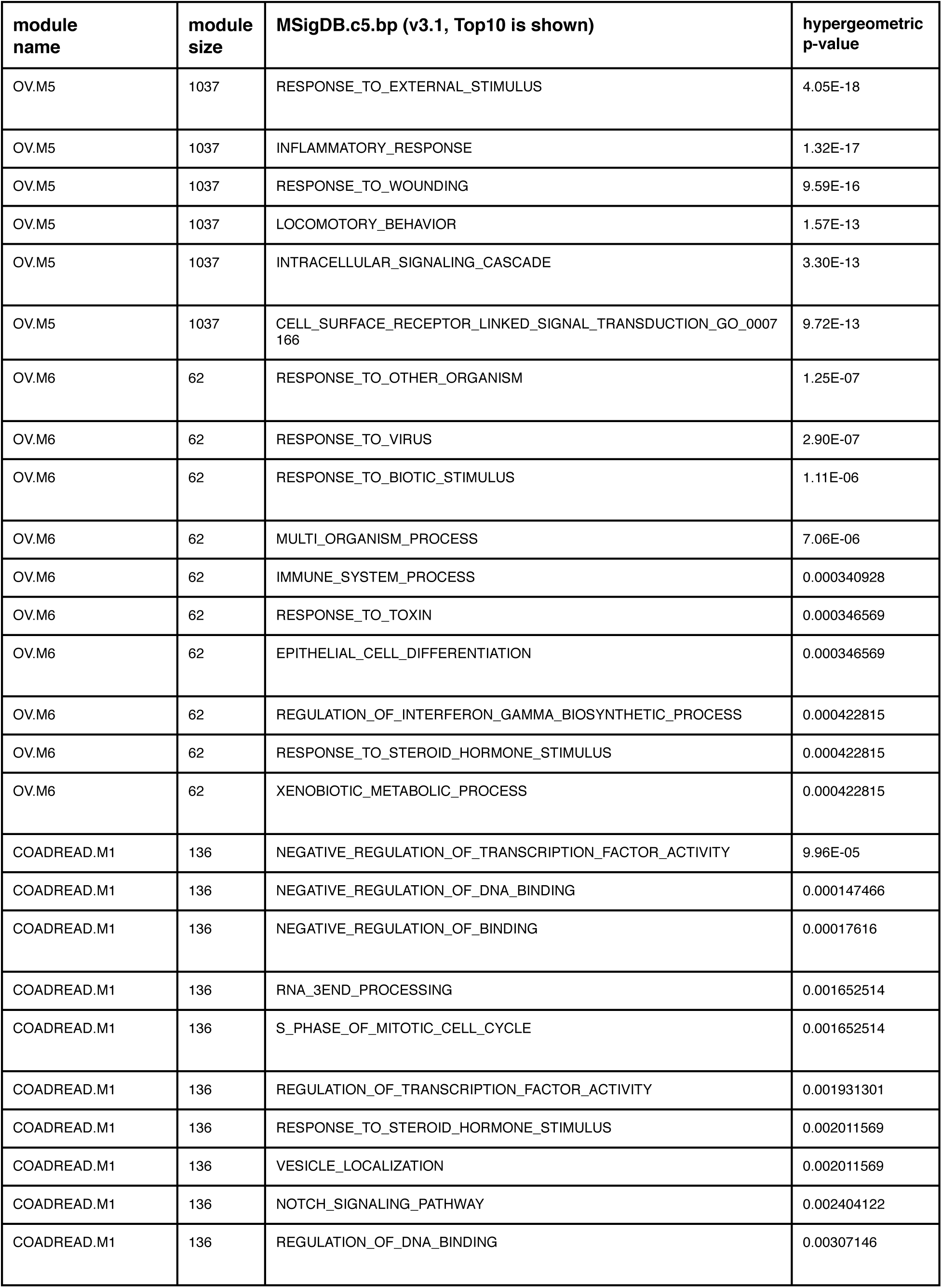
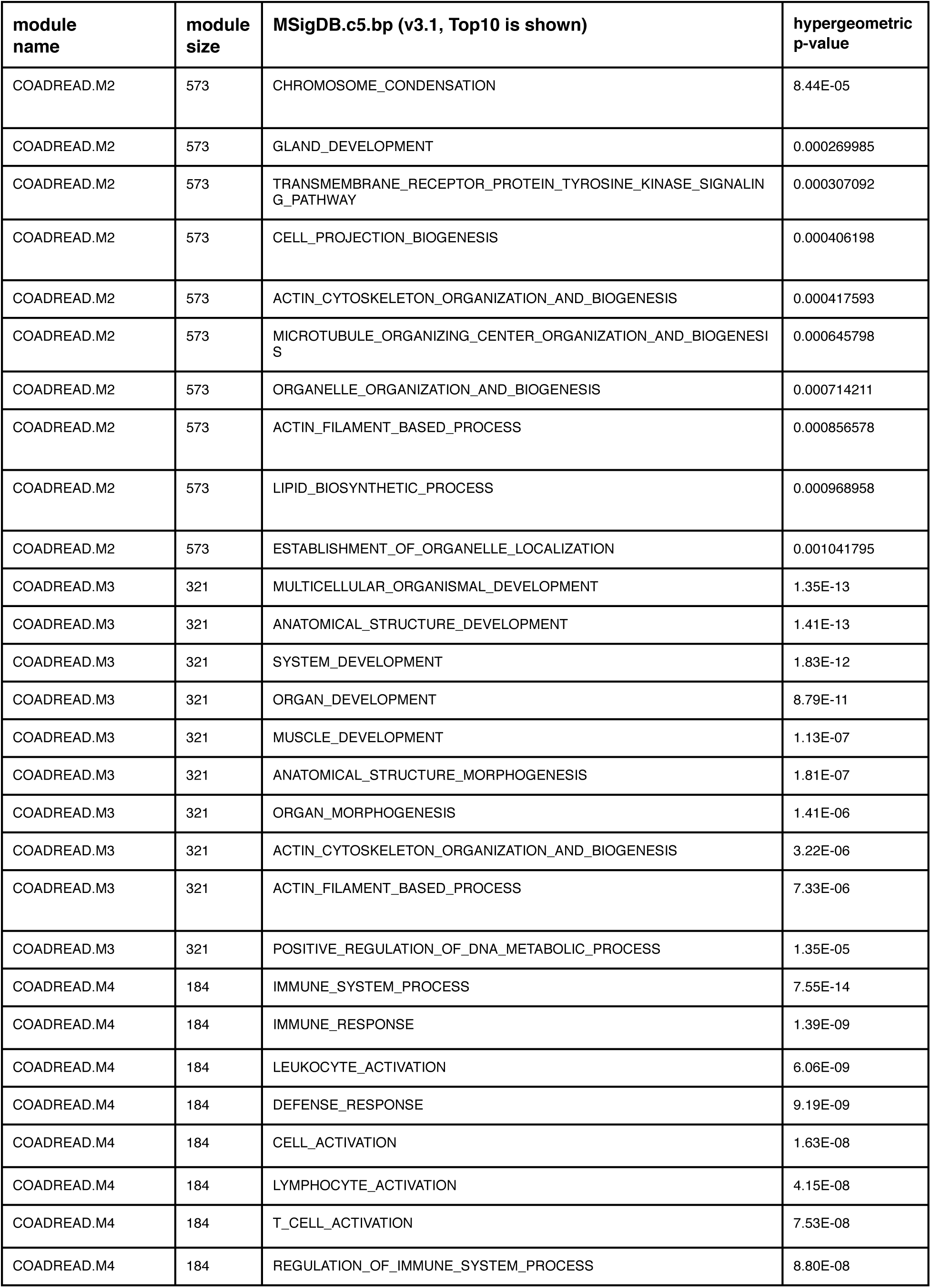
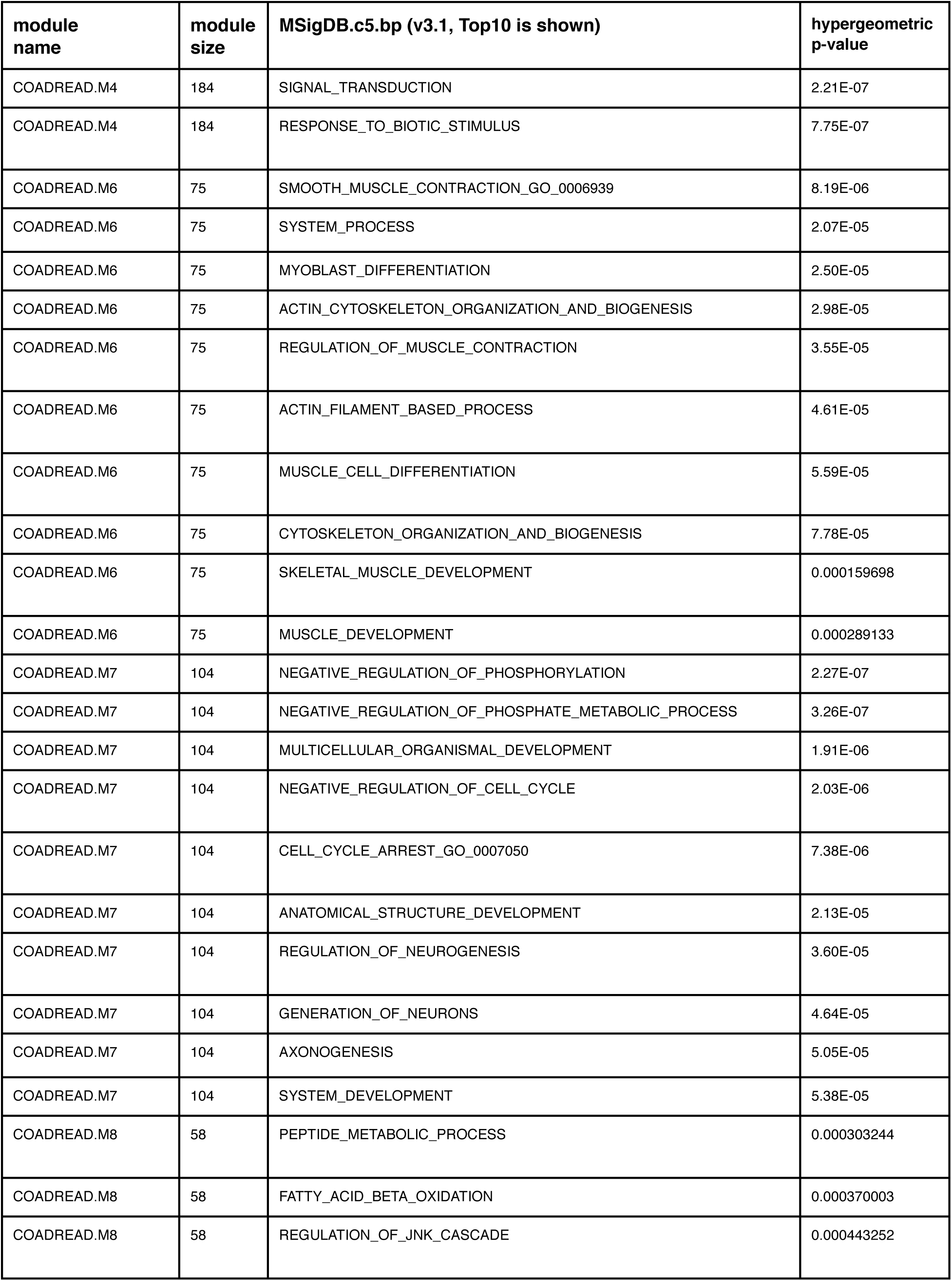
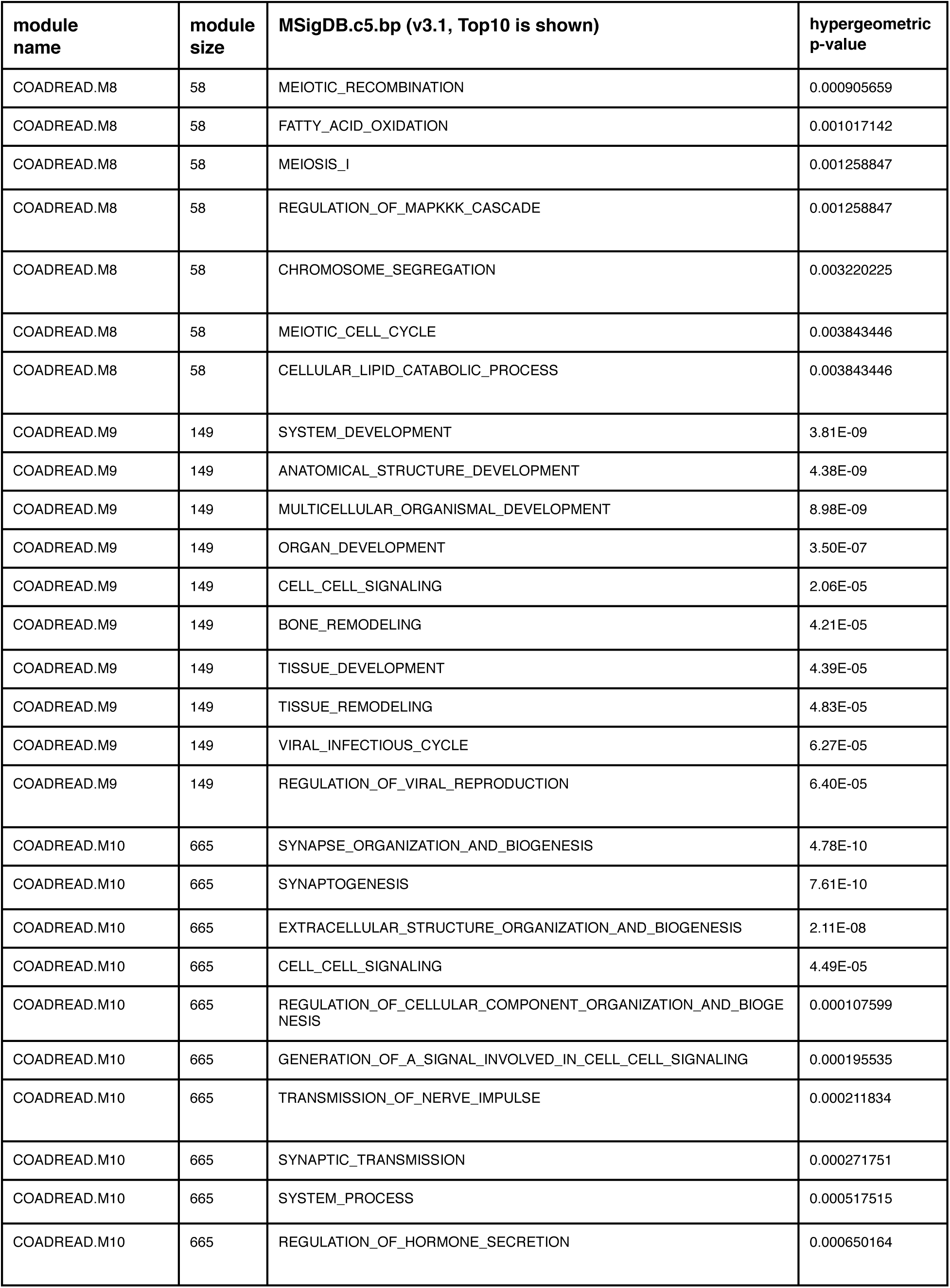
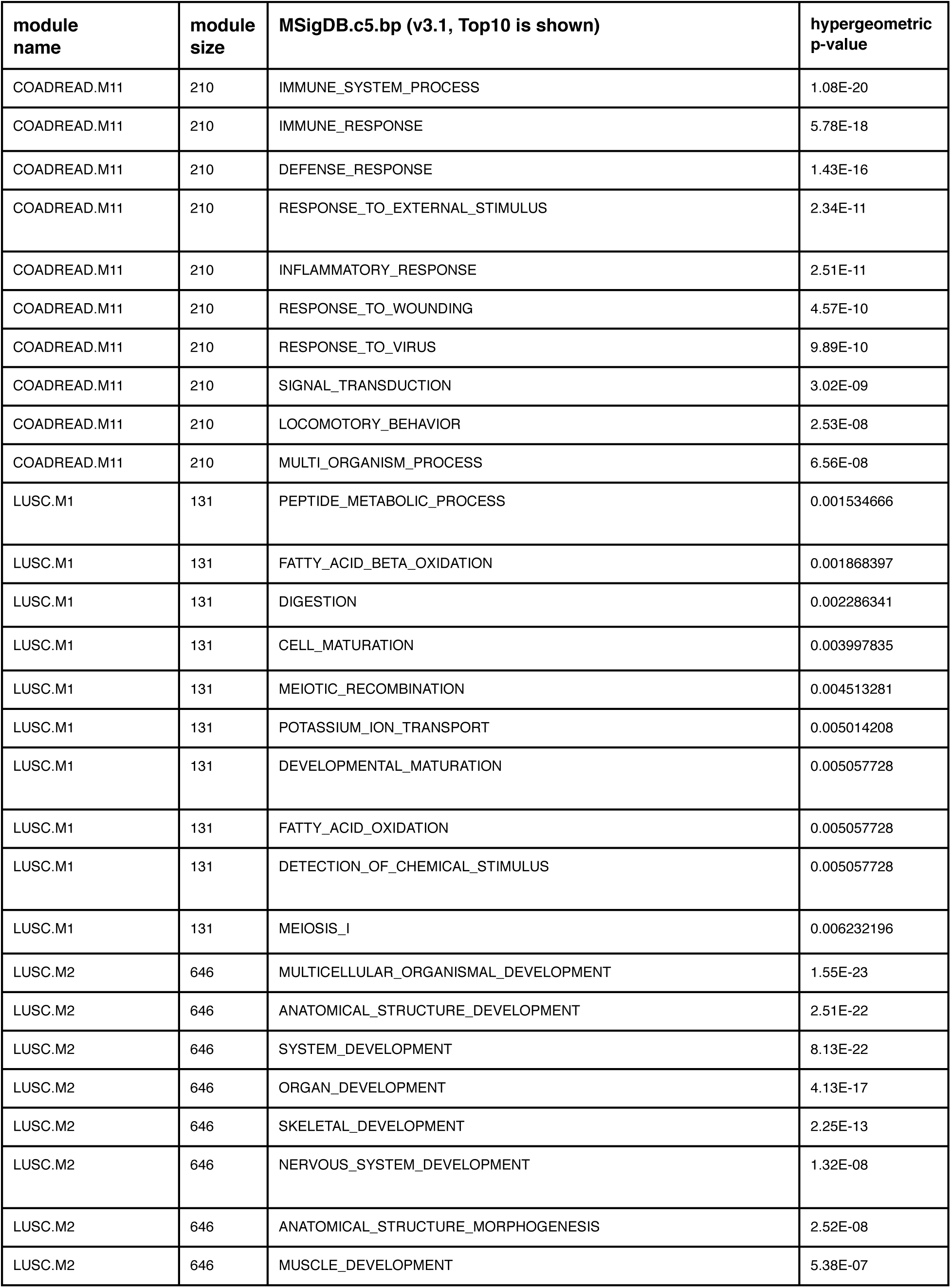
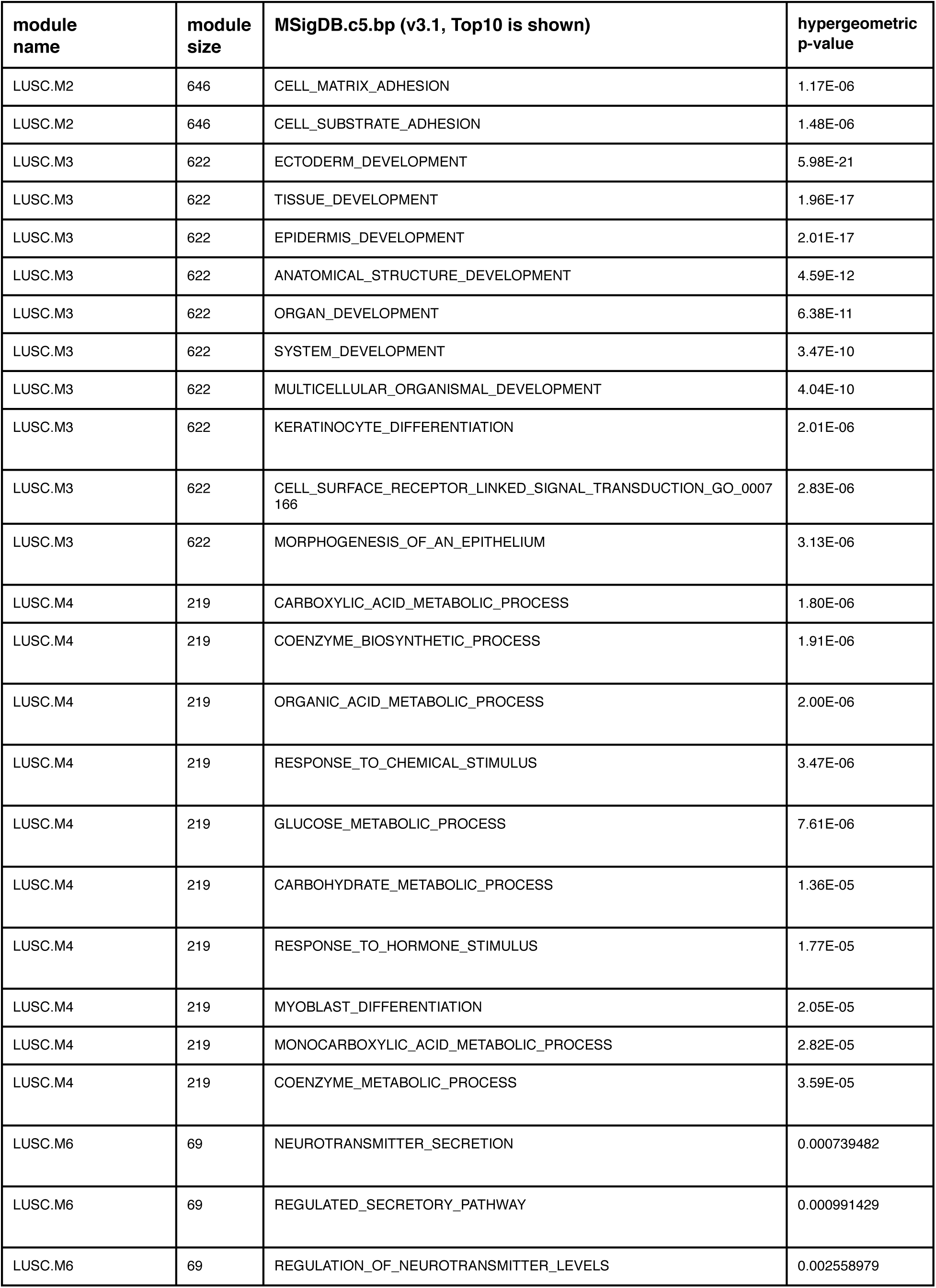
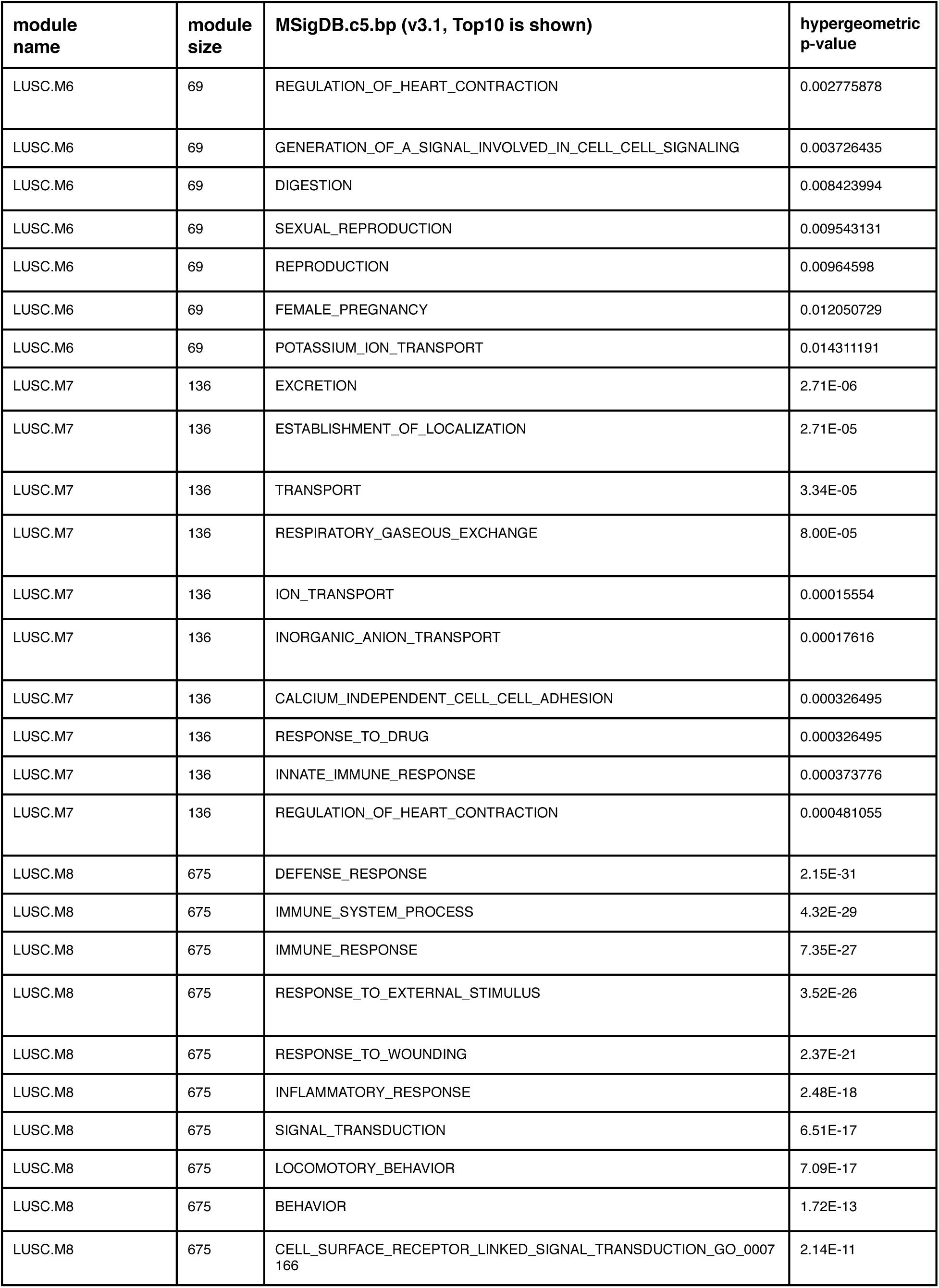
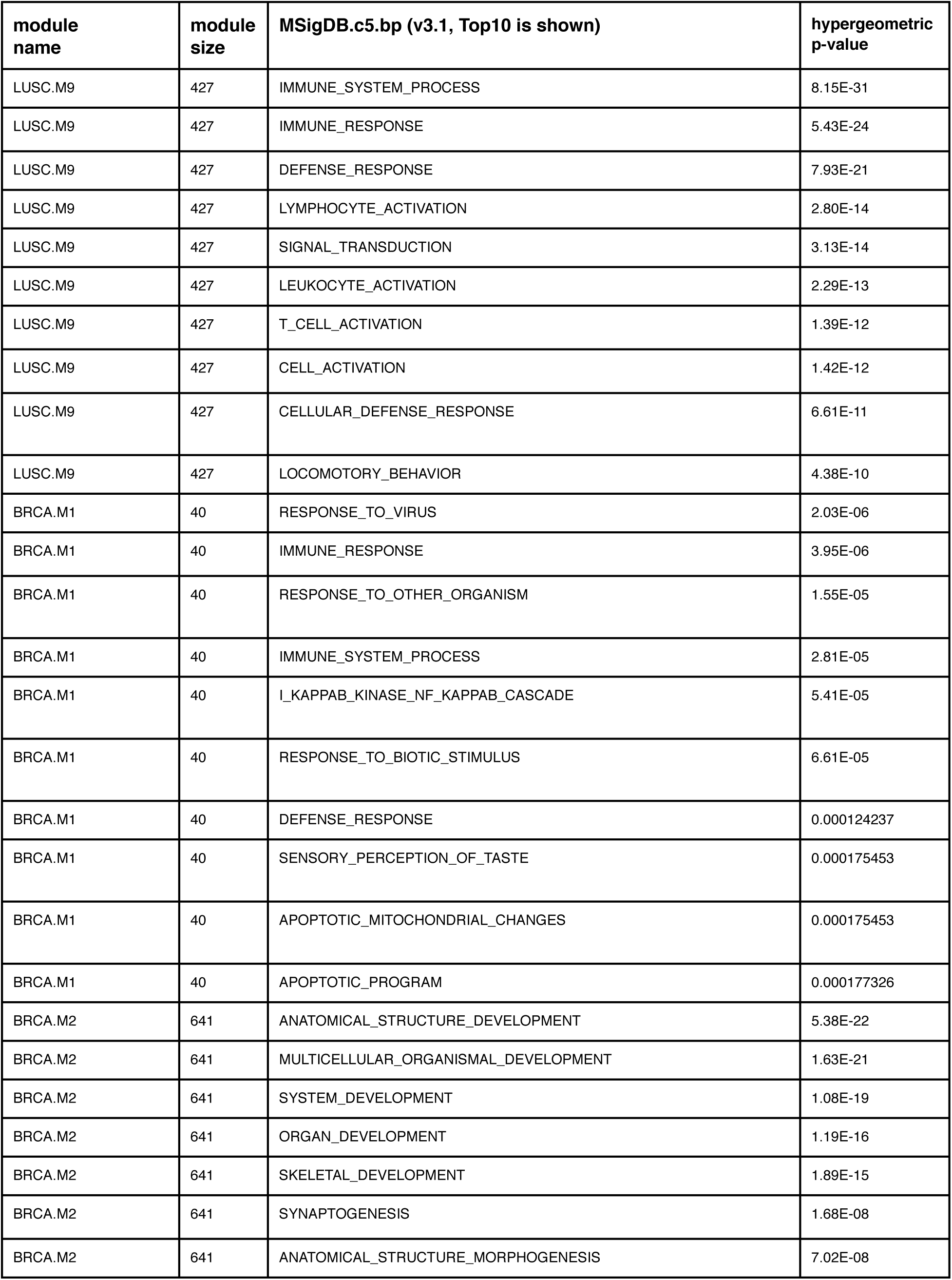
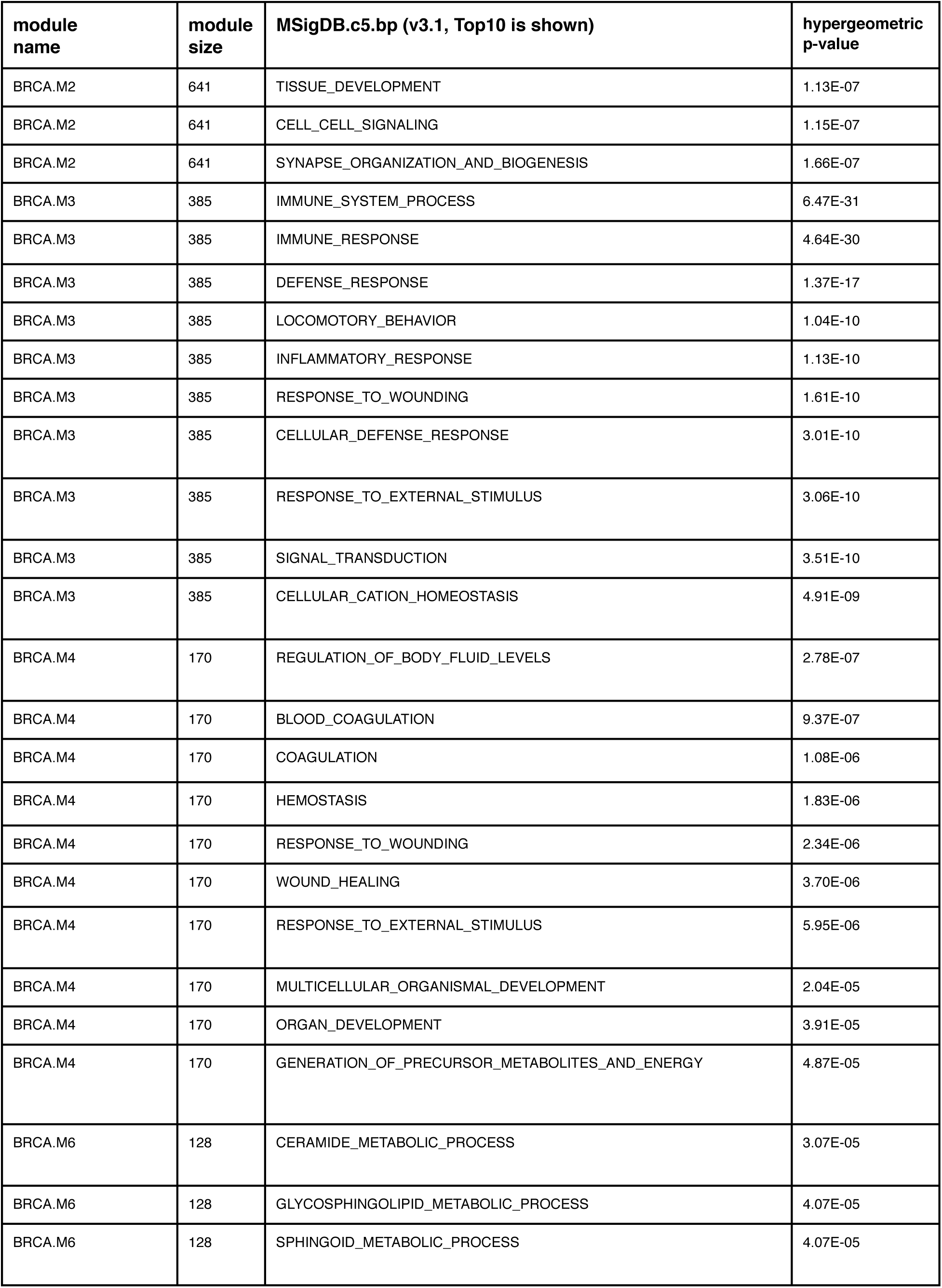
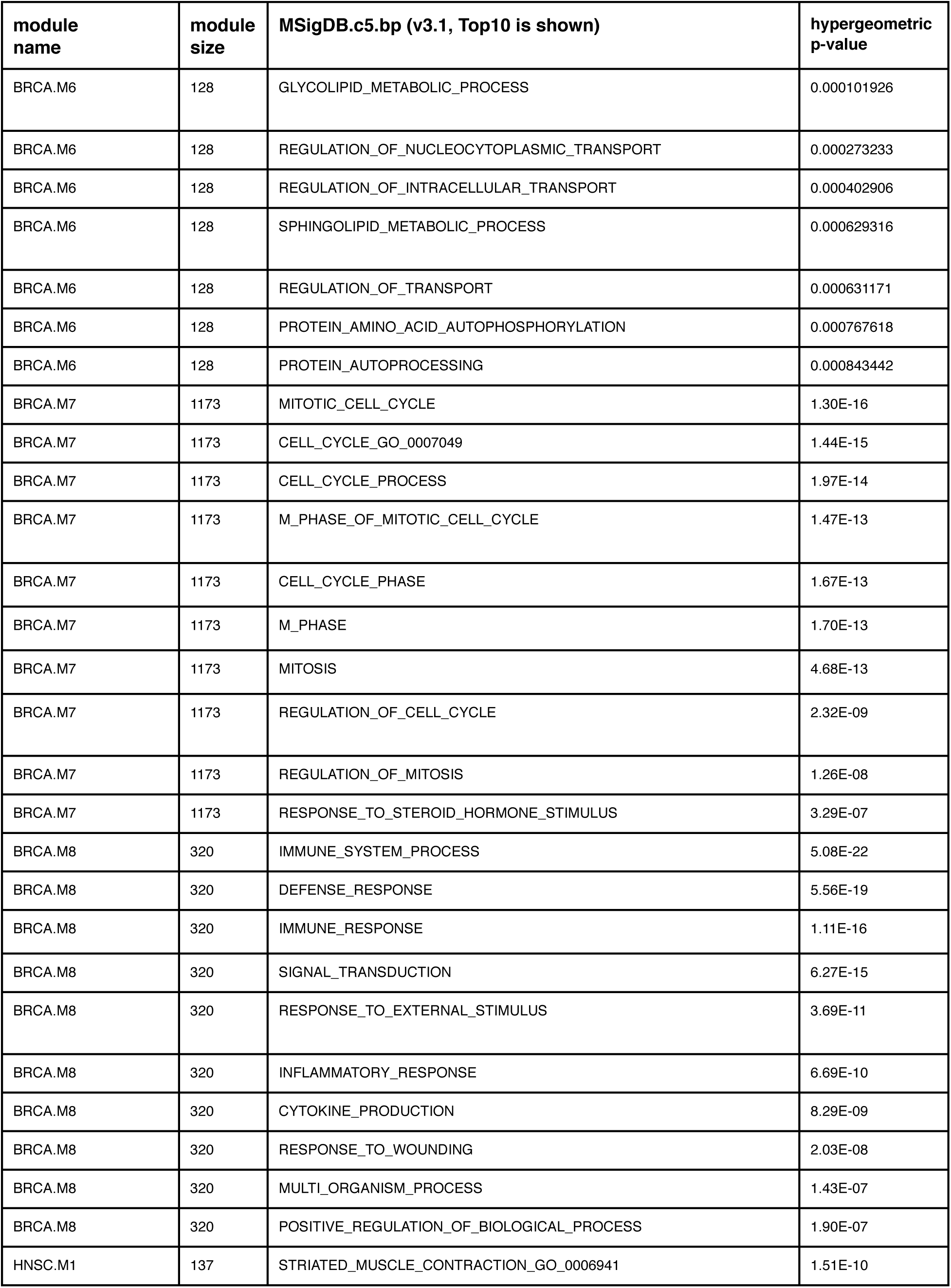
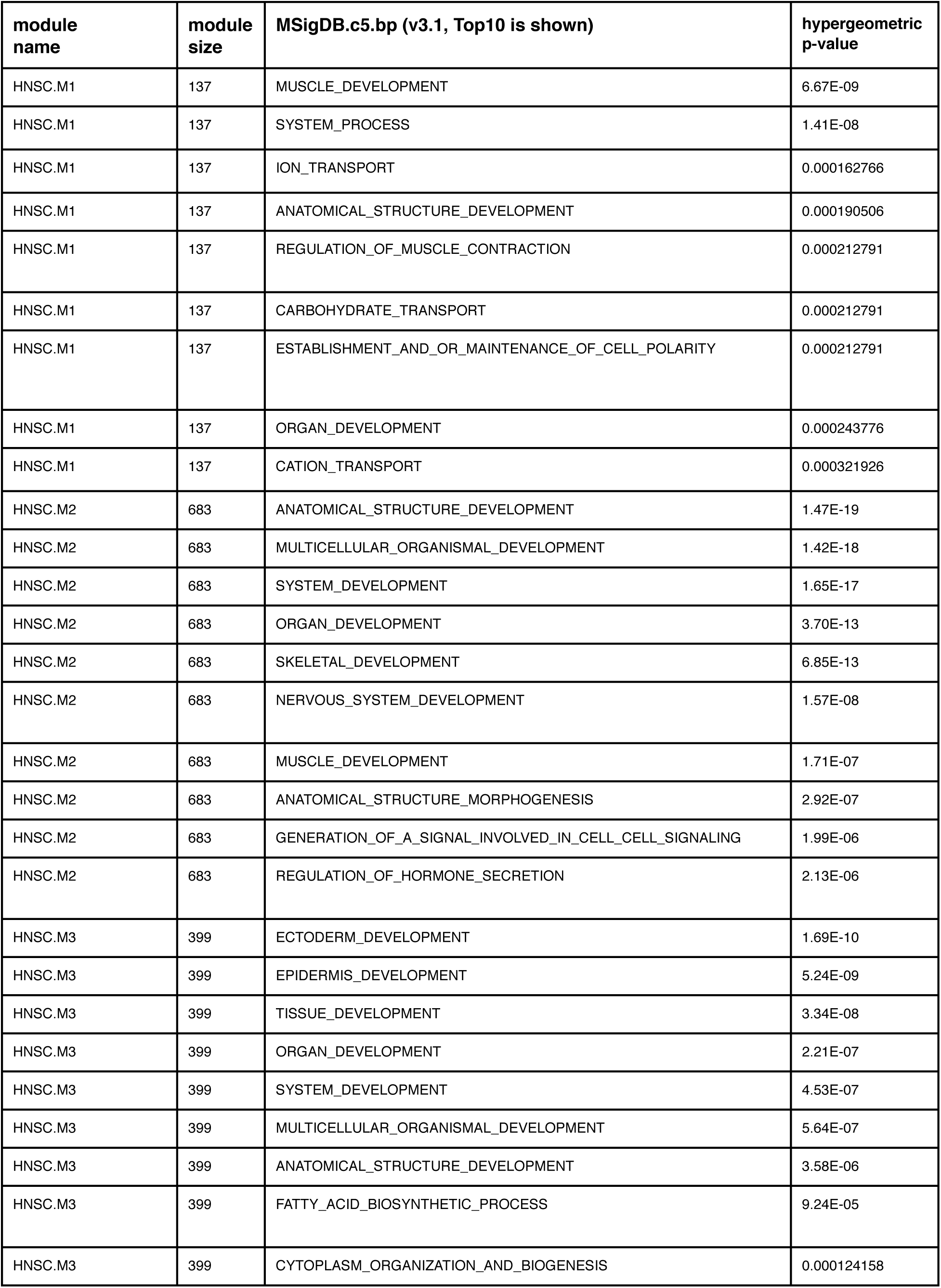
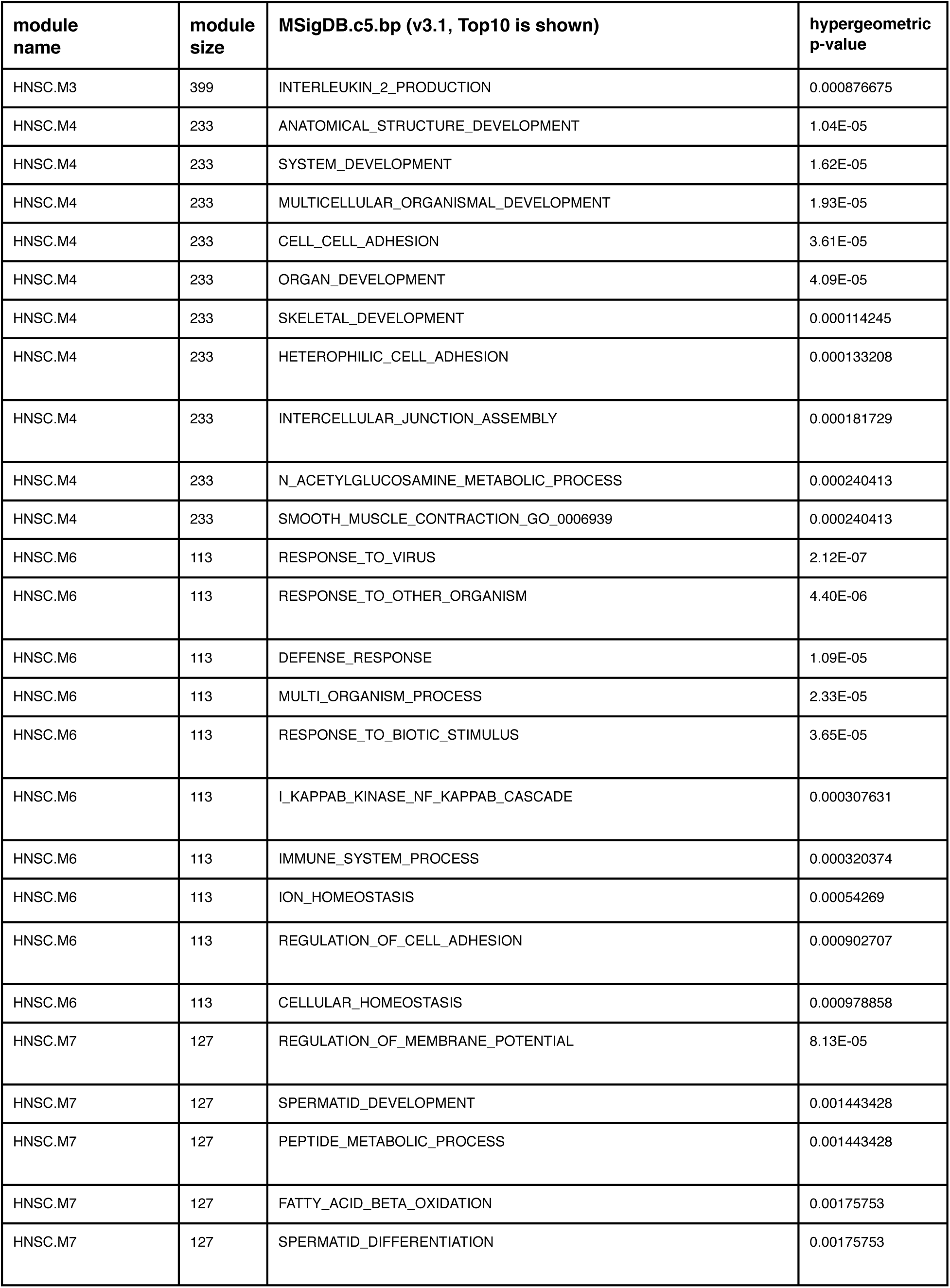
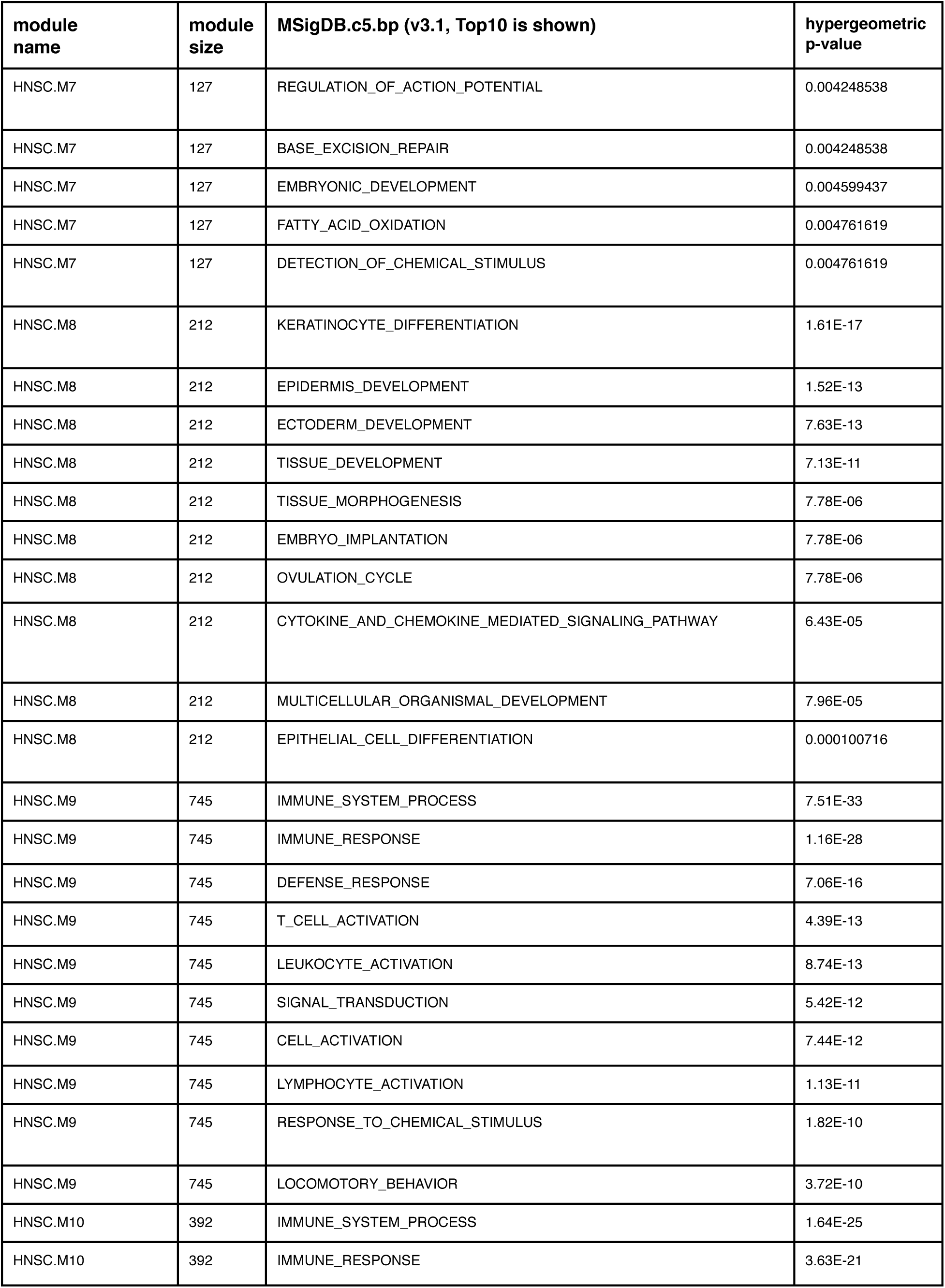
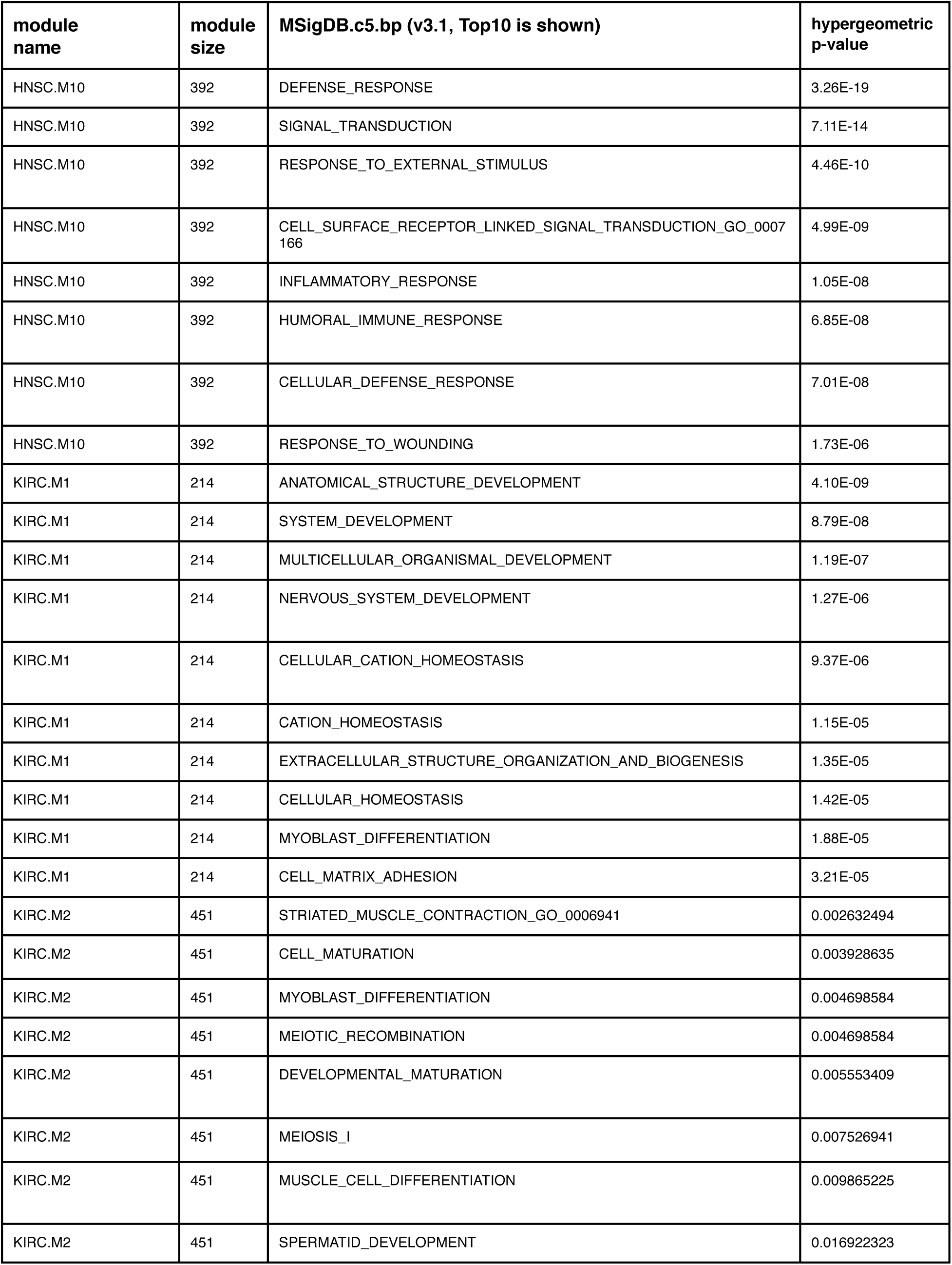
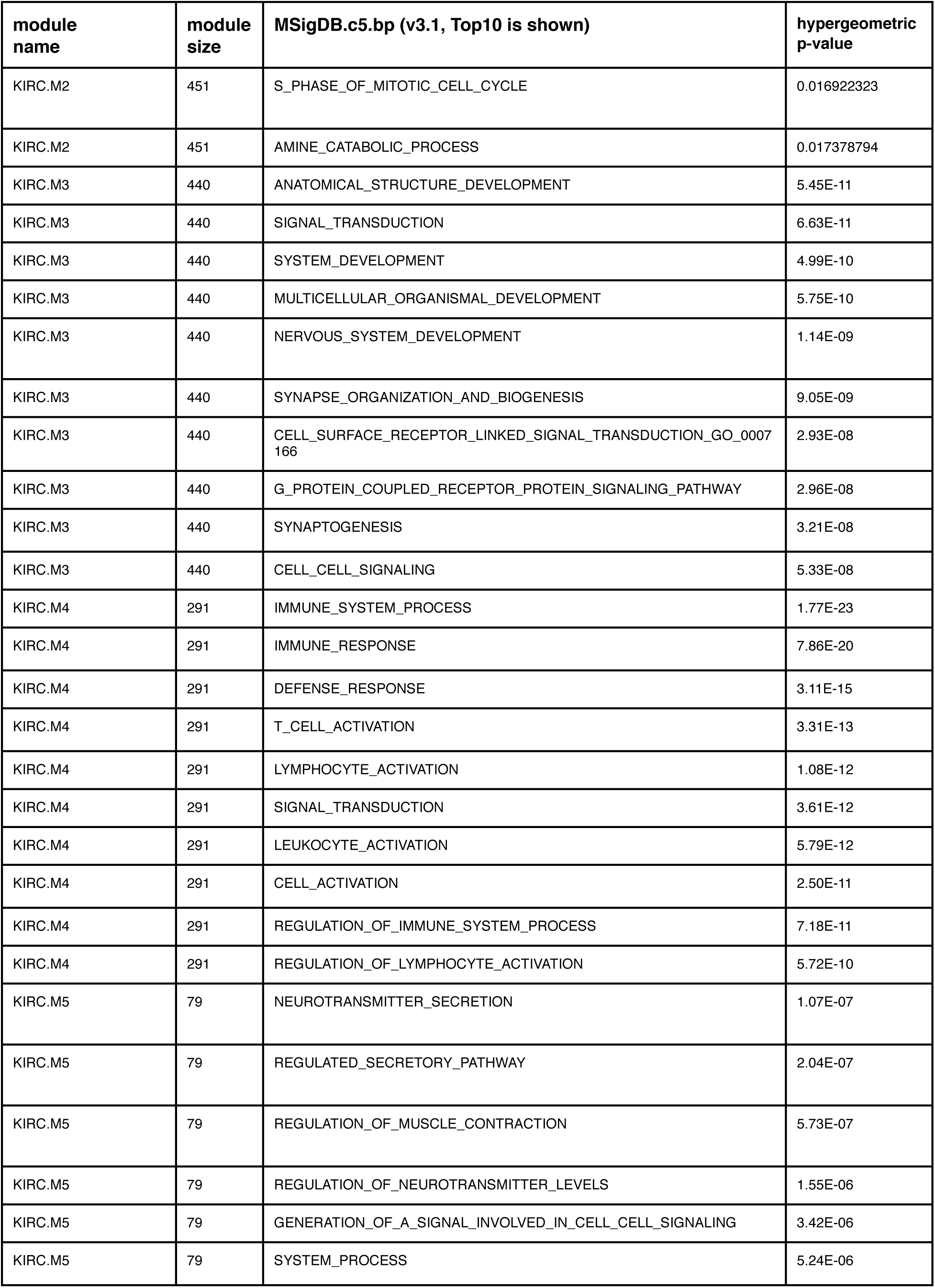
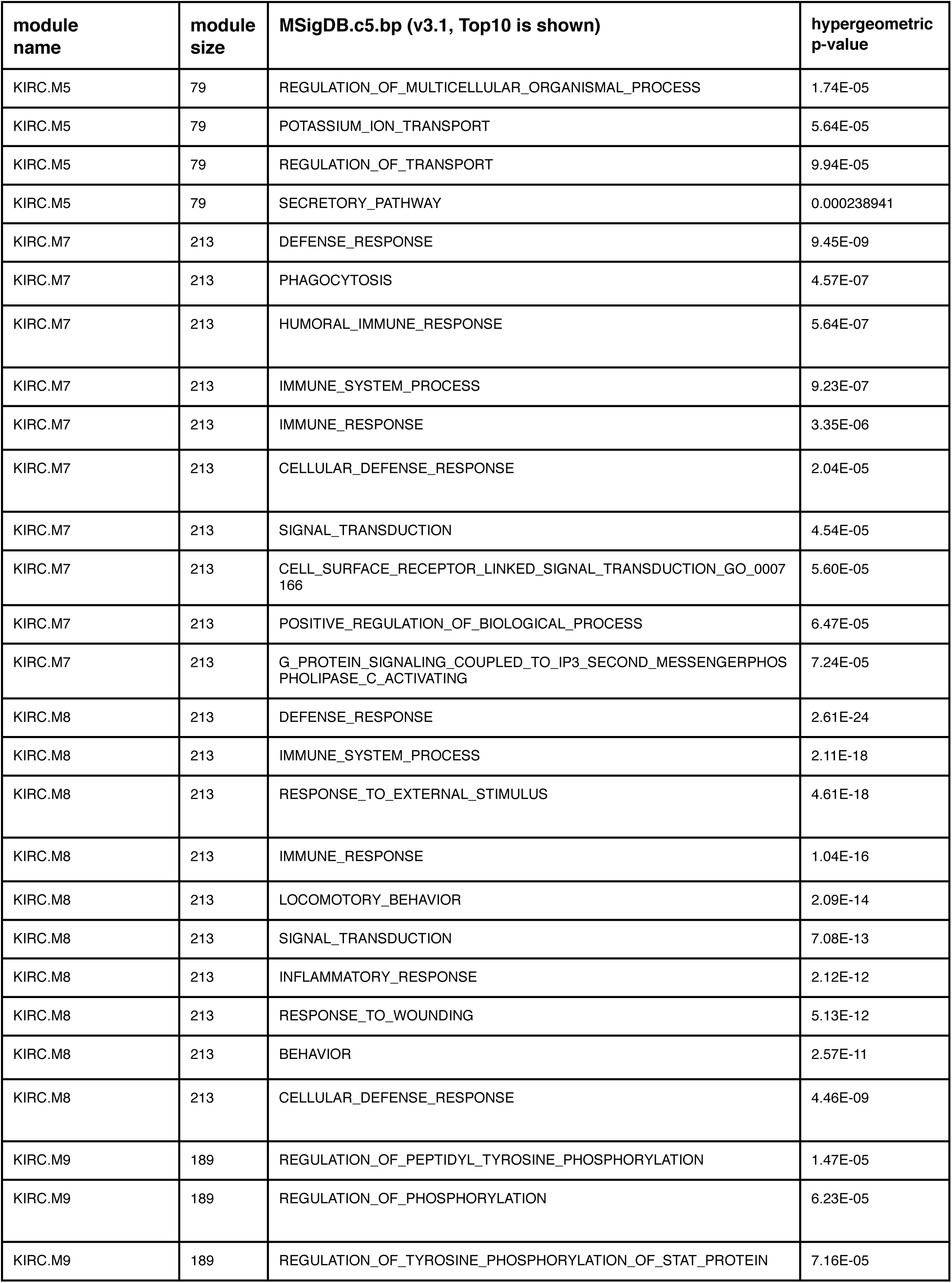
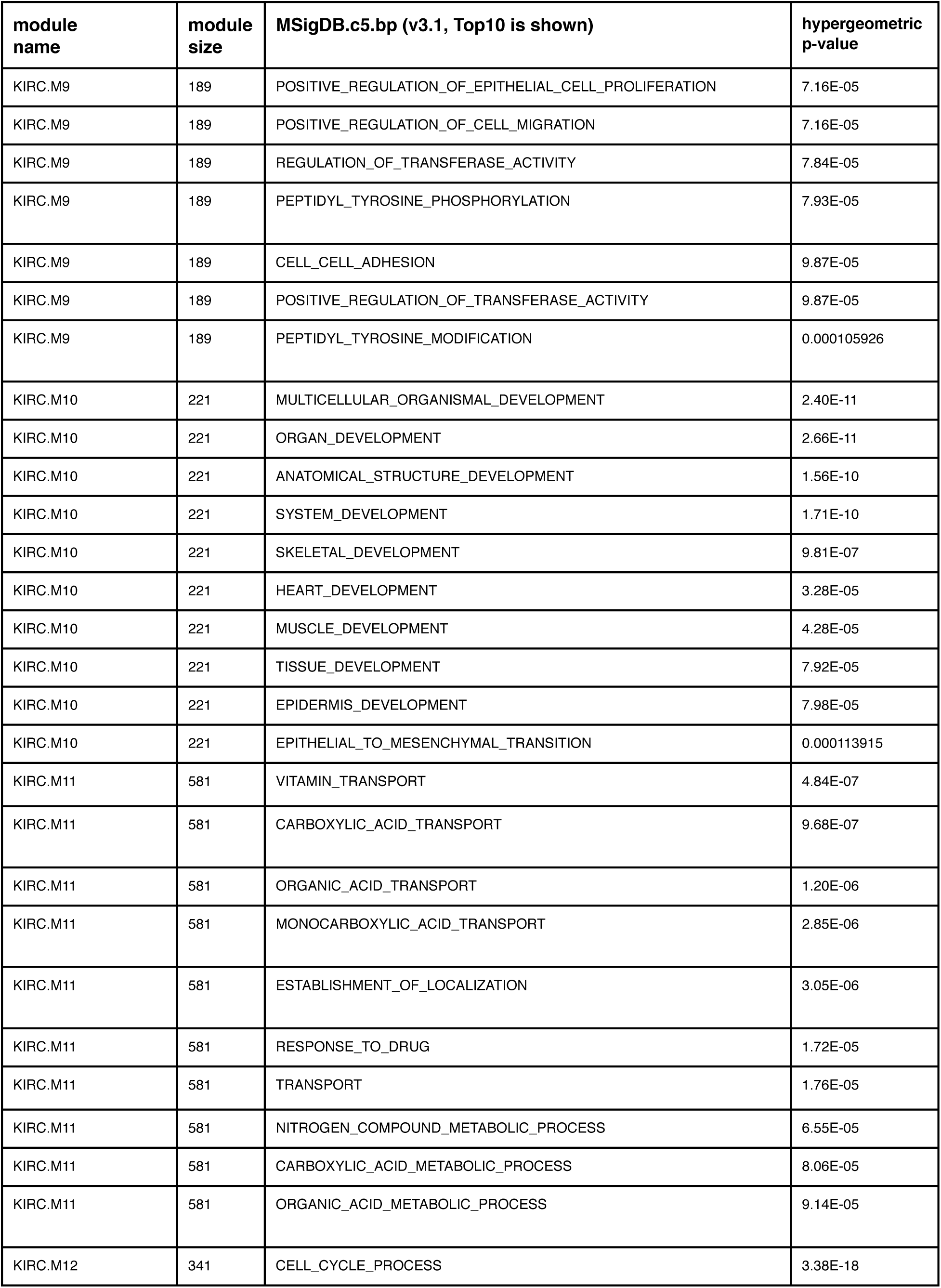
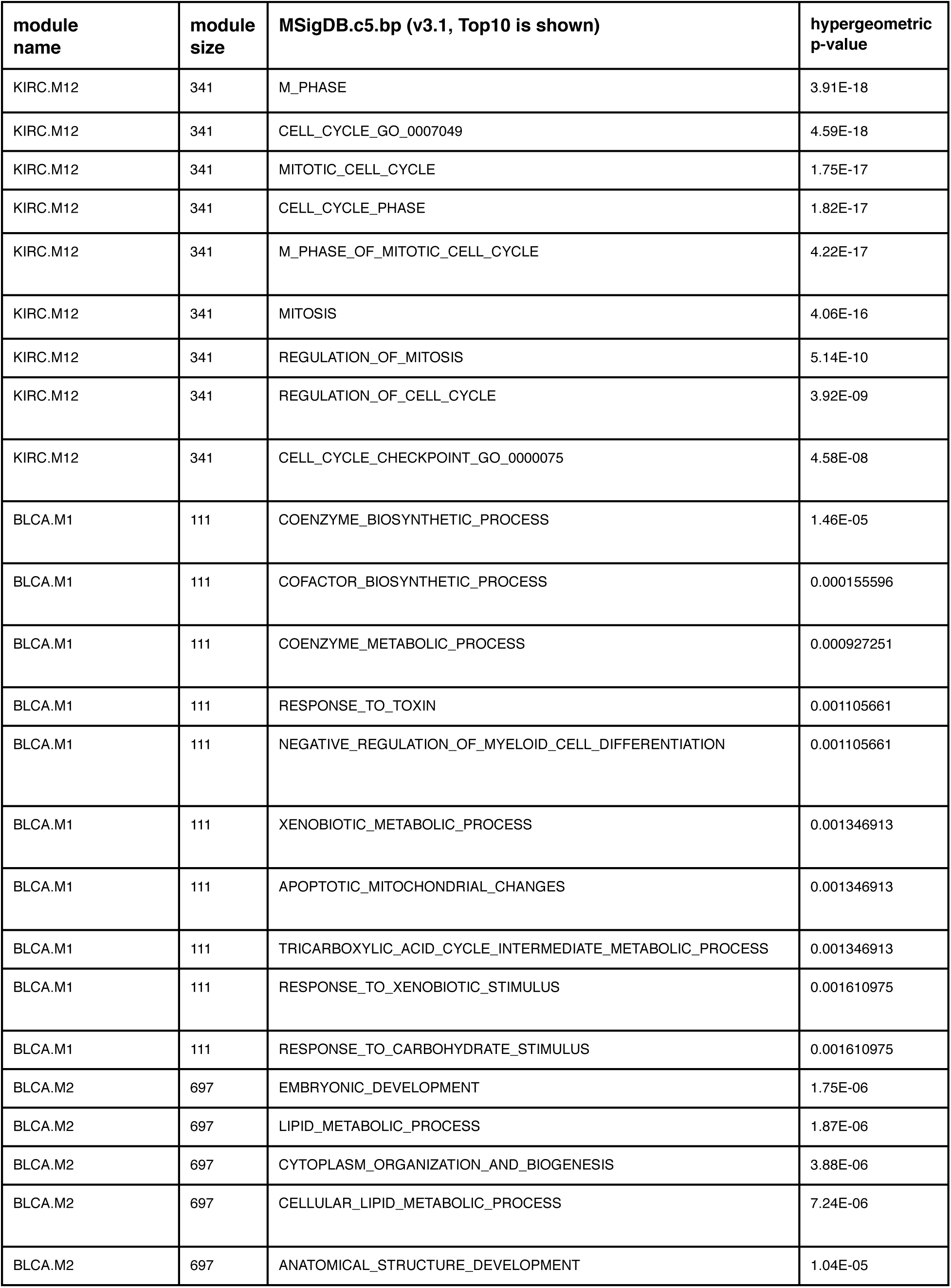
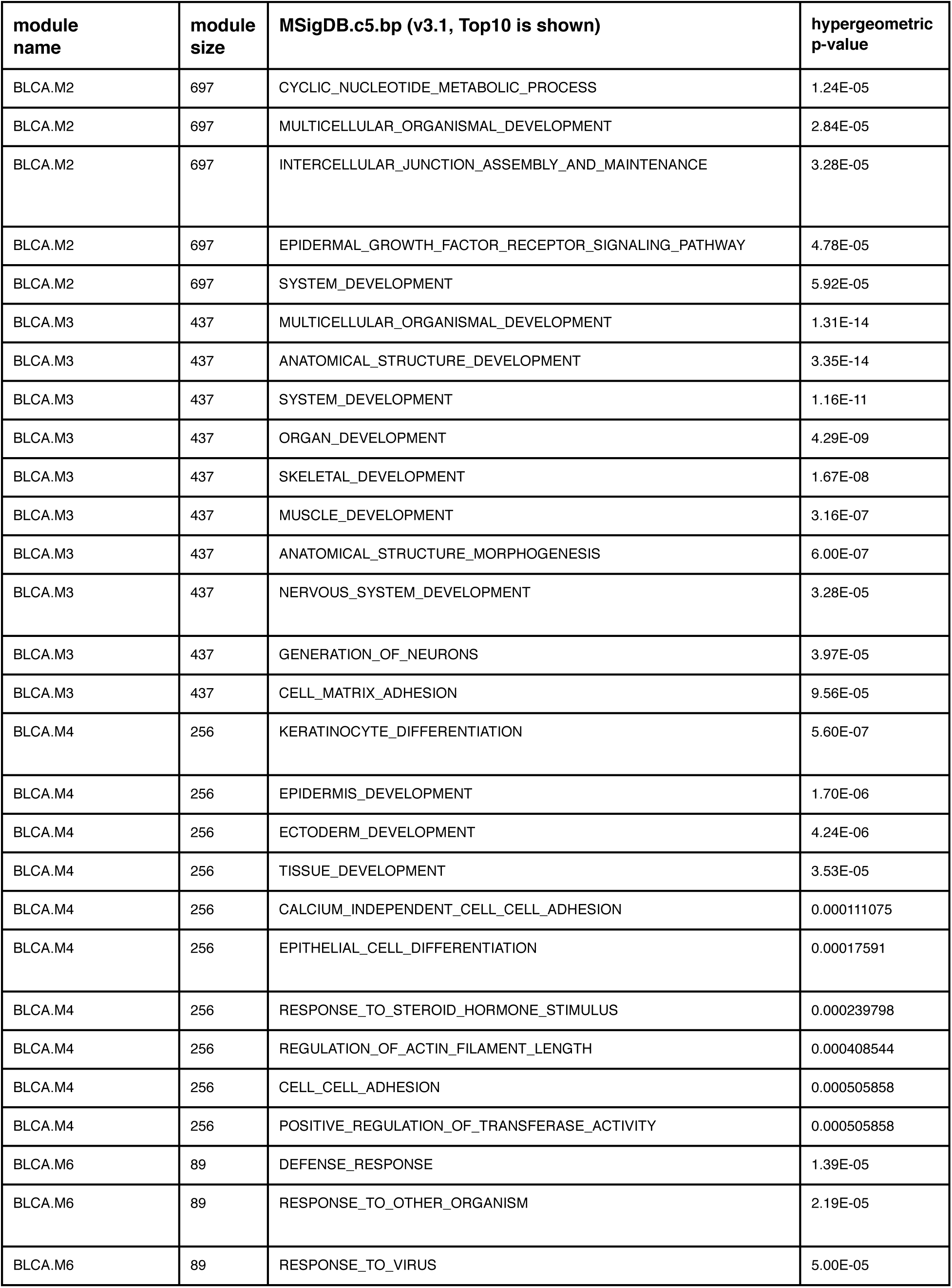
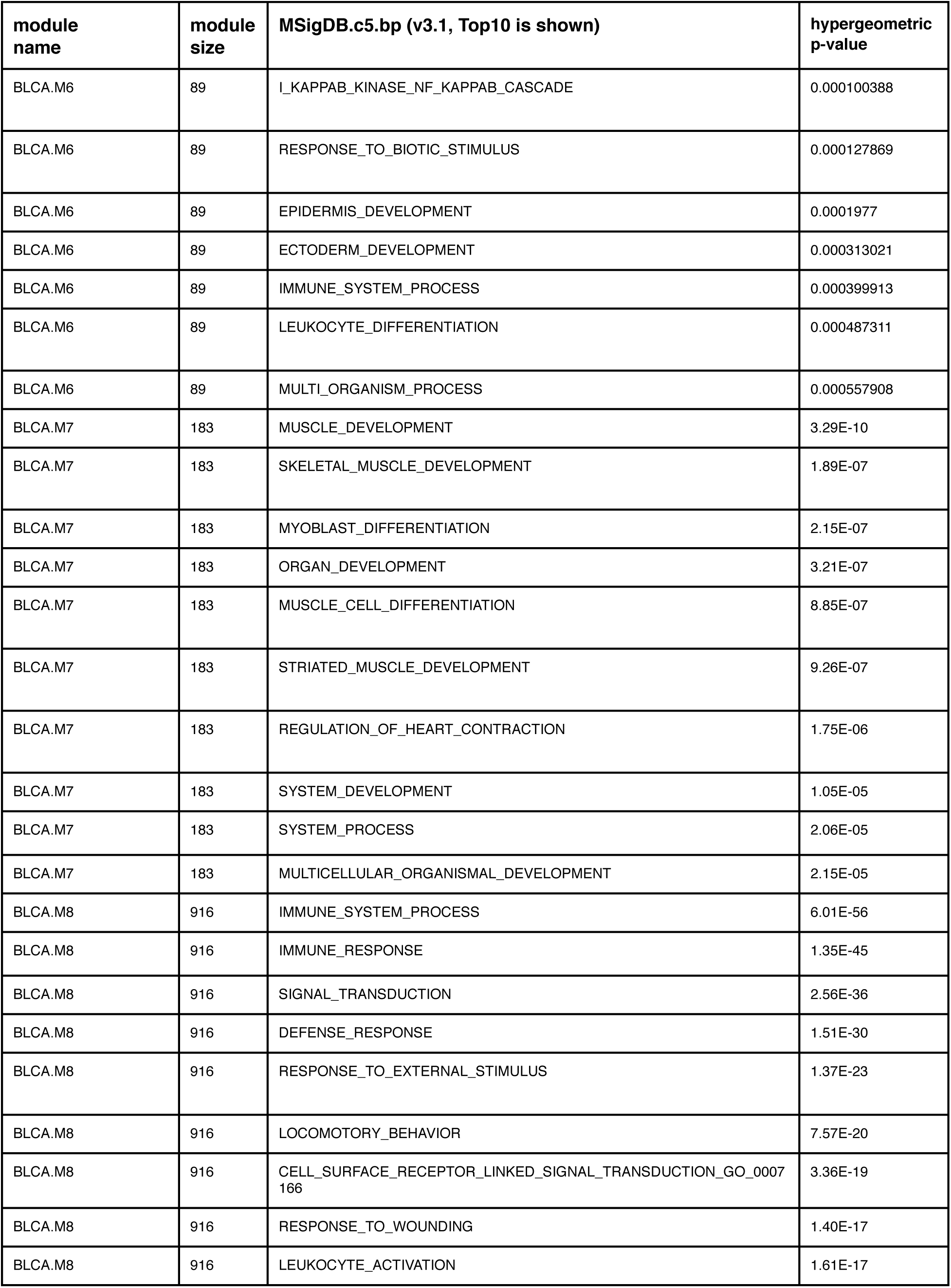
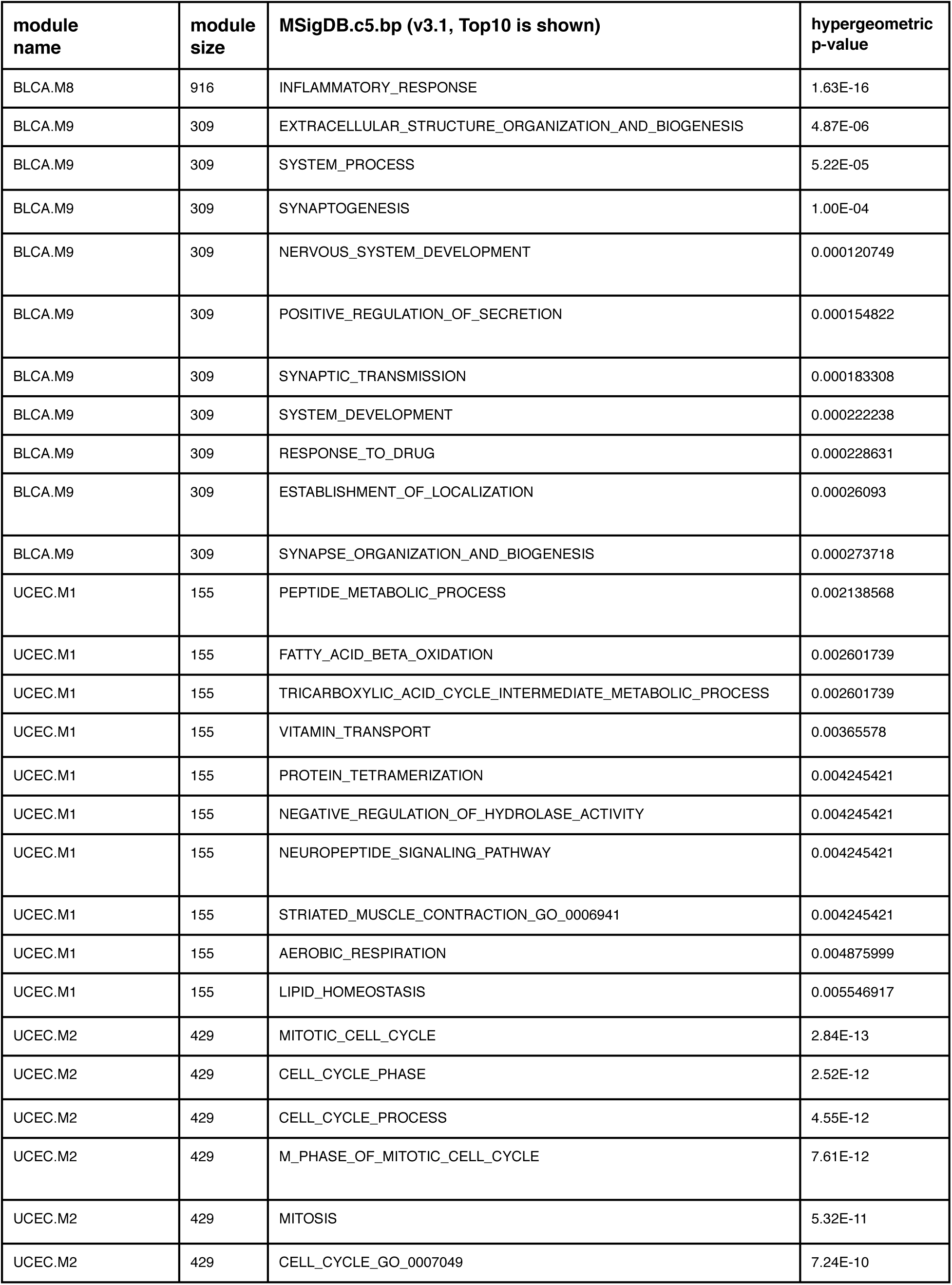
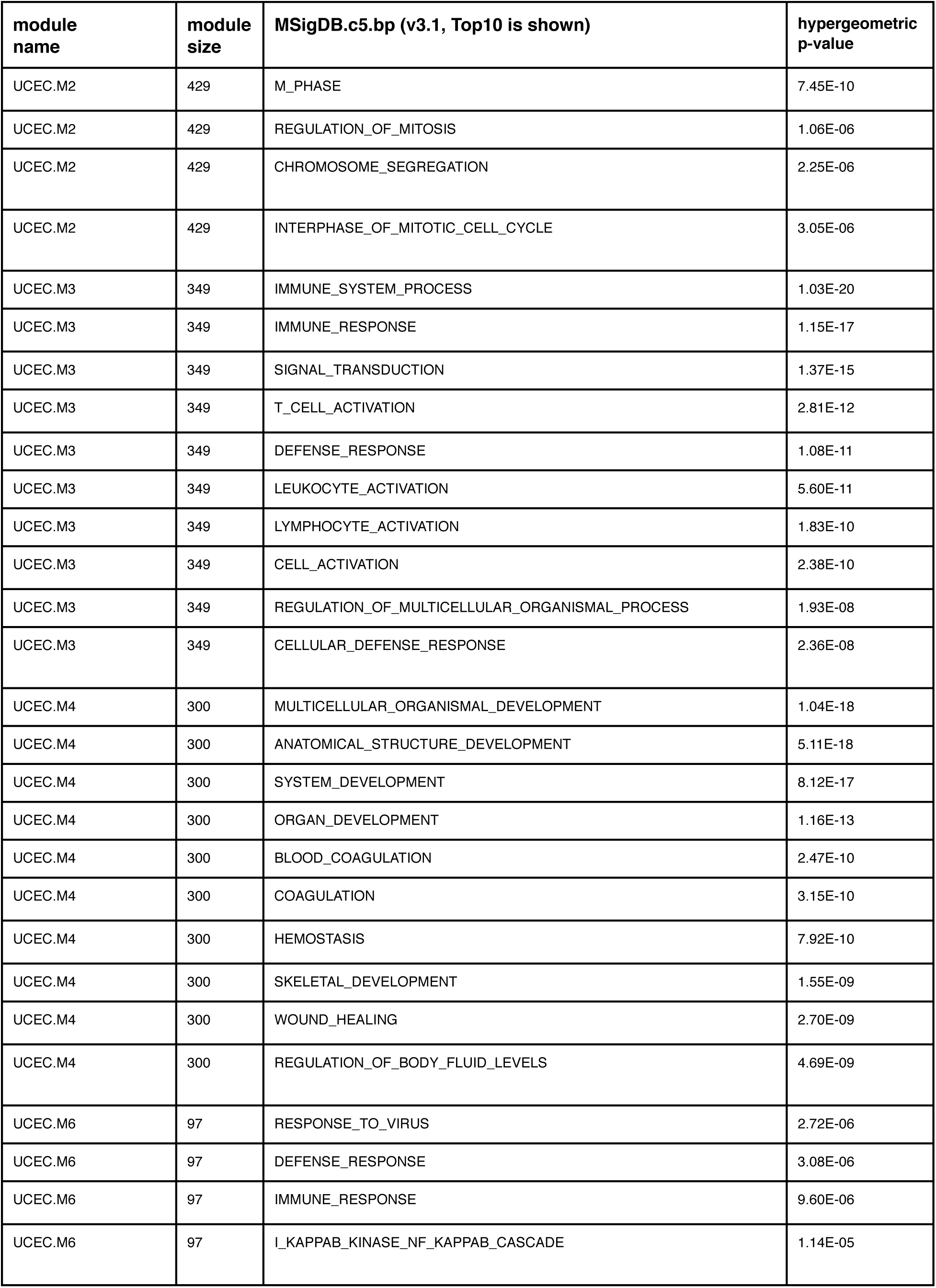
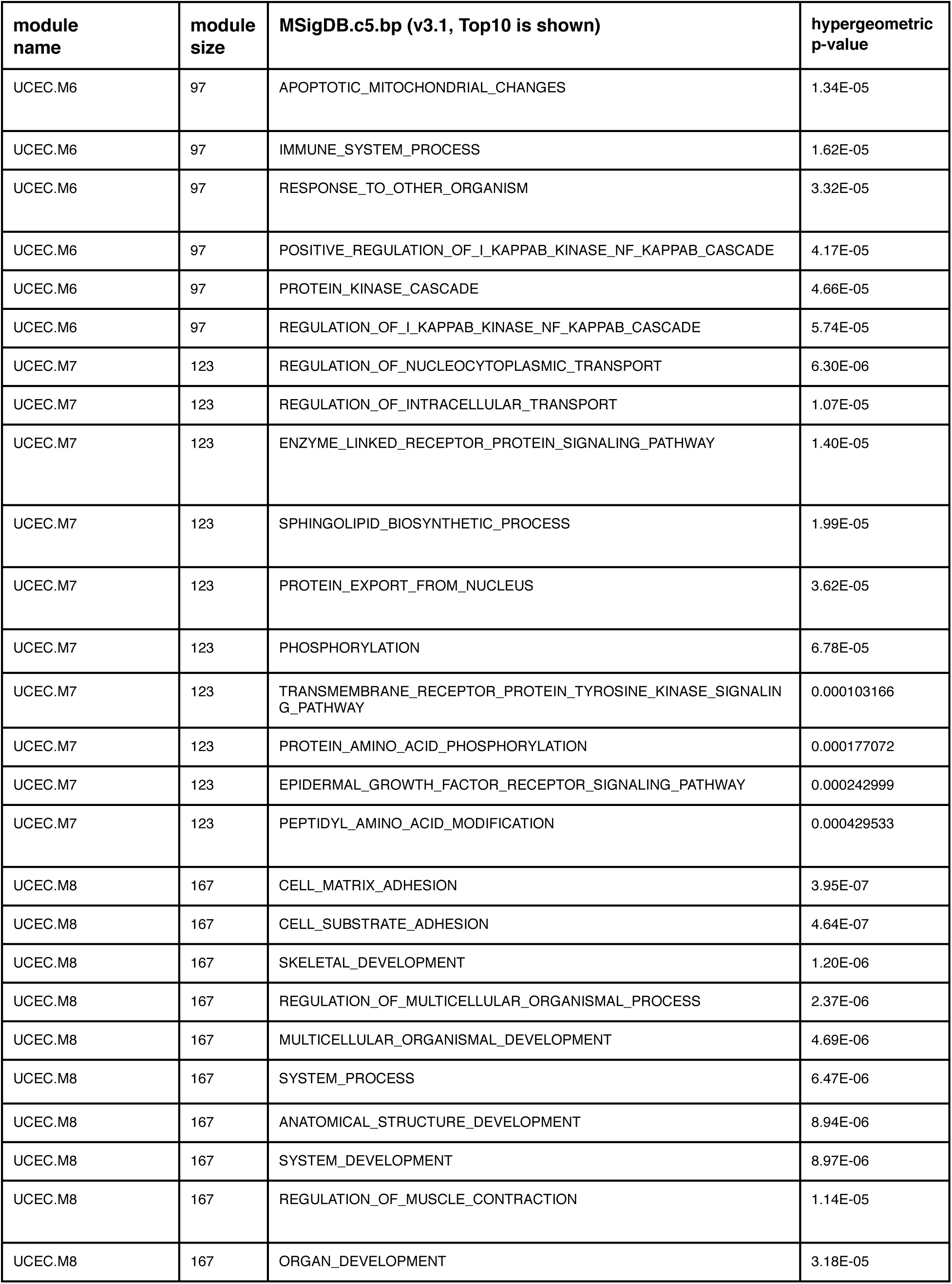
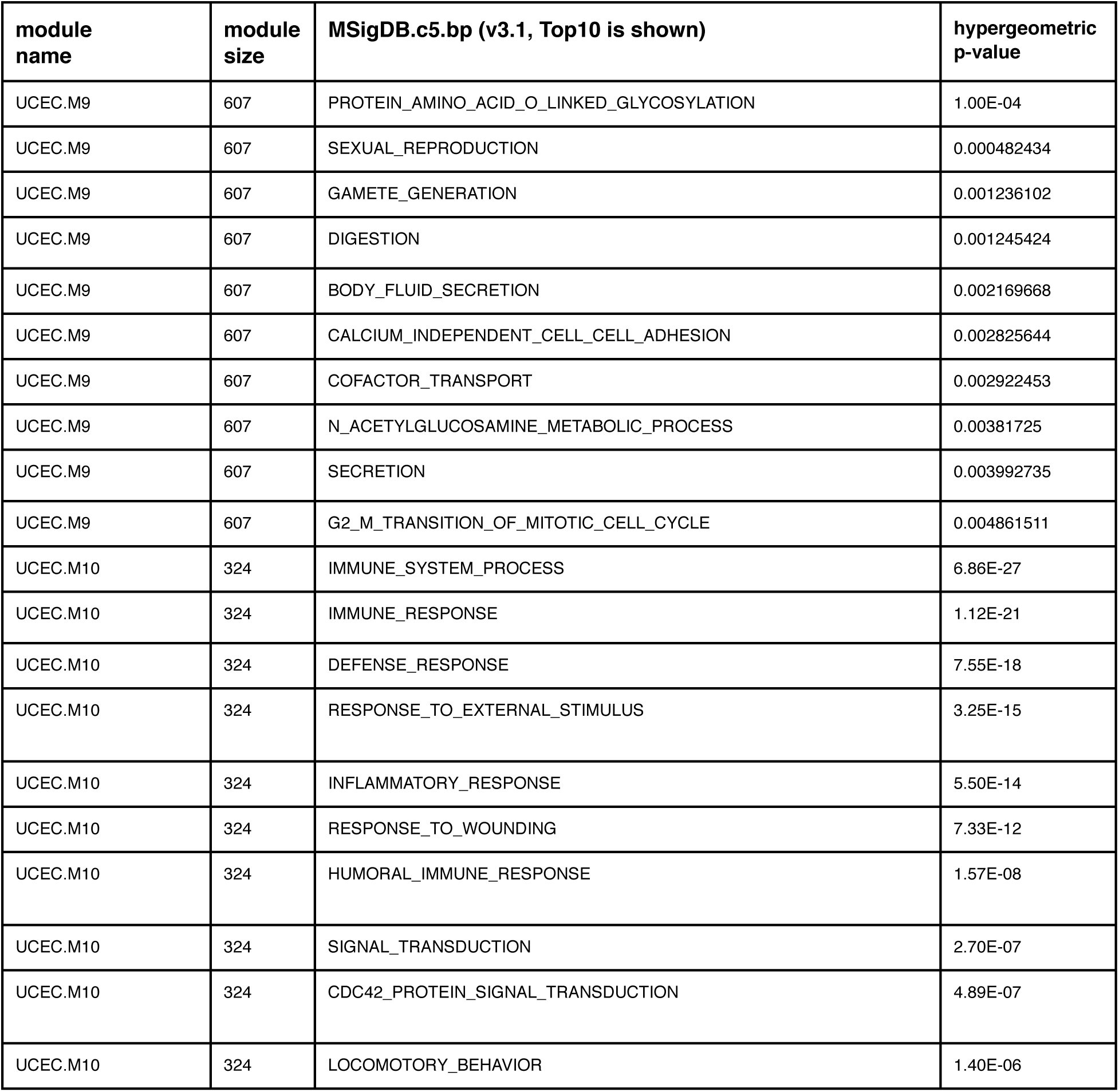

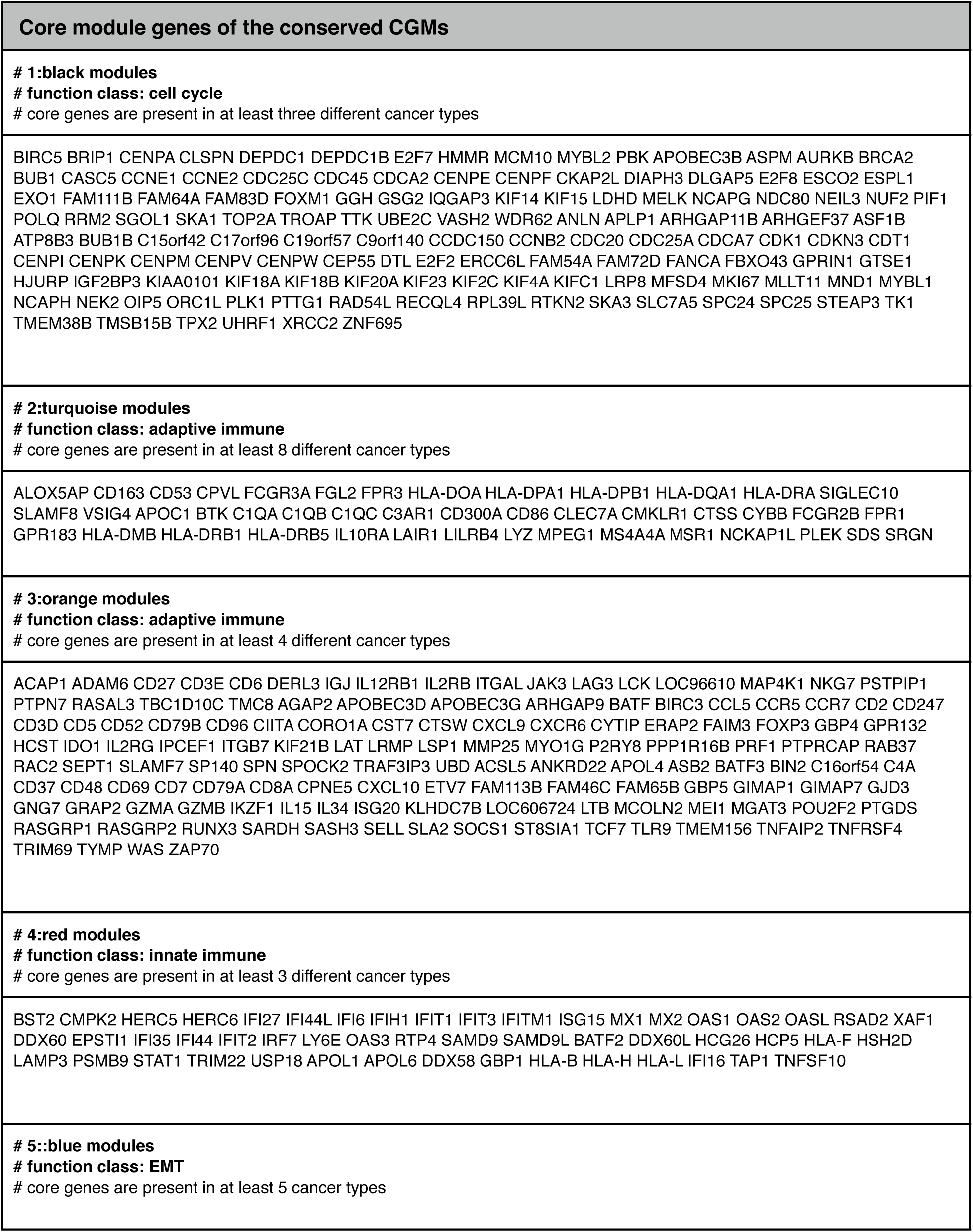
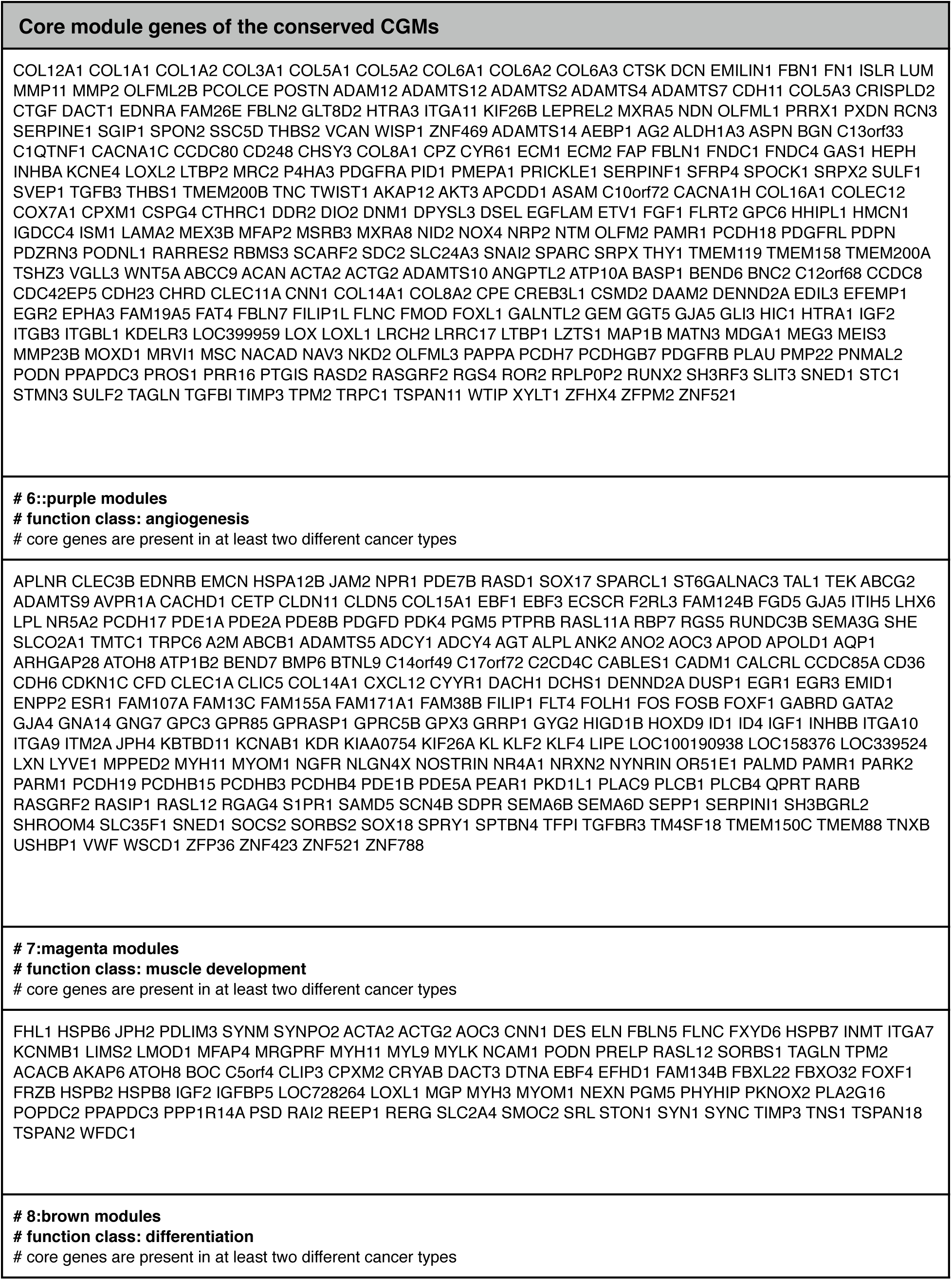
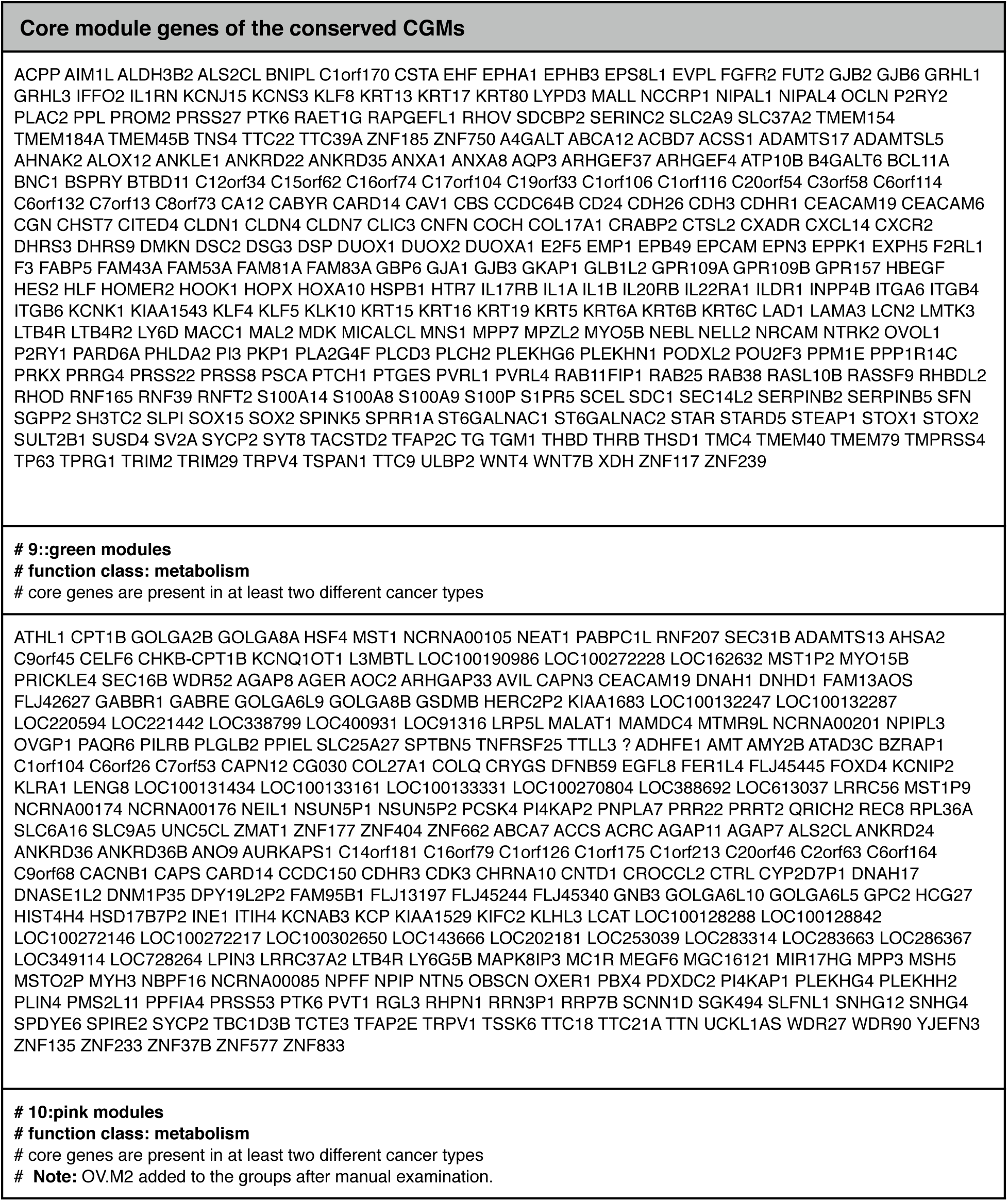
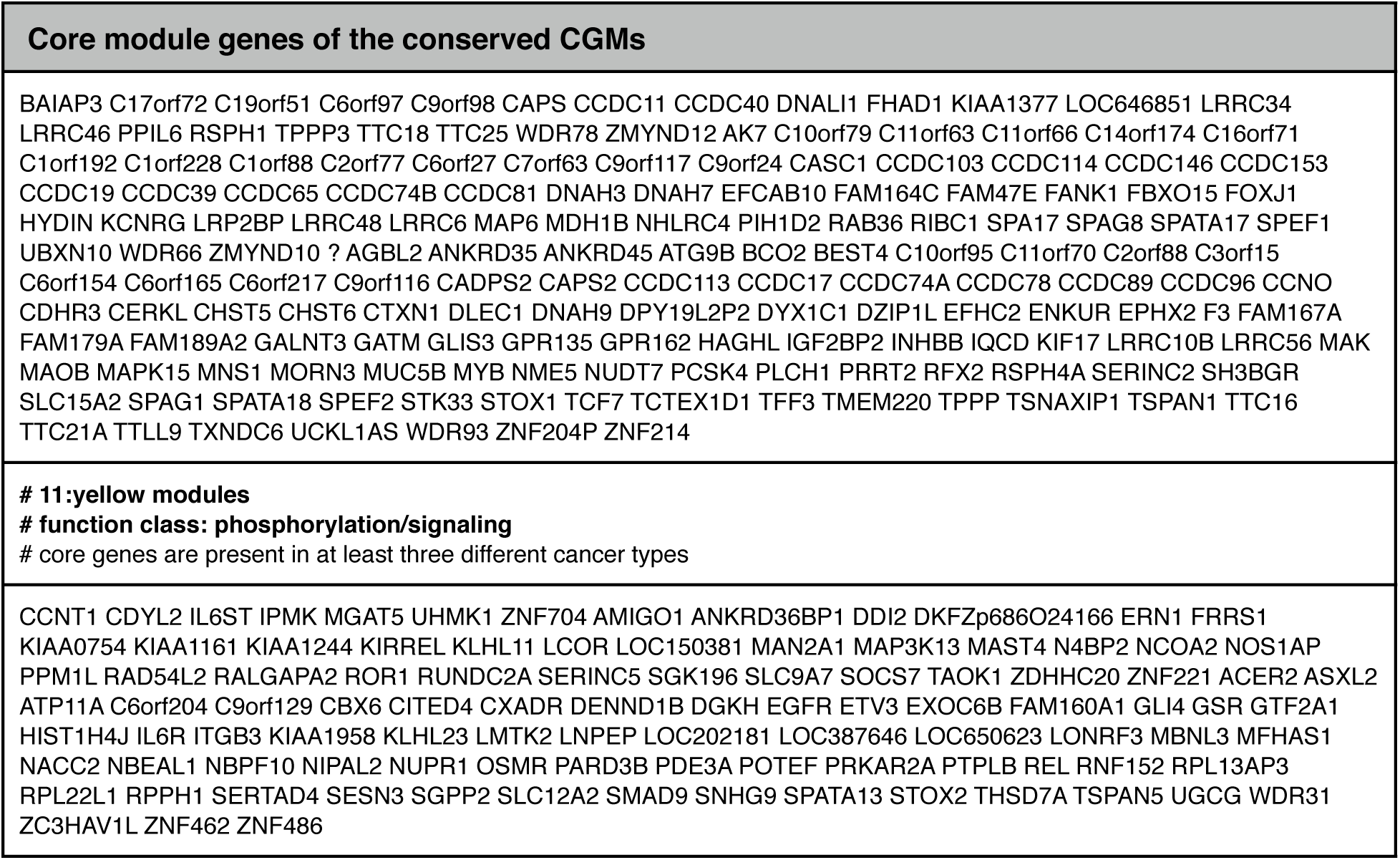

**Zou\_Suppl\_Table\_5**
|  | predicted module drivers |
| --- | --- |
| overlap with significantly mutated genes (MutSig) | TP53 BRCA1 BUB1B CDKN1A RB1 PTPN11 PIK3R1 JAK2<br>PIK3CA EGFR CTNNB1 RAC1 RASA1 AKT1 HRAS EP300<br>B2M HLA-A CCND1 SMAD4 CDH1 SMAD2 NCOA3 DMD<br>MYH9 ERBB2 STAG2 FGFR3 ERCC2 |
| overlap with cancer consensus genes | TP53 BRCA1 BUB1B XPO1 MYC RB1 CD74 LCK<br>PTPN11 CBL JAK1 ITK CD79B CD79A PIK3R1 JAK2<br>IL2 PIK3CA JUN EGFR SYK CTNNB1 AKT1 HRAS<br>EP300 CREBBP CYLD CCND1 ELN KDR TCF3 CDH1<br>NCOA1 PDGFRB MYH9 ERBB2 MET FGFR3 JAK3 ERCC2<br>ERCC3 |

